# Volumetric Cyclic Immunofluorescence for 3D Spatial Profiling of Tumour, Stroma, and Immune Structures in Human FFPE Tissue

**DOI:** 10.64898/2026.05.17.725158

**Authors:** Alex Y. H. Wong, Yi Daniel Lu, Ziyuan Zhao, Seo Woo Choi, Hojeong Park, Felix Zhou, Zoltan Maliga, Yvonne N. A. Anang, Soheil R. Talemi, Shannon Coy, Gaudenz Danuser, Sandro Santagata, Clarence Yapp, Peter K. Sorger

## Abstract

Mapping how parenchymal, immune, and stromal cells interact with extended structures such as blood vessels and nerves is challenging using conventional thin-section tissue imaging. Volumetric methods such as light-sheet fluorescence microscopy (LSFM) are ideal in these cases and LSFM is widely used in animal models such as fish and mice. This is enabled by well-established clearing methods, genetically encoded reporters, and vascular dyes. However, LSFM is rarely utilized for human tissue, which is primarily available as formaldehyde-fixed paraffin embedded (FFPE) specimens. Here, we present an approach to volumetric cyclic immunofluorescence (v-CyCIF) that enables 40-plex or greater imaging of immune and other cell types in human specimens up to 1 mm thick. We demonstrate the use of v-CyCIF to study neuroimmune interactions and lymphoid structures in normal and cancer tissues. Following LSFM imaging, specimens can be re-embedded and sectioned to ∼50 µm thick for high-resolution confocal imaging of punctate structures and cell-cell junctions. v-CyCIF therefore provides a flexible framework for multi-scale 3D profiling of clinical specimens across imaging formats and resolutions.

## INTRODUCTION

Functional tissue units (FTUs)^1,2^ such as colonic crypts, lymphoid tissue and breast lobules commonly span hundreds of microns to several millimetres in three dimensions. They include blood and lymphatic vessels, nerves, and other extended structures whose connectivity is difficult to discern in conventional 2D tissue sections. Immune cell types are also abundant in barrier tissues and are often found in dense organized assemblies that contain nerves and vessels. For example, gut-associated lymphoid tissue (GALT) normally includes Solitary Intestinal Lymphoid Tissue (SILTs)^3,4^ and, when chronic inflammatory signals are present (e.g. due to tumour infiltration) Tertiary Lymphoid Structures (TLSs) are also present. These immune aggregates contain 10^4^ −10^6^ or more T and B cells^5^ and play important roles in both normal tissue homeostasis and in diverse diseases, including solid cancers and inflammatory conditions^6,7^. Multiple collaborative programmes including the Human BioMolecular Atlas Program^8^ (HuBMAP) and the Human Tumor Atlas Network^9^ (HTAN), aim to build 3D spatial atlases of healthy and diseased FTUs. These efforts recognize that 3D imaging is essential for accurate immune profiling and morphological analysis^10^, but high-plex 3D datasets of human tissue remain rare and those that do exist are largely reconstructed from serial 5 μm thick sections^11–13^, supplemented more recently by high-resolution confocal imaging of sections up to ∼50 μm thick^14^.

Volumetric imaging of cleared tissues using methods such as light-sheet fluorescence microscopy (LSFM) is an ideal means of studying FTUs in intact tissue: it enables imaging of specimens 200-500 fold thicker than conventional histological methods (typically 4 to 5 µm thick) at single-cell resolution with minimal photodamage. LSFM of tissues is routine in model organisms, particularly in mouse neurobiology,^15,16^ and there has been substantial recent interest in extending LSFM to human specimens^17–21^. However, whereas tissues from model organisms can be fixed under conditions optimized for clearing, most clinical specimens are available only as formalin-fixed paraffin-embedded (FFPE) tissue. Furthermore, whereas structures of interest can be labelled in animal models using genetically encoded fluorescent reporters and perimortem perfusion of dyes^22^, such approaches are not applicable to human tissues. This makes labelling of specific proteins and other biomolecules entirely dependent on antibodies, other affinity reagents, and dyes. Prior to staining, FFPE specimens must also be dewaxed, rehydrated, and subjected to antigen retrieval to reverse formaldehyde cross-links; procedures not generally required for whole-mount animal tissues^23^. A further limitation for immune profiling is that many of the Cluster of Differentiation (CD) markers used to define immune cell types are plasma membrane proteins that are poorly preserved by standard clearing methods, most of which are strongly delipidating. Finally, LSFM is typically restricted to 3-4 fluorescent channels, compared to the 20–100 channels achievable with 2D multiplexed immunofluorescence, limiting the ability to dissect complex cellular ecosystems. These factors have limited the use of LSFM for the study of human tissue in health and disease.

In this paper, we describe volumetric cyclic immunofluorescence (v-CyCIF), an approach to high-plex LSFM and confocal microscopy optimized for analysis of FFPE human tissue. v-CyCIF combines membrane-preserving, delipidation- and dehydration-free clearing protocols; rapid antibody labelling via stochastic electrotransport; sample holders that support iterative imaging; and methods for re-embedding specimens following volumetric imaging so that they can undergo subsequent 2D or high-resolution 3D analysis. We have demonstrated 40-plex (16 cycle) v-CyCIF involving a dozen immune cell (CD) markers without significant tissue degradation, suggesting that higher-plex images are achievable. v-CyCIF has also been implemented across multiple microscopes using complementary protocols to create a flexible toolbox for studying human biology. This toolbox includes “virtual H&E” (v-H&E)^19,24–26^ which uses nuclear staining and autofluorescence to rapidly visualize tissue architecture prior to antibody-based imaging without the cost of antibodies while enabling analysis with machine learning (ML) computational pathology foundation models^26^. We apply v-CyCIF to multiple human tissue types, with a focus on secondary and tertiary lymphoid structures in the normal colon and in colorectal cancer (CRC). Germinal centres in these structures play a critical role in immunosurveillance of gut microbiota, tumours and response to immune checkpoint inhibitors^7^.

## RESULTS

### Membrane-preserving clearing for immune-rich FFPE tissue

In thick tissues, light scattering caused by lipids and extracellular matrix (ECM) reduces image contrast and must be mitigated by tissue clearing. Many clearing methods extensively delipidate tissue to improve transparency^27–32^, but we found that the resulting disruption of plasma membranes is broadly incompatible with antibody-based staining of membrane proteins, including the CD markers required for immune cell subtyping. In our hands, hydrogel-based methods optimized for unembedded paraformaldehyde (PFA)-fixed tissue, such as CLARITY^30^ (without modification) also resulted in poor immunostaining with FFPE specimens. Methods that omit hydrogels but combine delipidation with detergents, such as OPTIClear, preserved staining for structural and nuclear proteins (e.g., Ki67; E-cadherin, ECAD; α-smooth muscle actin [αSMA]) as previously reported^33^ but performed poorly with many CD antigens (DS1; **Extended Data Fig. 1**).

We therefore developed a delipidation-free v-CyCIF clearing protocol compatible with standard heat-induced epitope retrieval (HIER^34^); epitope retrieval is required to rehydrate FFPE tissue and reverse formaldehyde cross-links prior to immunostaining (**Supplementary Fig. 1a,b)**. Our clearing approach uses a one-step refractive index matching solution (RI=1.52) containing a modified formulation of four previously described clearing reagents: 2,2′-Thiodiethanol (TDE)^35^, Iohexol, D-sorbitol, and N-Methyl-D-glucamine, at a pH of 7-8 (see **Supplementary Note** for further discussion of tissue clearing). Staining is performed following epitope retrieval for 2-3 hr at room temperature or 12-16 hr at 4°C and clearing was found to be effective with agarose-stabilized sections of multiple human tissue types (**Supplementary Fig. 2**). The eleven specimens analysed and described in this paper gave rise to 12 datasets overall (DS1-12; **Supplementary Table 1**).

### Multiplex immunostaining of millimetre-thick human FFPE tissue

Tissues cleared using the v-CyCIF protocol were stained with dyes and antibodies via active or passive diffusion and imaged on one of several different confocal and light sheet microscopes (**Fig. 1a-e**; **Supplementary Note**). By staining tissues with DAPI and imaging nuclei together with autofluorescence, we could generate virtual haematoxylin and eosin (vH&E) images (**Fig. 1a**). DAPI-staining also made it possible to evaluate the quality of clearing using the Fourier ring correlation quality estimation score (FRC-QE)^36^. FRC-QE data demonstrated near-uniform clearing of most tissues tested up to ∼1 mm thick (**Fig. 1f, g**). Although this depth is less than that achievable with strongly delipidating approaches (**Supplementary Table 2)**, it is well suited to imaging many epithelial and immune-rich human tissues while preserving membrane antigens required for immune profiling.

**Figure 1:**
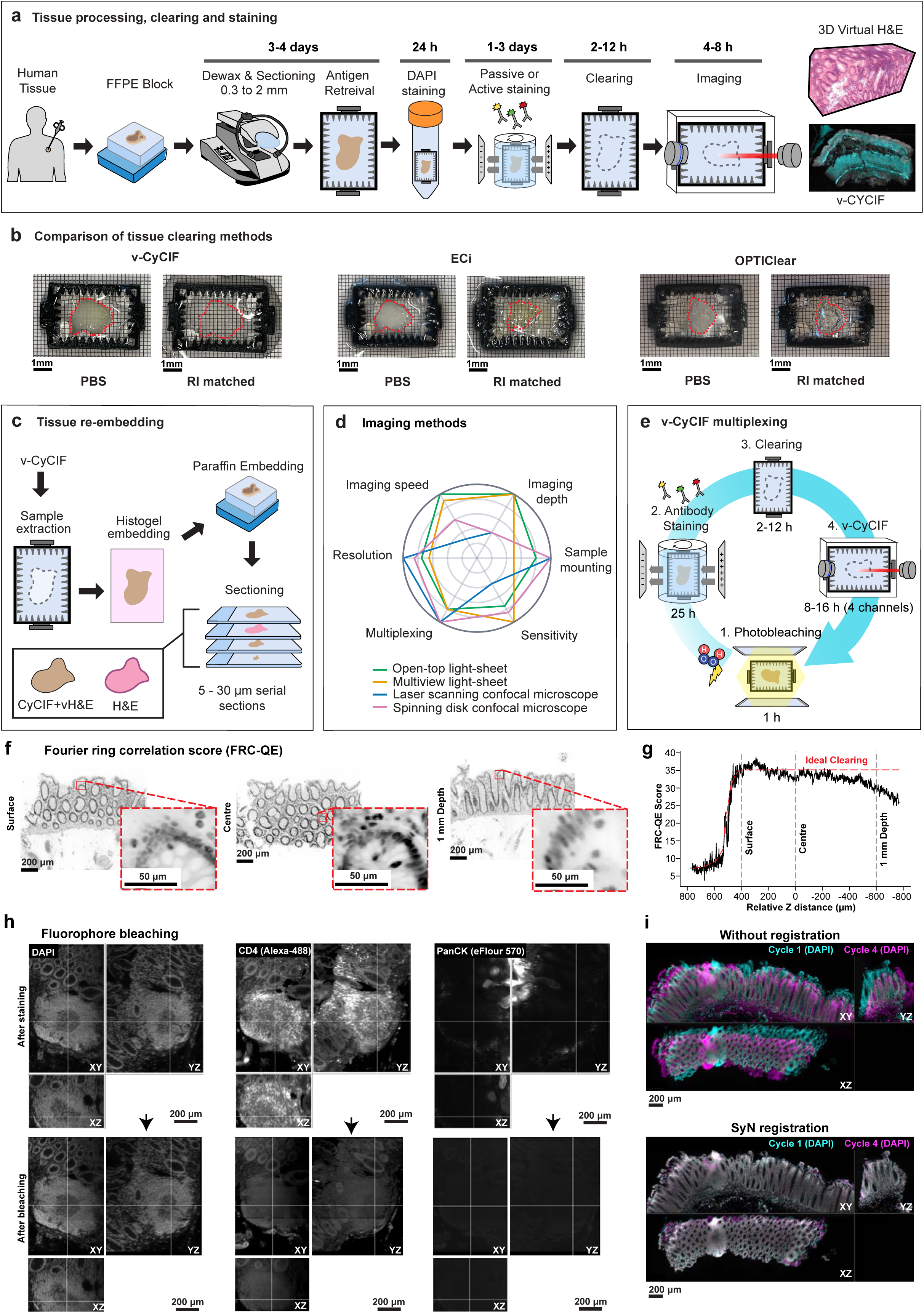
Overview and validation of the v-CyCIF pipeline for multiplexed 3D tissue imaging. **a,** Workflows for v-CyCIF. **b,** Representative photographs of a cleared human colon sample imaged in PBS (left) or refractive-index (RI)-matched imaging buffer (right) following v-CyCIF, ECi, and OPTIClear clearing solutions. Red outlines indicate the sample boundary. Scale bars, as indicated. **c,** Re-embedding of tissues into paraffin blocks using HistoGel for conventional 2D CyCIF on tissue sections or high-resolution 3D CyCIF. **d,** Radar-plot comparison of Open-top light-sheet (OTLS), Multiview selective-plane illumination microscope (MuViSPIM), Laser-scanning confocal microscopy (LSCM) and Spinning-disk confocal microscopy (SDCM) across key attributes (resolution, multiplexing, sensitivity, sample mounting, imaging depth, and imaging speed). **e,** v-CyCIF multiplexing includes iterative photobleaching, staining, clearing and 4 channel v-CyCIF imaging. **f,** Representative single optical plane images sampled near the tissue surface, relative centre, and 1mm depth. Scale bar: 100 μm. **g,** FRC-QE score plotted as a function of relative Z distance to the tissue centre. Dashed vertical lines mark the Z-positions corresponding to the surface, relative centre, and 1mm depth respectively. **h,** Example of v-CyCIF demonstrating fluorescent signal (DAPI, CD4, PanCK) removal by photobleaching: after staining (top) and the same field after photobleaching (bottom), shown as orthogonal views. Scale bars: 200 μm. **i,** Mounting accuracy between cycle 1 and 4 with and without registration using rigid and affine transformation with symmetric normalization (SyN) within ANTs.

Cyclic IF imaging requires efficient inactivation of fluorophores with preservation of antigenicity across imaging cycles. We accomplished this using base-catalysed oxidation (bleaching) of fluorophores following each imaging cycle with alkaline hydrogen peroxide solution and light^37^. We found this approach to be superior in most cases to chemical stripping of antibodies, which leads to protein loss and damages tissue^38^. The completeness of fluorescence signal extinction was confirmed for each tissue type and antibody conjugate by imaging before and after bleaching (**Fig. 1h** and **Supplementary Fig. 3**). Consistency in the orientation of samples across cycles of staining, clearing, imaging, and bleaching was maintained by embedding tissues in agarose and mounting them in detachable 3D-printed sample holders (**Extended Data Fig. 2; Supplementary note 1**).

Multi-cycle images were registered using the DNA channel (**Fig. 1i**), with the first cycle serving as a reference for subsequent cycles. Coarse global alignment was achieved using rigid and affine transformations followed by non-linear warping to correct local deformations, with transformations propagated to all channels within each cycle. High-plex v-CyCIF of large specimens generated multi-terabyte volumetric datasets (∼2.5 × 10^11^ voxels) and we found it difficult to perform accurate registration at full resolution. We therefore subdivided specimens into regions of ∼5 × 10^10^ voxels each and registered them using symmetric normalization (SyN) in ANTs^39^, a software package originally developed for MRI analysis. Single-cell data were generated by 3D segmentation using a combination of Cellpose^40,41^ and u-Segment3D^42^, which leverages a computationally efficient 2D-to-3D segmentation framework (**Supplementary Fig. 4**, see **Methods** for details).

The specificity of antibody-based staining was evaluated at single-cell resolution by comparing v-CyCIF images with standard CyCIF of the same specimen via re-embedding and sectioning as described below (**Fig. 2 a**-**d**). In the case of CD8, for example, the comparison of 25 μm tissue sections with virtual slices from a 1 mm cleared volume confirmed a characteristic morphology corresponding to a membrane ring with a dimmer centre at the nucleus (red arrows; **Fig. 2e**). Intensity profiles computed along lines spanning single cells and multi-cell structures in virtual sections from v-CyCIF volumes were also used to evaluate the specificity of staining. For example, tissue stained with βIII-tubulin (TUBB3), a class III β-tubulin isoform and pan-neuronal marker^43^, generated high signal-to-background ratio (SBR) images of extended neuronal structures (**Fig. 2e**). We also checked for consistency in staining patterns when antibodies are conjugated with different fluorophores (**Supplementary Fig. 5**). Having antibodies conjugated with a range of fluorophores can be a valuable approach during panel design. For any specific antibody it was necessary to evaluate penetration depth and staining quality empirically. The relevant parameters remain incompletely understood but include protein abundance, tissue type, antibody clone, and fluorophore. In some cases, poor staining of a marker could be improved by changing the clone or fluorophore. Passive staining over a ∼2-day period was generally effective with colon specimens up to 1 mm thick and ∼20 mm in X and Y, but for some antibodies and tissues, active staining by stochastic electrotransport (SE)^44,45^ was necessary. SE applies a rotational electric field to actively distribute dyes and antibodies throughout tissue and can be accomplished using SmartBatch instruments and reagents from LifeCanvas Technologies (**Fig. 2f & 1h**). To date, we have tested ∼70 antibodies previously validated for 2D CyCIF^46^ for compatibility with v-CyCIF across multiple tissue types (DS2-3; **Extended Data Fig. 3 a-g**). Of these, 49 were compatible with passive staining and 21 performed well with stochastic electrotransport, particularly in thicker, dense specimens (**Supplementary Table 3**). Most antibodies were directly conjugated to fluorophores, although secondary antibodies could also be used (e.g., donkey anti-rabbit IgG against rabbit anti-Septin-2, a GTP-binding cytoskeletal protein^47^ as shown in **Extended Data Fig. 3e**). As in 2D CyCIF, each secondary antibody species (e.g., goat, rabbit, mouse) can only be used once per multi-cycle experiment.

**Figure 2:**
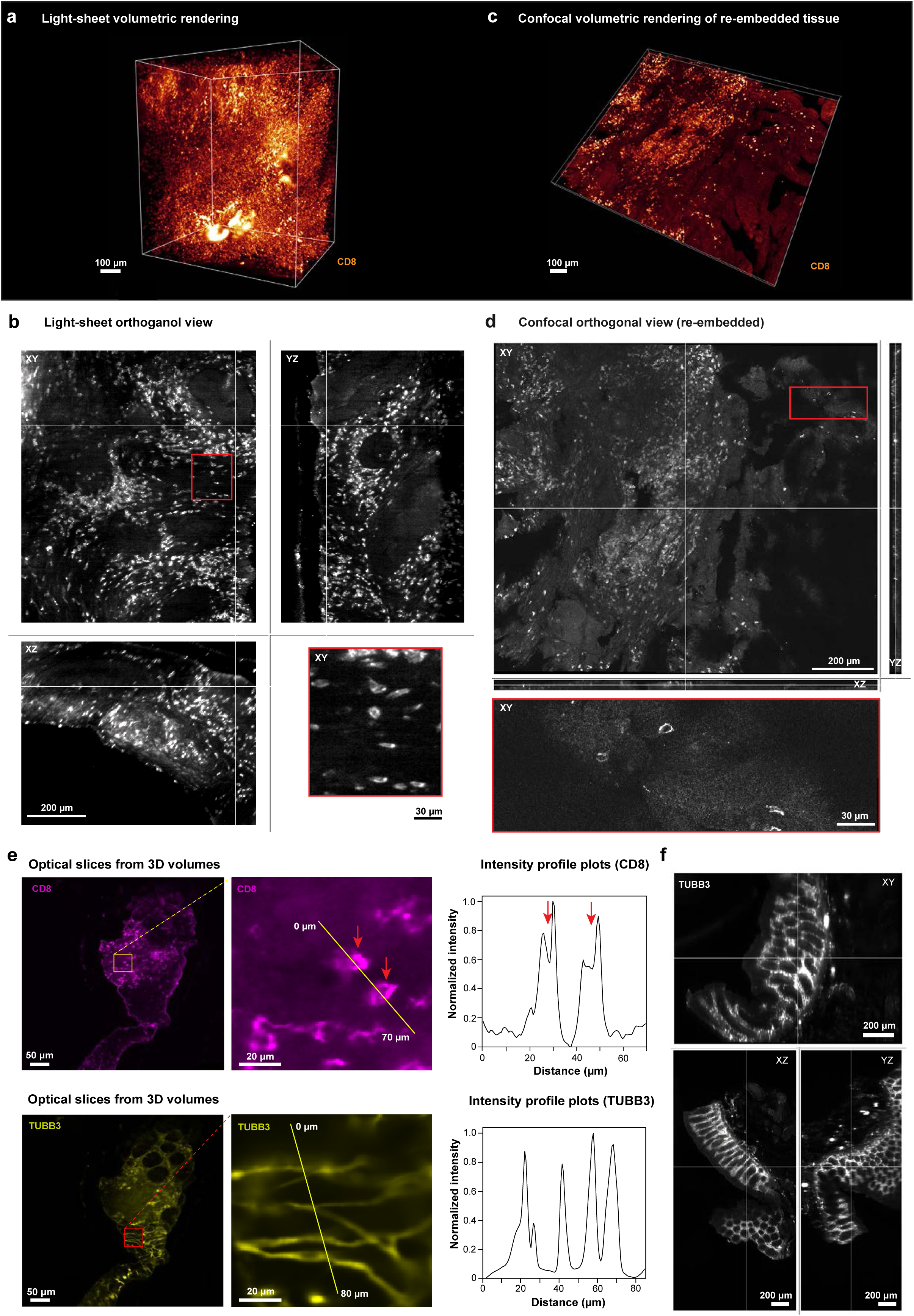
Comparison of CD8⁺ T cells at single-cell resolution in cleared and re-embedded tissues. **a,** 3D light-sheet volume rendering of CD8 signal (glow) in the sample. Scale bars: 100 μm. **b,** Orthogonal view of CD8 imaged by LSFM from panel **a**. Scale bars: 200 μm; Red box highlights diverse CD8+ T cell morphologies, Scale bars: 30 μm. **c,** Corresponding 3D view of the same specimen after re-embedding, sectioning and imaging by confocal microscopy (glow mode). Scale bars: 100 μm. **d,** Confocal orthogonal view of CD8 from a comparable region. Scale bars: 200 μm; Red ROI shows magnified view of CD8^+^ cells, Scale bars: 30 μm. **e,** Single-channel views (left) of optical slices with zoom in regions (right) for TUBB3 (yellow) and for CD8 (magenta) Scale bars: 50 µm. Yellow line indicates the transect used for corresponding intensity profile plots. Scale bars: 20 µm. **f,** Orthogonal views of TUBB3 staining with v-CyCIF clearing in 1 mm thick section. Scale bars: 200 µm.

### Exemplary v-CyCIF of tissue-resident immune structures

In the normal colon, SILTS develop postnatally^48,49^ to support local immune surveillance and adaptive immunity; their dysregulation contributes to chronic intestinal inflammation, inflammatory bowel disease, and susceptibility to infection.^50^ To study these immune structures we performed 40-plex LSFM-based v-CyCIF on the mucosal layer of a 1.4 mm × 8 mm × 1 mm (H×W×D) specimen of normal colon containing a single classical ovaloid SILT (Fig. 3a,b; **Extended Data Fig. 6; Supplementary Fig. 6,7;** DS4). Upon review, ∼40 antibodies were judged to have sufficient signal to noise for subsequent registration and analysis. Note that we commonly combine validated antibodies with ones being tested as a cost-effective means of extending antibody panels; only high-quality data (as described above; **Supplementary Table 5**) are retained in the final multiplexed image.

**Figure 3:**
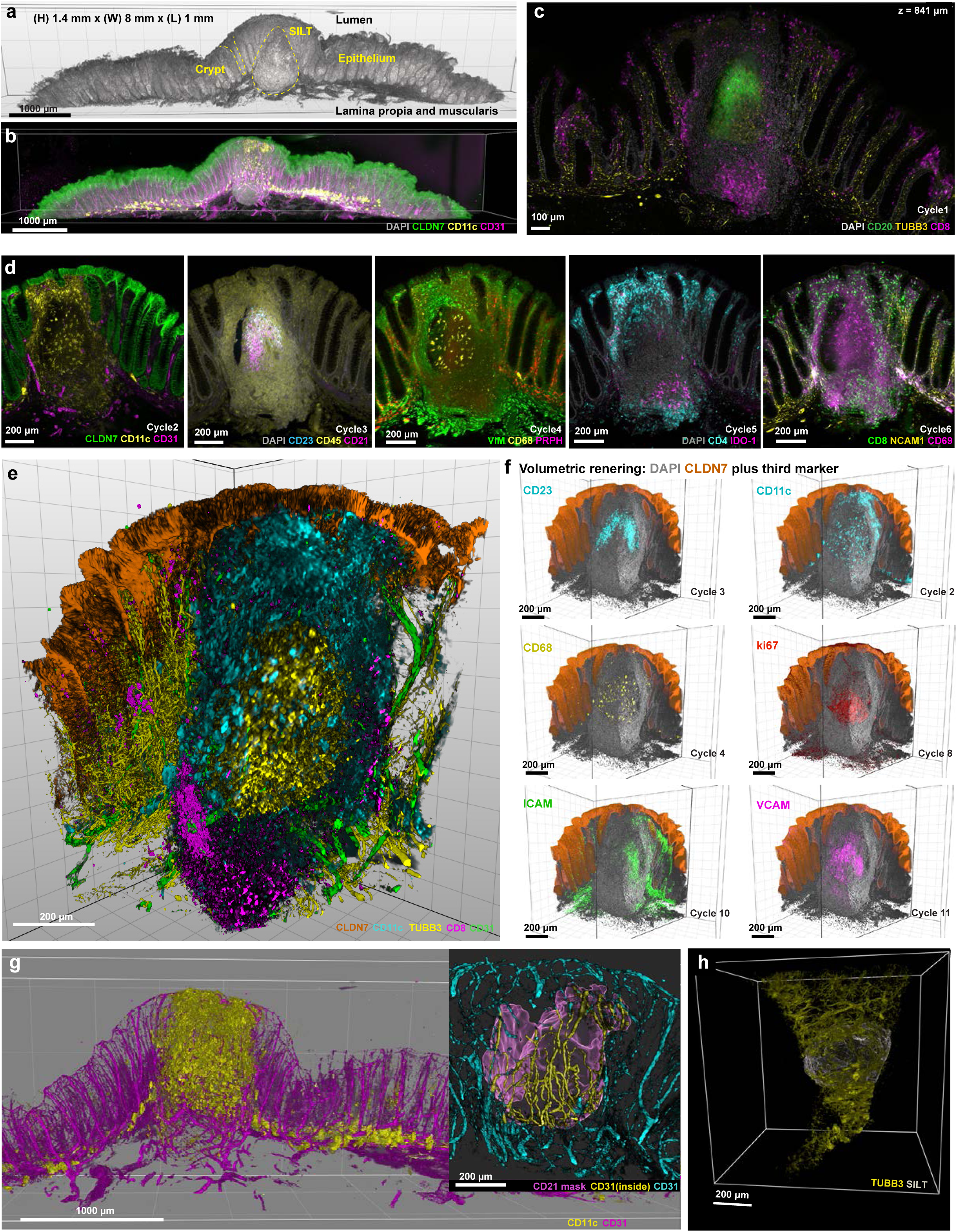
v-CyCIF imaging of a solitary intestinal lymphoid tissue in normal colon. **a,** 3D rendering of the cleared normal colon tissue imaged in its entirety (grey, DAPI) on DALISPIM light sheet microscope; volume dimensions 1.4 mm (H) × 8 mm (W) × 1 mm (L). **b,** Maximum intensity projection of cycle 2 (DAPI, grey; Claudin-7 (CLDN7); green, CD11c; yellow and CD31; magenta). **c,** Single optical section from Cycle 1 at z = 841 µm (DAPI, grey; CD20, green; TUBB3, yellow; CD8, magenta). Scale bar: 1000 µm **d,** Optical section through a germinal centre showing follicular dendritic cell and myeloid cell organization together with stromal markers across six imaging cycles. Cycle 2: CLDN7 (green), CD11c (yellow), CD31 (magenta). Cycle 3: DAPI (grey), CD23 (cyan), CD45 (yellow), CD21 (magenta). Cycle 4: VIM (green), CD68 (yellow), peripherin (PRPH, magenta). Cycle 5: DAPI (grey), CD4 (cyan), IDO-1 (magenta). Cycle 6: CD8 (green), NCAM1 (yellow), CD69 (magenta). Scale bar: 200 µm. **e,** 3D rendering of the SILT stained for CLDN7 (orange), TUBB3 (yellow), CD8 (magenta), CD11c (cyan), and CD31 (green). Scale bar: 200 µm. **f,** 3D rendering of 6 SILT associated markers acquired over 11 cycles from the same crop volume; the cycle of acquisition is indicated above each panel and the marker below. Epithelium (CLDN7, orange) and DAPI (grey) are shown in every panel as a spatial reference. Scale bar: 200 µm. **g,** 3D rendering (blend) of vascular architecture in the normal colon with CD11c dendritic cells, inset shows the external CD31 vessels (cyan) and CD31(yellow) within the CD21 mask (magenta) **h,** Rendering of SILT (DS5; grey) within the basket of nerves (Yellow; TUBB3). Scale bar 200 µm.

v-CyCIF imaging revealed a mature ovaloid SILT embedded in the mucosal epithelium with B cells polarized towards the lumen and coincident with a germinal centre (GC) defined by a network of CD21^+^ CD23^+^ follicular dendritic cell (FDCs; **Fig. 3a-d; Supplementary Video 1**). We observed high SBR staining for 8 CD antigens, markers of cell division such as Ki67, and multiple cell adhesion proteins (e.g. ICAM-1 and VCAM.). In the lamina propria, blood vessels were relatively straight but in SILTs they exhibited the tortuosity characteristic of high endothelial venules (HEVs; **Fig. 3g**)^51^. Nerve fibres were ∼5-fold more numerous than blood vessels when expressed as length per unit tissue volume (at ∼100 µm of fibre per 1000 µm³ of tissue). The opposite was true in SILTS however: vascular density was ∼20% greater than in the lamina propria but nerve density was 32-fold lower. Thus, nerves commonly formed a dense basket surrounding SILTs (**Fig. 3h**); the few nerves the penetrated the SILT were tightly associated with the vasculature.

### Volumetric virtual H&E imaging

One limitation of multiplexed immunofluorescence (mIF) relative to conventional H&E staining is that, even in high-plex images, substantial populations of cells remain unlabeled and acellular structures such as the ECM and basement membranes are incompletely visualized, often due to the lack of definitive markers^52^. However, eosin cannot be fully photobleached and its use therefore precludes parallel or subsequent IF. To mitigate this, we implemented a “virtual H&E” approach (vH&E)^53^ in which tissues underwent antigen retrieval and clearing without photobleaching followed by staining with DAPI. DAPI approximates haematoxylin staining of nuclei while autofluorescence (at 488 nm excitation/510-550 nm emission wavelengths) approximates eosin staining of cytoplasm and ECM. A Beer-Lambert transformation was applied to images to convert fluorescence intensities into pseudo-absorbance values^53,54^ and colour mapping used to generate blue-purple nuclear and pink stromal/cytoplasmic features resembling conventional H&E staining (**Fig. 4a**). The use of low-plex v-CyCIF following vH&E made it possible to localize specific cell types within the overall tissue architecture (illustrated for CD4^+^ T cells in DS5; **Fig. 4b**. The Beer-Lambert framework also enabled generation of “brown stain” images resembling chromogenic immunohistochemistry (IHC) performed using 3,3’-diaminobenzidine (DAB) chemistry (**Fig. 4c**); this facilitates comparison v-CyCIF and conventional 2D IHC, which is common in pathology. vH&E images could also be segmented using computational pathology algorithms^55,56^, enabling quantitative analysis of tissue architecture (**Extended Data Fig. 4 a-d**). For example, colonic crypts in DS5 were estimated to have a volume of ∼1.4 ± 0.55 × 10⁶ µm³ (mean ± s.d., n = 60 crypts; **Fig. 4d-f**) consistent with historical estimates of 0.6 to 2.2 × 10⁶ µm³ ^57–59^. vH&E followed by low-plex v-CyCIF is particularly advantageous with complex networks of nerves and vessels, as illustrated for TUBB3-stained tissue (**Fig. 4g,h; Supplementary Video 2**).

**Figure 4:**
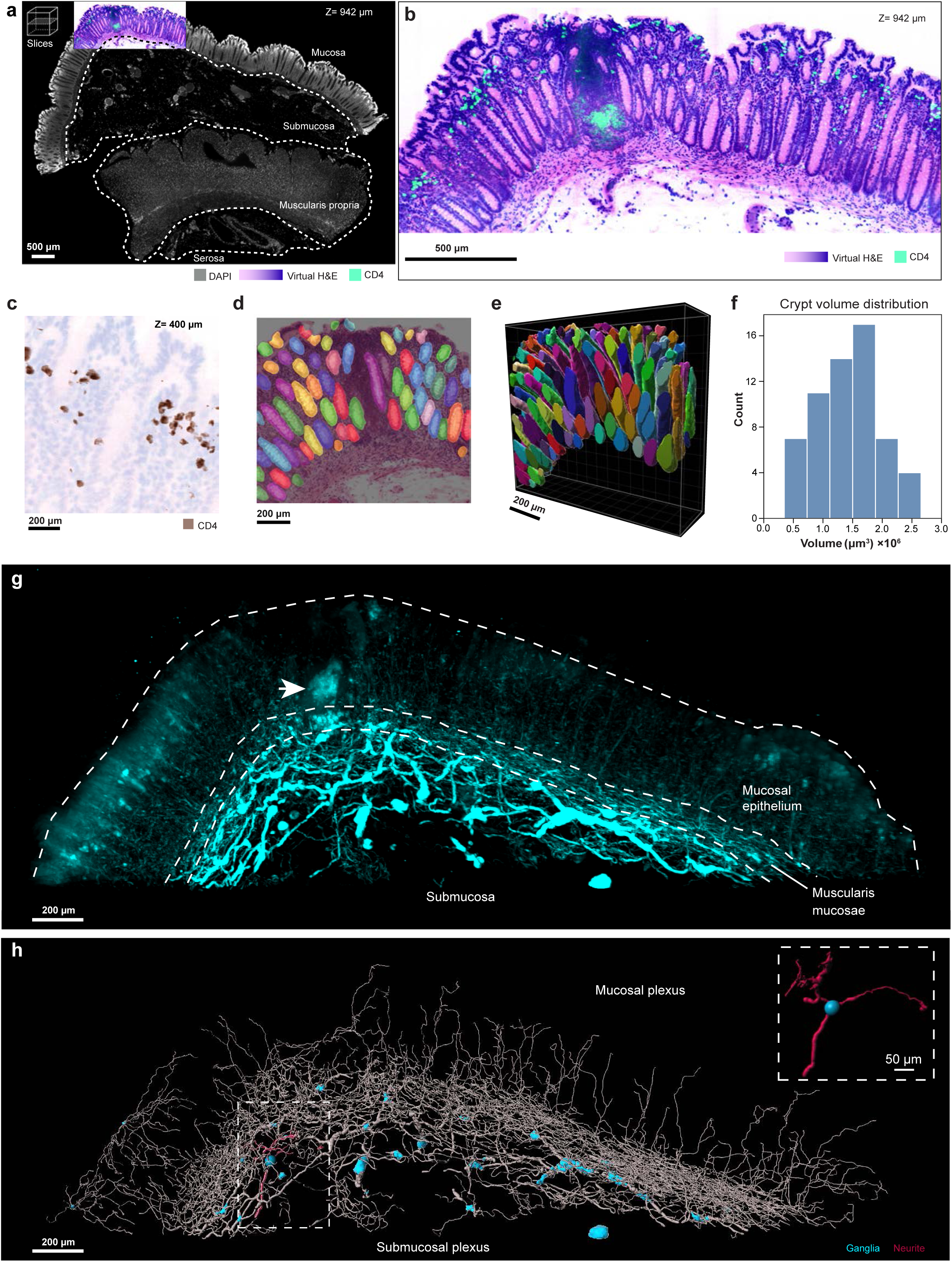
Segmentation and analysis of 3D virtual H&E. **a,** Annotation of colon regions including mucosa, submucosa, muscularis propria, and serosa, outlined with dashed lines. Scale bars: 500 µm. **b,** Single optical plane at 942 μm of virtual H&E showing the cross section of the colon crypts beside a CD4^+^ SILT (cyan). **c,** Single optical plane from virtual CD4 IHC stain at depths of 400 μm from the same tissue volume. Scale bar: 200 μm. **d,** Binary labelled masks of segmented crypts overlaid onto a single optical plane of virtual H&E acquired with 15×/0.4 NA objective on the megaSPIM light-sheet microscope. Crypts segmented via a sparse autoencoder (SAE) to select morphology-rich patches, followed by U-Net based segmentation trained on token-level targets from the UNI foundation model. **e,** 3D rendering individual colonic crypts selected for minimal truncation and overlay view in vH&E; the histogram of crypt volume distribution. **f**, Histogram of crypt volume distribution. **g,** Maximum projection intensity view of the colon showing the distribution of enteric neural structures (cyan) relative to anatomical layers. Dashed outlines delineate the epithelium, submucosa, and muscularis mucosae. Scale bars: 200 μm **h,** 3D rendered view of the enteric plexuses illustrating the mucosal plexus and submucosal plexus. Ganglia are shown in cyan and neurites in magenta; grey traces depict the broader neurite network. Dashed box highlights a representative ganglion with associated neurites. Scale bars: 50 μm.

### Neuroimmune interactions in the normal human colon

Crosstalk between the enteric nervous system and immune cells is a fundamental feature of colon homeostasis. In CRC, innervation can promote cancer progression via secretion of neurotransmitters and neuropeptides; conversely, cancer cells stimulate nerve growth via section of growth factors such as NGF^60,61^. To map neural-immune interactions in normal colon, we used both confocal-based (DS6; **Fig. 5a-b**) and LSFM-based (DS7; Fig. 6a-d) v-CyCIF. Crypts were visualized by staining epithelial cells with anti-CLDN7 (blue, **Fig. 5a-b**), the surrounding network of enteric nerves with TUBB3 (yellow), blood vessels with CD31 (green), and conventional T cells with CD8 (pink). Nerve fibres were found to have a mean diameter of 3.8 ± 1.2 µm, consistent with small axonal bundles (**Supplementary Fig. 8a, b**), and blood vessels a diameter of 6.75 ± 0.60 µm, consistent with individual capillaries. Both nerves and blood vessels were predominantly oriented along the major axis of the crypts (**Supplementary Fig. 8c; Supplementary Video 3**). Following single-cell segmentation^40–42^ a k-nearest neighbour analysis of CD8^+^ T cells revealed a non-uniform distribution within the lamina propria, the connective tissue underlying the colonic epithelium (**Fig. 6c, d**). When the Euclidean distance transform was computed, ∼40% of CD8 T cells were found to be in direct contact with nerve fibres (down to the resolution limit of the analysis, ∼ 0.7 µm; **Fig. 6e**). To confirm that this exceeded what would be expected of a random distribution, we compared the cumulative density function (CDF)^17,62^ of observed T cell-to-nerve distances (Fig. 6f, blue curve) with 1,000 simulations of complete spatial randomness (CSR; grey curve). The observed distribution fell outside the 98% confidence envelope of the CSR simulations, demonstrating a non-random spatial association between immune cells and nerves. An attraction distance of ∼9 µm (red line), defined by the intersection of observed and randomized probability density functions, marked the range over which T cells were enriched near nerves relative to random expectation. Moreover, when T cells were colour-coded according to nerve proximity, cells in direct contact with nerve fibres (black) localized preferentially along crypt walls rather than at crypt tips; this was not observed for T cells more distant from nerves (magenta; **Fig. 6g**; **Supplementary Fig. 9a, b**).

**Figure 5:**
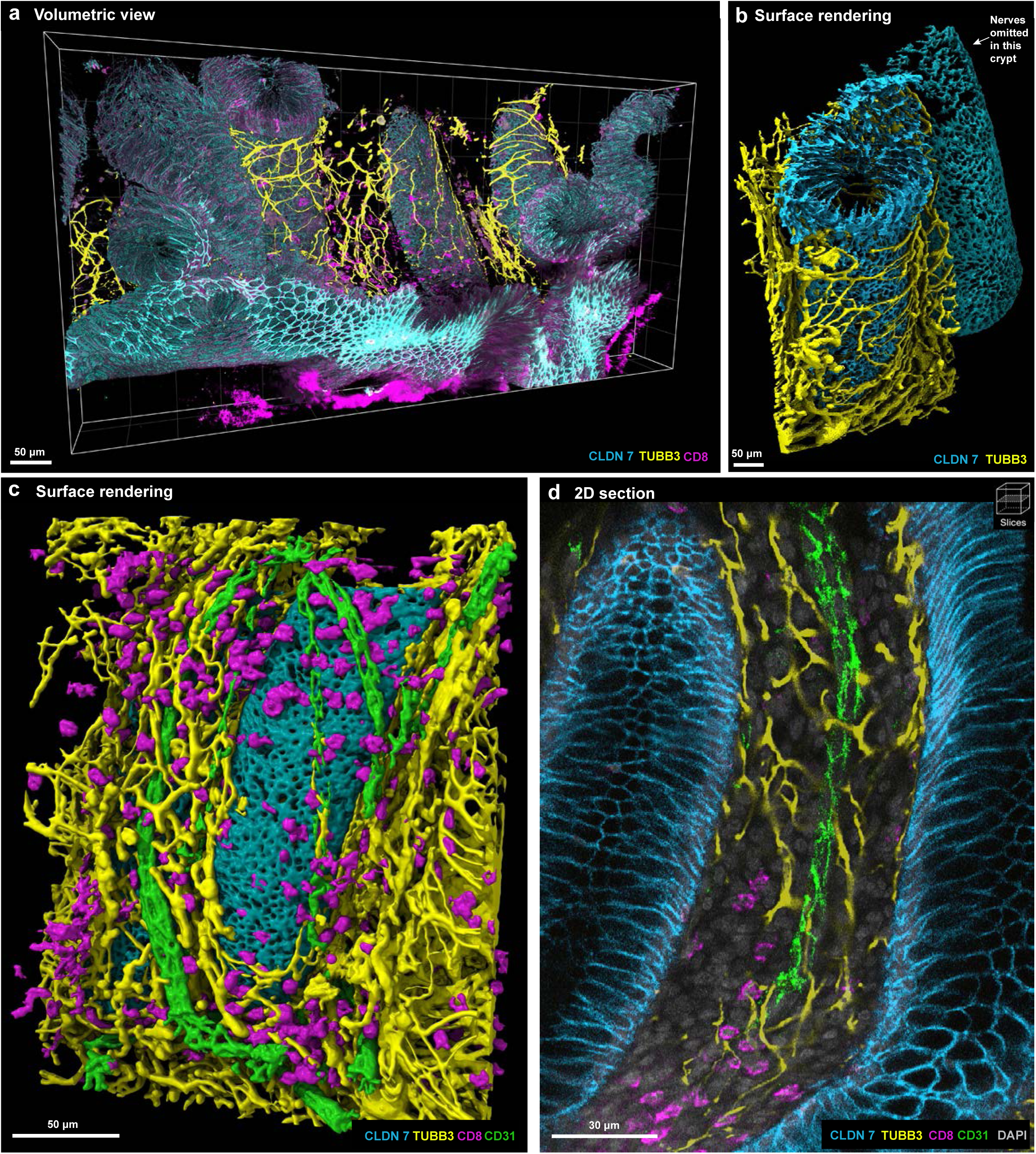
High-resolution mapping of nerve fibre geometry and vasculature. **a,** 3D volumetric views showing spatial relationships between CLDN7+ epithelium (cyan), TUBB3+ nerve fibres (yellow), and CD8+ T cells (magenta) within the same volume. Scale bars: 50 µm. **b,** 3D surfaces rendering of a cleared adjacent normal colon tissue showing Claudin7+ (CLDN7) epithelial structure (cyan) with TUBB3+ neuronal network (yellow). Scale bar: 50 µm. **c,** 3D surface rendering of Claudin7+ epithelium (cyan), TUBB3+ nerve fibres (yellow), CD31+ vasculature (green), and CD8+ T cells (magenta). Scale bar: 50 µm. **d,** Representative 2D section illustrating CLDN7+ epithelial structure (cyan) flanked by the stromal compartment containing TUBB3+ nerve fibres (yellow), CD31+ vasculature (green), and CD8+ T cells (magenta). Scale bar: 30 µm.

**Figure 6:**
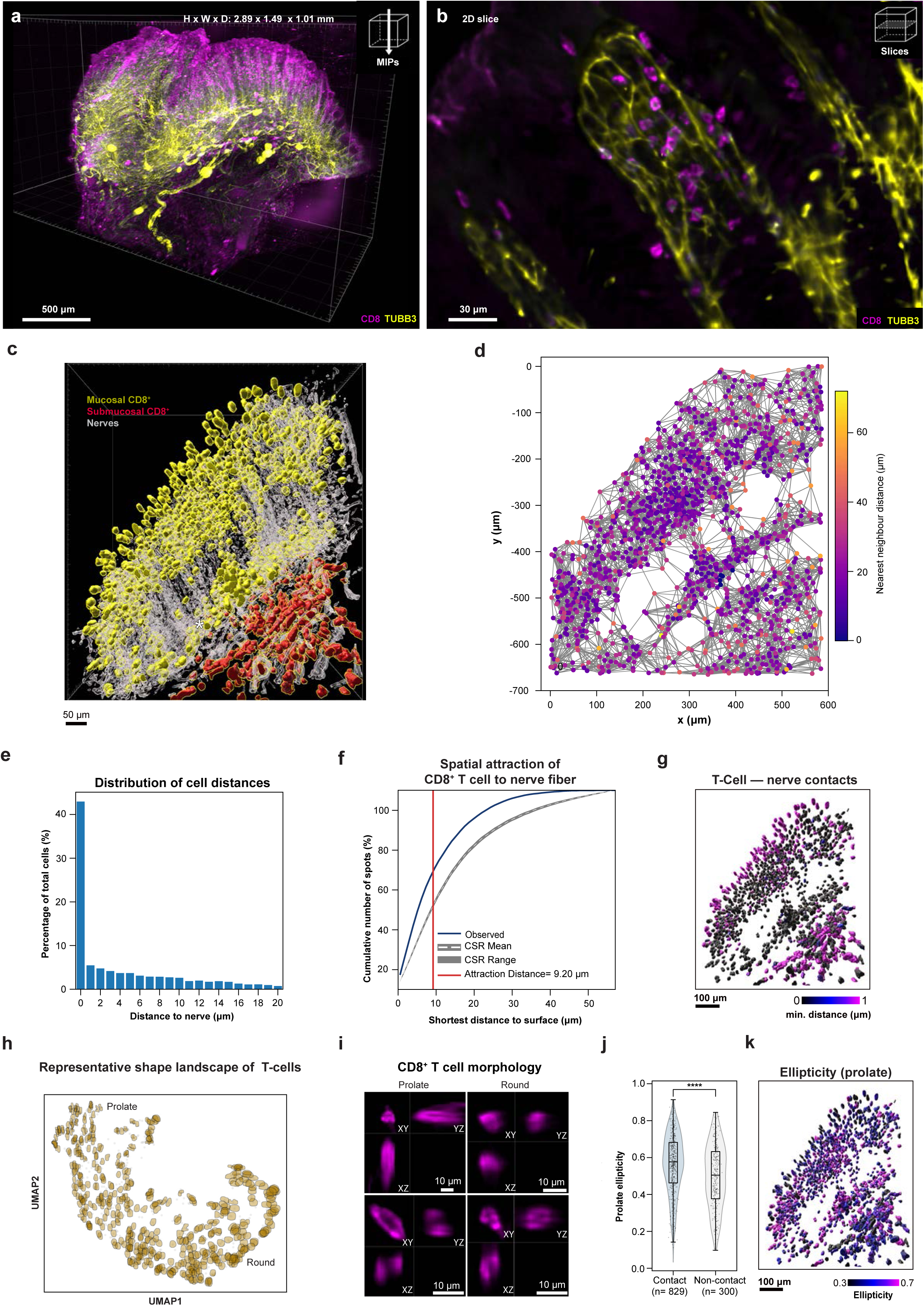
Neuroimmune interactions in normal colon. **a,** 3D volumetric view of cleared FFPE human colon (2.9 x 1.5 x 1.0 mm) with TUBB3+ for neurons and CD8+ for T cells and imaged on MuVI-SPIM with an optical resolution of 650 x 650 x 2000 nm (W x D x H). Scale bar: 500 μm. **b,** zoomed-in view of single optical plane from the same volume in **(a)** showing physical interactions of CD8^+^ T cells and nerves in the lamina propria. Scale bar: 30 μm. **c,** Segmentation of individual T cells (assorted colours) onto nerves (grey) from a cropped ROI demonstrating spatial separation between T-cell populations of the mucosa (yellow) and submucosa (red) regions. **d,** XY view of the reconstructed 3D CD8+ T cell network based on nearest-neighbour distances. **e,** Relationship between T-cell density (shown as percent of total cells) and nerve proximity within 20 μm. **f,** Spatial attraction analysis also shows strong attraction of CD8 T cells to nerve fibres with an attraction distance of 9.2 µm. At this distance, 96% of observed spots were located near the surface compared with 39% expected under CSR, corresponding to a 56% enrichment. **g,** Spatial distributions of T cells annotated by nerve contact. Colour denotes contact distance, increasing black intensity (“0”) indicates greater association with nerves. **h,** UMAP embedding of CD8⁺ T-cell morphological features. **i,** Orthogonal view of distinct CD8+ T cell morphology in the normal colon from. Scale bar: 10 μm. **j-k,** Comparison of CD8^+^ T cell morphology (prolate ellipticity) between nerve-contact and non-contact populations. Colour denotes ellipticity, with higher values corresponding to a more elongated cell shape. Scale bars:100 μm.

Visual inspection suggested that T cell morphology also varied with nerve proximity. To quantify this, segmented cell surface outlines were embedded using a Dynamic Graph Convolutional Neural Network^63^. Visualization of T cells along different axes confirmed that perceived differences in morphology were not simply due to differences in resolution along the X, Y and Z axes (**Fig. 6h, i**). Instead, T cells in direct contact with nerves were significantly more likely to exhibit prolate morphology than more distant cells (**Fig. 6j, k, Supplementary Fig. 10**). Deviations from a spherical shape have previously been associated with T cell activation^64–66^ and spatial maps of T cell-nerve contact closely paralleled maps of cellular ellipticity (**Fig. 6g, k**; **Supplementary Fig. 11**). Together, these data demonstrate non-random association between nerves and T cells in normal human colon and illustrate the potential of volumetric imaging to quantify neuroimmune interactions across tissue compartments.

### Germinal centre organization in tertiary lymphoid structures

TLS arise ectopically in response to chronic inflammation^6,67^ and have been studied extensively in cancer, in which their presence is associated with extended disease-free survival^68^ and improved responses to immune checkpoint inhibitor (ICI) therapy in some cases^69^. However, the role of TLS in natural and ICI-mediated therapy remains only partly understood, reflecting the wide diversity of TLS states and phenotypes. This arises in part because TLS formation is a multi-step process in which accumulation and sorting of B and T cells is followed by assembly of a cage-like structure comprising interdigitated FDCs to form a GC zone within which antigen-specific B and T cell programming and activation occurs^70^. Early TLS contain B cells but not FDCs, primary TLS contain FDCs positive for complement receptor type 2 (CD21), and mature secondary TLS contain FDCs positive for both CD21 and the IgE receptor CD23. Classically, CD23 is restricted to light-zone FDCs, which sustain somatic hypermutation and affinity-based selection of antibodies occurs. We used v-CyCIF to study TLS organization in CRC specimens and found that some had the expected ovaloid shape (**Extended Data Fig**. **7;** DS8) with CD21^+^ CD23^+^ FDCs nested within an CD21^+^ FDC network (**Supplementary Fig. 12a-c)**. However, many GCs had a highly irregular morphology with multiple branches that could be traced continuously through the full tissue volume (DS9; **Fig. 7a-i** and DS10**; Extended Data Fig**. **8a,b**). Strikingly, over half of TLSs examined exhibited distinct CD23^high^ regions overlapped only partially with CD21^+^ FDCs (**Fig. 7j,k**; **Supplementary Table 4**); TLS-6 was the most extreme example with up to 80% of the FDC network CD23^+^ and CD21^−^, but with retention of the cage-like morphology. This phenotype was not observed in SILTs in which CD23 staining was restricted to FDCs that were also CD21-positive (**Fig. 7l-p**). To our knowledge, this diversity of FDC phenotypes in GCs has not previously been observed, most likely because the complex shapes of GCs are not evident in virtual 2D sections. Moreover, since CD21 and CD23 are expressed by cells other than FDCs, 3D makes it possible to identify the subset of cells having interdigitated, dendritic morphologies. Our findings are consistent, however, with emerging evidence that GCs found in mature TLS can contain distinct compartments with differing regulatory properties^71,72^.

**Figure 7:**
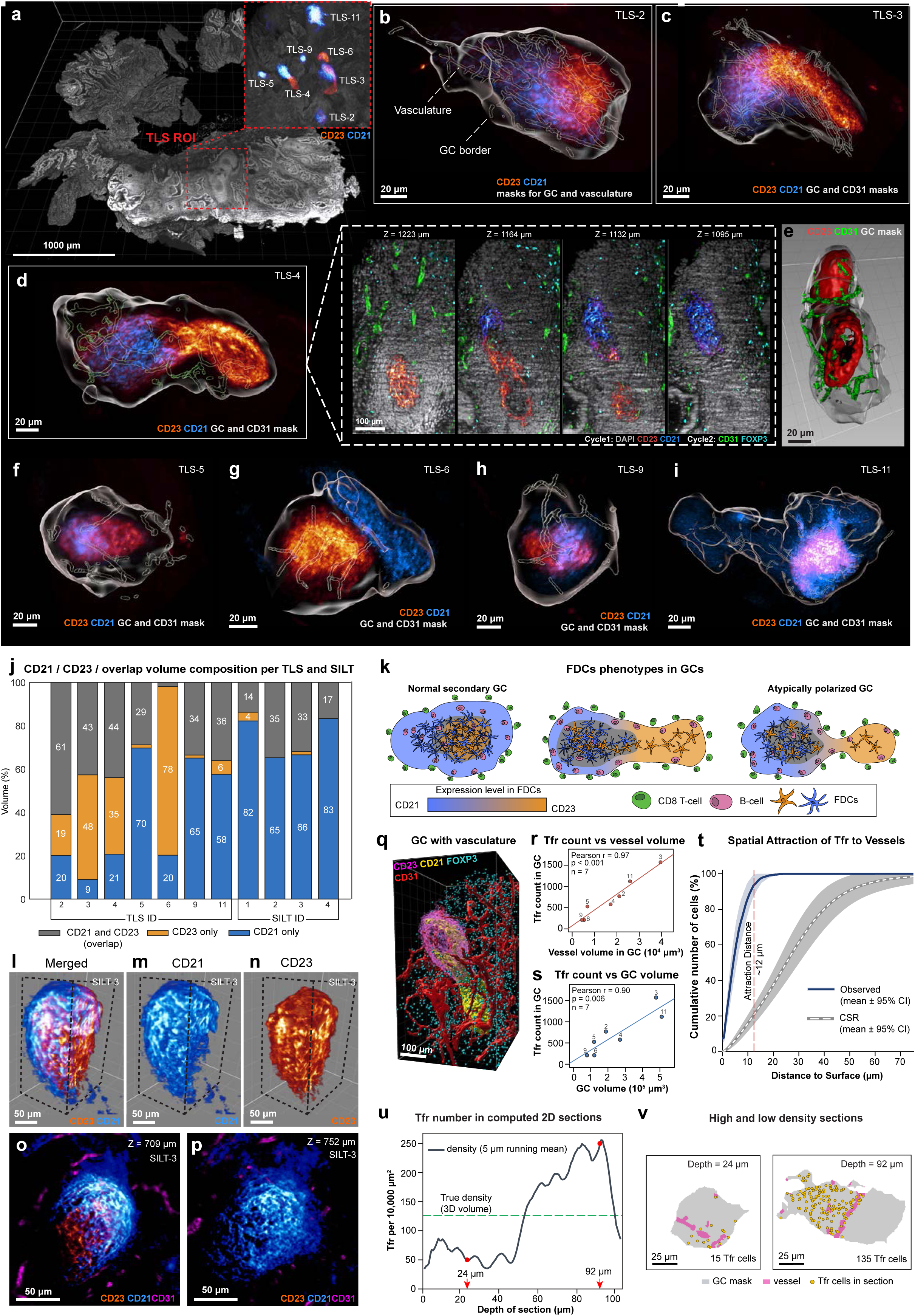
Spatial organization of follicular dendritic cell network and Treg–vascular associations in colon tertiary lymphoid structures. **a,** 3D rendering of the cleared colorectal cancer tissue (DAPI, grey) at 3.6× magnification. The red dashed outline marks the tertiary lymphoid structure (TLS) region of interest selected for high-resolution analysis (9× magnification). **b–i,** Volume renderings of individual TLS (identified as TLS-2, 3, 4, 5, 6, 9 and 11, TLS-1, 7, 8, 10 were excluded due to the lack of CD21 or CD23 expression) showing CD23 (glow) and CD21 (blue) within the segmented germinal centre mask (transparent grey) and CD31+ vascular mask (contours). Dashed box, expanded from TLS-4 (d), showing single optical sections through TLS at different z depths, acquired over two cycles: cycle 1, DAPI (grey), CD23 (glow) and CD21 (blue); cycle 2, CD31 (green) and FOXP3 (cyan). **e,** 3D rendering view of TLS-4 (d). **j,** Stacked-bar quantification of the volume fraction occupied by CD21 alone (blue), CD23 alone (orange) and the region of overlap (CD21 and CD23 double positive, grey), computed per segmented structure for seven TLS and two solitary intestinal lymphoid tissues (SILT); values are percentages of total segmented follicular volume and sum to 100% for each structure. **k,** Schematic illustration summarizing the observed organization: a CD21-to-CD23 expression gradient (blue to orange) partitions the follicular dendritic cell network into CD21+ (blue) and CD23+ (orange) subsets, with B cells (pink) concentrated within the FDC network and CD8+ T cells (green) restricted largely to the perimeter; the three cartoons depict the range of compartment arrangements seen across structures. **l–n,** 3D renderings of SILT-1 showing the merged CD21/CD23 signal (l) and each channel separately: CD21 (m, blue) and CD23 (m, glow). **o-p,** Single optical sections through SILT-1 at z = 709 μm (n) and z = 752 μm (o) showing CD23 (red), CD21 (blue) and CD31 (magenta). **q,** 3D rendering of the TLS-4 germinal centre showing CD23 (magenta), CD21 (yellow), CD31 (red) and FOXP3 (cyan). **r-**s, Correlation between the number of FOXP3+ Tfr cells within the germinal centre and CD31+ vessel volume (r) and total germinal-centre volume (s). Each point is one TLS, labelled with its TLS ID; Two-sided Pearson correlation: r = 0.97, P < 0.001 (r) and r = 0.90, P = 0.006 (s); Lines show the least-square linear fit. **t,** Spatial attraction analysis of FOXP3+ Treg cells to CD31+ vessel surfaces, plotted as the mean cumulative distribution of cell-to-surface distances. The observed distribution (blue, mean ± 95% confidence interval) is shifted towards the vessel surface relative to a complete spatial randomness (CSR) null model generated by (n = 1000) randomizations (grey dashed, mean ± 95% CI); the red vertical line marks the attraction distance (∼12 μm). **u,** Tfr cell density plot across virtual 2D sections, depths were selected whose density lay closest to 50 (z = 24 μm) and 250 Tfr cells (z = 92 μm) per 10,000 μm². **v,** Tfr cells (yellow), vessels (magenta), and GC mask (grey) are shown for low and high Tfr density sections respectively. Scale bars, 1,000 μm (a), 100 μm (inset, d), 50 μm (l–p), 25 μm (v) and 20 μm (b–i).

Follicular regulatory T cells (Tfr) are a set of specialized FOXP3^+^ CD4⁺ T cells that maintain GC homeostasis and, in cancer, suppress anti-tumour immunity^73–76^. The presence of TLS with high proportions of Tfrs may also curtail the efficacy of immune checkpoint immunotherapy^77^. When we quantified the number of Tfrs per GC, we found that it correlated strongly with GC volume (r = 0.97, P < 0.001) implying a density of ∼2.5 Tfr per 1000 µm^3^ of GC volume. The fractional volume of blood vessels (most likely high HEVs) was also constant across TLS at ∼38 ± 12 µm of blood vessels per 1000 µm^3^ of GC volume; **Fig. 7q-s**). Locally, Tfr cells were significantly associated with blood vessels (**Fig. 7t and Supplementary Fig. 13a,b**), consistent with delivery of precursors cells (e.g. Tregs) or regulatory factors from the vasculature (this was also true in SILTs^78^). However, in clinical specimens Tfr numbers are commonly evaluated using 2D histological specimens. We found that when virtual 2D sections of a single GC were generated from a 3D volume, 10% of sections contained no Tfr at all and when present, Tfr number varied 5-fold or more across sections: conversely, only 20% of 2D sections examined yielded a value similar to 3D density (**Fig. 7u,v** and **Supplementary Video 4**). These data demonstrate a scaling relationship between GC volume and the volume of the enclosed vasculature, and between Tfr numbers and local vasculature density. Since HEVs are discrete, tortuous, structures of irregular diameter, it follows that Tfr density will also fluctuate locally. However, Tfrs are migratory (at speeds >10 μm per min.)^79^ and it is reasonable to assume that their density in a full GC volume (as estimated by 3D imaging) is the more meaningful than density in a 5 µm slice of that volume. Tfr abundance in human mesenteric lymph nodes has also been observed to fluctuate locally, but remain nearly constant when normalized by volume^80^. These findings illustrate errors that arise when subsampling a complex 3D structure in 2D without consideration of the microanatomy. Since TLS density and maturation state have been incorporated into potential prognostic scoring systems for CRC^81^ based on standard 2D histology sections, we speculate that knowledge of volumetric scaling relationships may improve such assessments by better accounting for natural fluctuations in cell density.

### Multiscale profiling of tertiary lymphoid structures by re-embedding

TLSs and other immune structures contain a variety of specialized immune cell types in addition to Tfrs; the activation states of these cells and their interactions and determine the functional status of the TLS. Functionally distinct immune cells commonly share markers, making it essential to rely on high-dimensional marker combinations to establish specific state. Cells in lymphoid aggregates are densely packed, however, making single-cell phenotyping difficult in both conventional 2D mIF images obtained at “standard” scanner resolutions^14^ and in LSFM-based v-CyCIF. To address these limitations, we developed a workflow for re-embedding tissue specimens in paraffin following volumetric imaging, enabling subsequent sectioning and mounting on standard glass slides for high-resolution CyCIF imaging with of thick sections. As previously described^14^, this approach to imaging 30-50 µm thick specimens enable very precise discrimination of cell processes and subcellular organelles in one to two layers of intact cells whereas conventional imaging of 5 µm thick specimens results in fragmentation of >90% of cells. To illustrate this approach, we acquired a 6-plex (4-cycle) v-CyCIF image of a 1 mm thick CRC specimen (DS11). Tumour cells were labelled with caudal-type homeobox 2 transcription factor (CDX2; yellow) and TLSs were identified by CD21 staining (green; **Fig. 8a**)^82^. More than 10 TLSs were identified within this specimen (**Fig. 8a**). Following v-CyCIF, the specimen was re-embedded and sectioned on a microtome using volumetric images to guide selection of germinal centre-containing 40 µm thick sections (DS10; hydrated thickness), which then underwent 25-plex high-resolution thick-section CyCIF imaging on a confocal microscope (**Supplementary Table 6; Extended Data Fig. 5a-c**).

**Figure 8:**
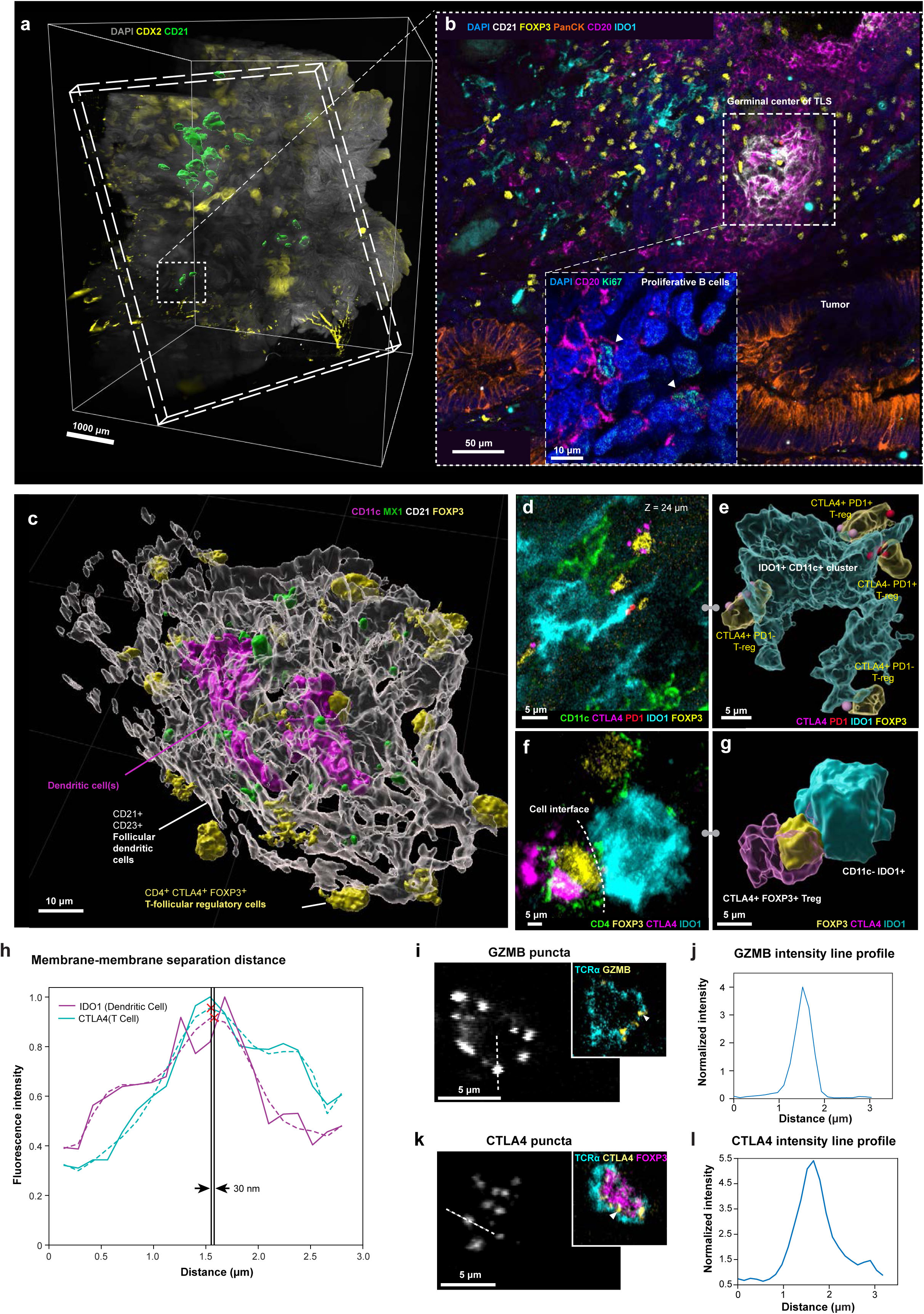
High-resolution 3D-CyCIF of re-embedded tissue reveals an immunosuppressive FOXP3 - IDO1 axis in tertiary lymphoid structures. **a,** 3D volumetric view of a TLS with CD21 (surface rendered in green) CDX2 (yellow) and DAPI (Grey) imaged on MuVi-SPIM light-sheet microscope. Dashed white box highlights a TLS cluster (green) selected for downstream 3D high resolution CyCIF on laser scanning confocal microscopy. Scale bar: 1000 μm. **b,** Cropped max intensity projection of 18 optical planes from 25 µm thick re-embedded tissue imaged with 3D CyCIF. CD21 (white), FOXP3 (yellow), PanCK (orange) CD20 (Magenta), IDO1 (Cyan) are shown. White dashed box indicates a germinal centre, tumour region is marked by PanCK (orange). Scale bars: 50 µm. Zoomed-in view of Ki-67+ proliferative B cells (centroblasts) in the TLS. Scale bars: 10 µm **c,** Surface mesh of Tfr cells interacting with FDC in the germinal centre. CD11c (magenta), MX1 (green), CD21 (white), and FOXP3 (yellow) are highlighted. Scale bars: 10 µm. **d,** A ROI from (**b)** shows CTLA-4^+^ or PD-1^+^ FOXP3^+^ Treg interactions with IDO1^+^ cell cluster. **e,** surface meshes of Treg-IDO1 interactions from (**d**). Scale bars: 5 µm (**d-e**). **f**, 2D section shows CTLA-4^+^ Treg interaction with IDO1^+^ stromal cell. Scale bars: 15 µm. **g**, 3D surface rendering of interactions from (**f**). **h,** membrane interaction graphs between IDO1^+^ dendritic cell (cyan, IDO1 channel) and Treg (magenta, CTLA-4 channel) from (**e-g**). **i,** Single-channel views of GZMB puncta (white), white dashed line indicates the transect used for intensity profiling. Inset shows a representative image of GZMB^+^ cytotoxic T cell. Scale bars: 5 µm. **j** Line-intensity profiles along the transect for GZMB. **k,** Single-channel views of CTLA-4 puncta (white), white dashed line indicates the transect used for intensity profiling. Inset shows a representative image of CTLA-4^+^ FOXP3^+^ regulatory T cell. Scale bars: 5 µm. **l,** Line-intensity profiles along the transect for CTLA-4.

We found that high-resolution imaging of remounted specimens addressed three limitations of LSFM-based v-CyCIF: (i) single-cell phenotyping in densely packed tissue (ii) detection of cell-cell interactions including juxtacrine signaling and points of close membrane apposition (iii) quantifying intracellular organelles and condensates (**Supplementary Fig. 14** and **Supplementary Video 5**). As an example of single-cell phenotyping, we found that Ki67 positivity could be localized to individual CD20^+^ B cells and other cell types within a GC and CD21^+^ CD23^+^ FDCs could be distinguished from CD11c^+^ conventional dendritic cells, which have highly irregular morphologies (**Fig. 8b, c**). Biologically significant cell-cell interactions not apparent at lower spatial resolutions^14^ included CD4^+^ CTLA4^+^ FOXP3^+^ Tfr cells interacting with CD11c^+^ dendritic cells positive for indoleamine 2,3-dioxygenase 1-positive (IDO1; this is thought to be a tolerizing interaction that promotes Tfr differentiation; **Fig. 8e-g; Extended Data Fig. 5**).^83^ Using previously described methods^14^, membrane-membrane distances in these interactions could be resolved down to ∼40 nm, consistent with direct membrane apposition and local paracrine signaling (**Fig. 8h, Supplementary Fig. 15**). Many organelles, biomolecular condensates, and receptors appear as diffraction-limited puncta in the cytoplasm or on the cell membrane; such puncta are difficult to detect in low-resolution images. Examples of puncta whose quantification benefitted from re-sectioning and high-resolution imaging were granzyme B-containing cytotoxic granules in CD4^+^ and CD8^+^ T cells (**Fig. 8k, l**) as well as immune checkpoint regulators such as CTLA-4 and PD-1 in T cells (**Fig. 8d, e, i, j**).^14,84^ Overall, these data demonstrate the value of re-embedding tissue following LSFM imaging for either standard imaging of ∼5 µm sections or thick-section high-resolution confocal imaging that preserves local tissue architecture in a flexible workflow (**Supplementary Fig. 16**).

## DISCUSSION

Multiplexed v-CyCIF imaging enables routine high-plex 3D analysis of FFPE specimens, the type most readily available from humans and mouse models. v-CyCIF methods have been implemented on multiple LSFM and confocal microscopes with no requirement for specialized instrumentation or proprietary reagents. In colon no appreciable tissue degradation has been observed over at least 16–17 cycles (i.e. a ∼40 plex image), suggesting that, as additional validated antibodies become available, it will be possible to achieve as high plex in 3D as in 2D spatial profiling (i.e. >100-plex if desired). v-CyCIF clearing and staining protocols have been effective to date with specimens up to ∼1 mm thick.

Although this is several-fold less than what is achievable with conventional strongly delipidating clearing reagents, it is sufficient for visualization of many mammalian FTUs. Prior to antibody staining, inexpensive and non-destructive 3D v-H&E imaging can also be used to visualize tissue architecture, guide subsequent IF imaging, and enable computational analysis using pathology foundation models^55^. Critically, the development of new clearing protocols that do not fully delipidate specimens, and thus, do not destroy plasma membranes, makes it possible to use membrane-localized CD antigens for immune cell subtyping.

v-CyCIF increases specimen thickness 200-fold relative to conventional histology while also accommodating specimens up to 1-2 cm in lateral dimensions, making it a whole-slide imaging method.^85^ However, greater tissue thickness and size comes at the cost of lower resolution. Resolution in LSFM is heavily dependent on instrumentation, but with the commercial instruments used in this paper it averaged ∼700 nm laterally. This is comparable to conventional slide-scanning spatial profiling instruments, but substantially less than what can be achieved with confocal-based 3D high-plex CyCIF (∼120 nm lateral resolution in super-resolution mode)^14^. A key feature of our approach is that it allows millimetre-thick tissues subjected to LSFM-based v-CyCIF to be re-embedded, sectioned at thicknesses between 5 and 50 µm, and imaged again at high resolution. In these images cells can be successfully segmented in the crowded environment of GCs, membrane-membrane separation can be resolved down to 30∼50 nm, and punctate structures and juxtacrine signalling complexes can be detected and quantified. Future integration of X-ray Computed Micro- and Nano-Tomography (microCT; nanoCT)^86^ promises to extend imaging of intact tissue to centimetre scale and focused ion beam scanning electron microscopy (FIB-SEM) to allow cell-cell contacts and organelles to be resolved down to nanometre resolution. Together, such approaches have the potential to generate comprehensive multiscale atlases of human tissues from a variety of diseases.

We explore the uses of v-CyCIF primarily using the colonic mucosa and CRC resections, which includes layers of epithelial, muscle, and connective tissue as well as extensive infiltration by both dispersed immune cells (in the lamina propria) and highly organized lymphoid aggregates such as SILTs and TLSs. These immune cells play important roles in gut health and in multiple diseases including cancer (particularly tumour immunosurveillance and responses to immune checkpoint inhibitors)^77,81^ and in Crohn’s disease^87^. We find that dispersed T cells in the human lamina propria are significantly associated with peripheral nerves, which are abundant in the gut, extending what has been shown previously in Crohn’s patients^88^. Nerves are significantly less abundant in lymphoid aggregates than in the lamina propria and instead form a dense surrounding basket. Those nerves that penetrate SILTs and TLS course alongside blood vessels (which likely represent HEVs). These anatomical arrangements have not previously been imaged in humans in 3D but we interpret them to represent classical neuro-immune-vascular coupling (triad interactions)^89^. In CRC, we find that follicular dendritic cells in GCs in TLS (but not SILTs) have an abnormal maturation pattern with implications for prognostication schemes reliant on TLS abundance and maturity. In future work, pairing this framework with perturbation-based approaches in patient-derived 3D models^90–93^ will help to determine how these structures form, mature, and influence antitumour immunity.

### Limitations and future directions

This paper has two primary limitations. First, insufficient fluorophore-labelled primary antibodies have been validated (for active staining) to go beyond ∼40-plex imaging, although achieving this is simply a matter of time and cost. Over 500 antibodies have been validated to date for 2D CyCIF and ∼60% of the first 72 antibodies tested have proven suitable for v-CyCIF. One difference between antibodies that perform well in 2D CyCIF and in v-CyCIF appears to be antibody-antigen dissociation rate: the extended time required for staining and imaging thick specimens by v-CyCIF requires stable binding in combination with good tissue penetration. These are not properties that are easy to predict. Thus, expanding the available v-CyCIF antibody repertoire is a matter of empirical testing across tissue types and staining conditions.

A second limitation involves the difficulty of accurately registering, segmenting, and analysing high-plex multi-terabyte LSFM datasets at full resolution. This can be addressed in part by new sample holders that better account for LSFM geometry and ensure repeatable positioning of specimens relative to the optical path to reduce sample movement (simply fixing the specimen to the glass coverslip with glue impairs staining, bleaching and clearing). However, the primary remaining challenge with v-CyCIF involves data processing and computation. Substantial innovation will be required to create methods that align and segment large high-plex images while accounting for optical aberrations, specimen distortion, and large file sizes. To promote such developments, we have made the data in this paper freely available in several Bio-Formats^94^ and Imaris-compatible formats (.ims, HDF5-based).

## EXTENDED DATA LEGENDS

**Extended Data Figure 1: v-CyCIF using spinning disk and laser scanning confocal microscopy**

**a,** 3D rendering of a cleared human colon specimen imaged on spinning disk confocal. αSMA^+^ vasculature (cyan) and Ki67 (magenta) rendered as surfaces. Scale bar, 500 µm. **b,** Close-up view of crypts from (**a**) of Ki67^+^ proliferative cells (magenta), nuclei (DAPI, grey) and vasculature (αSMA, cyan) within crypts. Scale bar: 100 µm. **c,** Single optical plane of the colon region. Scale bar: 100 µm. **d,** Maximum intensity projection image of the cleared human colon imaged on laser-scanning confocal microscopy (LSCM), labelled with E-CAD (green), αSMA (yellow) and CD31 (red). Scale bar: 30 µm. **e,** LSCM orthogonal views (XY, XZ and YZ). Scale bar: 100 µm.

**Extended Data Figure 2: Refractive-index matching, sample holder design, and fluorescence imaging before and after photobleaching**

**a,** MegaSPIM holder showing an agarose-embedded tissue mounted in a 3D-printed frame and secured in a “click-in” holder for dual-objective illumination and detection. **c,** MuVi-SPIM holder design, including the 3D frame, fastener, and click-in holder used to position the agarose-embedded sample within the dual illumination setup. **d,** DALISPIM side view showing the glued frame holder for a 6-well plate, with the sample immersed in oil and positioned between the illumination and detection objectives inside a sealed imaging setup.

**Extended Data Figure 3: Additional v-CyCIF examples demonstrating applicability across organs and disease contexts.**

**a,** 3D volumetric view of adjacent normal breast tissue highlighting ductal and lobular architecture (terminal duct and lobule annotated). Scale bars: 500 μm. **b,** DAPI signal of acinar region (dashed box in **a**) at the z depth of 226 μm. Scale bars: 100 μm. **c,** 3D volumetric view of breast locular structures (DAPI, grey; CK5, magenta). Scale bars: 500 μm. **d,** Single optical plane of (**c**) at the z depth of 2834 μm. Scale bars: 10 μm **e,** 3D volumetric view of melanoma lung metastasis showing tumour marker (SOX10, red) and associated structure (SEPTIN-2, dark yellow). Scale bars: 20 μm. **f,** 3D volumetric view of cutaneous melanoma with tumour marker (SOX10, red) over nuclei (DAPI, grey). Scale bars: 500 μm. **g,** A ROI showing immune-associated signal (CD11c, yellow) in the tumour at the z depth of 1714 μm. Scale bars: 100 μm.

**Extended Data Figure 4: Cell count, region density and SILT quantification across the colon.**

**a,** 3D segmentation of an intact colon specimen showing major anatomical layers (mucosa, submucosa, muscularis, and serosa; colour-coded) with total cell counts per layer and corresponding volumes and estimated cell densities (cells mm^−3^) as indicated. **b,** Identification and quantification of SILTs within the same specimen. Individual SILTs are pseudo-coloured and numbered; labels indicate the cell count for each SILT. Background greyscale signal shows DAPI to provide tissue context. **c,** 3D segmentation and layer-resolved total cell count for an additional colon segment (mucosa and submucosa shown), with estimated volume and cell densities indicated. **d,** SILT detection and cell-count quantification for the specimen shown in (**c**), pseudo-coloured SILTs over a DAPI background. Scale bars: 1000 μm (**a–d**).

**Extended Data Figure 5: Re-embedded 3D CyCIF of germinal centre in tertiary lymphoid structures (TLS)**

**a,** Re-embedded 3D CyCIF image showing spatial organization of a germinal centre defined by a CD21^+^ follicular dendritic cell (FDC) network (white) surrounded by PDPN^+^ lymphatic vessels (green) and CD31+ vasculature (dark blue), with TLS-associated immune populations including FOXP3^+^ regulatory T cells (yellow), CD11c^+^ dendritic cells (magenta), and IDO1^+^ immunoregulatory myeloid cells (cyan). Scale bar: 50 µm. **b,** Complementary marker set highlighting the immune composition of TLS, showing TCRα^+^ T cells (cyan), CTLA-4^+^ T cells (yellow), and GZMB^+^ cytotoxic T cells (magenta) arrayed around the CD21^+^ FDC network (white). Scale bar: 50 µm. **c,** Tissue-context view of the same region showing CD21^+^ FDC organization (white) relative to local tumour microenvironment, including HLA-DPB^+^ antigen presenting cells (yellow), CD31^+^ vasculature (magenta), Vimentin (Vim)^+^ mesenchymal/stromal area (dark blue), Claudin7^+^ tumour epithelial structures (Cyan), and TUBB3^+^ nerve fibres (orange). Scale bar: 50 µm.

**Extended Data Figure 6: v-CyCIF of solitary intestinal lymphoid tissue across cycles**

Representative volumetric renderings (left; scale bars: 1000 µm) and corresponding zoomed-in cross-sectional views (right; scale bars: 200 µm) of intact normal colon tissue imaged on DALISPIM light sheet microscope over 17 iterative cycles, with three to four markers visualized per cycle. Cycles 1–17 are shown; markers for each cycle are indicated in the panel labels.

**Extended Data Figure 7: Isolated intratumoral TLS within a 1 mm adjacent CRC tissue slab**

**a,** Overview of 1 mm CRC tissue slab stained with DAPI. Region of interest (white) shows the location of the TLS within the tumour. Scale bar: 500 µm. **b,** 3D rendering of the FDC network with CD21 (Cyan), CD23 (Magenta) and CD20 (green) surfaces. Scale bar: 100 µm. **c,** Surface rendering of CD31+ blood vessels surrounding the TLS. Scale bar: 200 µm. **d,** Registered images containing CD21 and CD23 in cycle 1, and CD20, FOXP3 and CD31 in cycle 2. Scale bar: 80 µm. **e,** Proliferative ki67+ germinal centre and glandular tumour. Scale bar: 100 µm.

**Extended Data Figure 8: 3D visualization of tertiary lymphoid structures and solitary intestinal lymphoid tissue within colorectal cancer tissue.**

**a,** 3D volume rendering of the cleared colorectal cancer specimen (DAPI, grey) with red boxes marking the locations of individual lymphoid aggregates (5 TLS and 1 SILT) identified within the tissue volume. Scale bar: 1000 µm. **b,** Serial optical sections at increasing z-depth (left to right) through five representative structures (TLS-12, TLS-13, TLS-15, TLS-16, and SILT-5; dashed outlines), stained for CD20 (cyan, B cells), CD23 (yellow, follicular dendritic cells), CD21 (magenta, follicular dendritic cells), and DAPI (grey, nuclei). Scale bars: 100 µm. Right column: 3D surface renderings of the CD23⁺ (yellow) and CD21⁺ (magenta) FDC networks for each structure. Scale bars: 100 µm.

## SUPPLEMENTARY FIGURE LEGENDS

**Supplementary Figure 1: Macroscopic inspection of FFPE tissues during sample preparation for v-CyCIF.**

**a,** Representative photographs of FFPE human colon (top row) and lung (bottom row) specimens at key processing steps. Images show the tissue in the FFPE block, followed by appearance after deparaffinization, after hydration, after methanol bleaching, and after antigen retrieval. **b,** Workflow of v-CyCIF and estimated time for each processing step.

**Supplementary Figure 2: Macroscopic assessment of colon tissue clearing**

**a,** Representative photographs of colon tissue before (left) and after (right) treatment with a refractive index–matching solution (RI = 1.52). Increased visibility of the background grid after treatment indicates enhanced tissue transparency following clearing.

**Supplementary Figure 3: Representative v-CyCIF optical planes across imaging cycles.**

**a,** Single optical plane (z = 364 μm) from a colonic region acquired by MuVI-SPIM in cycle 3. **b,** Single optical plane (z = 636 μm) from the same colonic region in (**a**) acquired by MuVI-SPIM in cycle 4. Neuronal structures are labelled by TUBB3 (yellow) and CD8^+^ T cells by CD8 (magenta). Scale bars: 100 μm.

**Supplementary Figure 4: Cell segmentation across colonic compartments and immune aggregates.**

**a,** Example cell segmentation masks (left; pseudo-coloured by object) and corresponding raw DAPI signal (right) for different compartments (mucosa, SILT, submucosa, muscularis, serosa) in the normal colon. Scale bar: 100 μm (Scale bar: 20 μm for SILT). **b,** Example cell segmentation masks (left; pseudo-coloured by object) and corresponding raw DAPI signal (right) for different compartments in the tumour. Scale bar: 50 μm

**Supplementary Figure 5: Comparison of antibody staining specificity across fluorophores**

**a-b**, Single-channel optical sections from the volume in (**Fig. 3a**) showing CD8 conjugated to Alexa fluor647 (**a**, magenta) and Alexa fluor 488(**b**, green) distribution relative to the SILT.

**Supplementary Figure 6: Orthogonal Views before and after registration of the centre slice in cycle 1-8.**

**a**, Quality control of centre slices from XY, YZ and XZ views showing before and after ANTs registration from image cycles 1-8.

**Supplementary Figure 7: Orthogonal Views before and after registration of the centre slice in cycle 9-17.**

**a,** Quality control of centre slices from XY, YZ and XZ views showing before and after ANTs registration from image cycles 9-17.

**Supplementary Figure 8: Nerve fibre diameter and orientation analysis**

**a,** Zoomed-in view of the TUBB3^+^ nerve fibre from (**Figure 6**) used for fibre analysis in (**b,c**). Scale bar: 20 µm. **b,** Distribution of TUBB3^+^ nerve fibre diameters from the analysed volume (n = 140,490 fibres). **c,** Polar histogram of TUBB3^+^ nerve fibre orientation distribution (colour indicates binned frequency). Red dashed line indicates dominant crypt orientation (117° and 297°).

**Supplementary Figure 9: Spatial analysis of CD8 T-cell morphology in relation to nerve proximity**

**a,** Surface meshes for individual T cells (colour coded, middle), nerve fibres (grey, right) and merged (left) from the zoomed-in ROI in (**Figure 6c**). Scale bar: 70 μm. **b,** CD8+ T cells colour coded by morphological and spatial attributes, including minimum distance to nerves, cell volume, and shape parameters. Scale bar: 100 μm.

**Supplementary Figure 10: Cell shape and size differences associated with cell–nerve contact.**

**a,** Quantification of 3D morphological features for CD8+ T cells classified as in contact with nerve fibres (“Contact”, *n* = 3,733) or not in contact (“Non-contact”, *n* = 5,838). From top to bottom: prolate ellipticity, cell volume, and oblate ellipticity. Each point represents a single cell; box plots show median and interquartile range with jitters indicating the distribution. Statistical significance is indicated (****).

**Supplementary Figure 11: Prolate ellipticity of T cells in relation to the distance to nerves.**

**a,** Scatter plot of prolate ellipticity versus distance to the nearest nerve across all T cells in the whole volume. The observed data show a negative correlation (*r* = −0.2206), while spatial simulations remain near zero (*r* = 0.0001); shaded area indicates the 98% simulation interval. **b,** Observed Pearson’s r compared with the simulated mean ± SEM, showing that the observed correlation is significantly more negative than expected by simulation (p = 1 × 10⁻⁵).

**Supplementary Figure 12: Vasculature, follicular dendritic cell network and Treg analysis in tertiary lymphoid structures and solitary intestinal lymphoid tissues**

**a**, Relationship between vessel metrics and FDC network volume (CD21 and CD23) across 7 TLS. Left column (blue): total vessel length within the CD21⁺ only (top), CD23⁺ only (middle), and CD21⁺/CD23⁺ union (bottom) compartment plotted against the corresponding compartment volume. Right column (red): total vessel volume within the same three compartments plotted against compartment volume. Solid lines indicate linear least-squares fits, individual TLS are labelled by ID. Pearson r and Spearman ρ with associated p values are shown in each panel (Pearson r = 0.86–0.95, Spearman ρ = 0.82–0.96, all p < 0.05). **b**, Top: Treg to nearest vessel distance (µm) stratified by FDC compartment (CD21⁺, CD23⁺, CD21⁺ and CD23⁺). Boxes show median and interquartile range, individual TLS are overlaid as coloured points. Pairwise comparisons are shown (p = 0.078, 0.109 and 0.688). **c**, per-TLS ratio of Treg counts within the CD21⁺ versus CD23⁺ compartment. The dashed red line marks a ratio of 1 (equal Treg numbers in both compartments), bars above the line indicate more Treg localization in CD21+ compartment. **d**, Relationship between vessel volume and FDC network volume in SILT (n = 4), plotted against CD21⁺ only (left), **e**, CD23⁺ only (middle) and **f**, CD21⁺/CD23⁺ union (right) volume. Pearson r with associated p values is shown in each panel (Pearson r = 0.74–0.85) but did not reach statistical significance given the small sample size (p = 0.150–0.258).

**Supplementary Figure 13: Spatial attraction of FOXP3^+^ cells to CD31^+^ Blood vessels**

**a,** probability density function (PDF) and **b**, cumulative density function (CDF) of the shortest distance to surfaces shown for TLS 2, 3, 4, 5, 6, 9 and 11. The red line indicates the attraction distance. The blue line shows the observed density (percentage of spots) as a function of the shortest distance to surfaces, plotted against the expectation under a complete spatial randomness (CSR) model.

**Supplementary Figure 14: Immune cell phenotype dendrogram**

**a,** Schematic dendrogram of the marker-based workflow used to classify immune cell populations. All immune cells (B cell, T cell, myeloid cell) were first identified and then subdivided into major lineages. **b,** Additional markers used to determine cell states and identify non-immune compartments.

**Supplementary Figure 15: cell-cell membrane interaction analysis**

**a,** membrane interaction graphs between CD11c^+^ IDO1^+^ cell (cyan, IDO1 channel) and Treg (magenta, CTLA-4 channel). **b,** membrane interaction graphs between CD11c^−^ IDO1^+^ cell (cyan, IDO1 channel) and Treg (magenta, CTLA-4 channel).

**Supplementary Figure 16: Workflow decision tree for selecting imaging modalities.**

**a**, Decision tree guiding selection from autofluorescence-based virtual H&E, light-sheet–based volumetric CyCIF for large-volume 3D organization, to re-embedding/sectioning for high-plex high-resolution 3D profiling in selected ROIs, with alternatives for downstream spatial assays listed. Virtual H&E can skip v-CyCIF and go straight to re-embedding, and remaining block after sectioning can also go back to v-H&E and v-CyCIF for further analysis.

## SUPPLEMENTARY TABLES

**Supplementary Table 1:** Sample metadata and related identifiers.

**Supplementary Table 2:** List of tissue clearing types with delipidation.

**Supplementary Table 3:** List of validated Antibodies for v-CyCIF.

**Supplementary Table 4:** Overview of TLS measurements.

**Supplementary Table 5:** Antibody panel for high-plex Lightsheet imaging of normal mucosal SILT.

**Supplementary Table 6:** Antibody panel for LSFM and thick tissue sections used for Re-embedding.

## SUPPLEMENTARY VIDEOS

<u>Supplementary Video 1: High-plex registered LSFM image of cleared normal colonic SILT (DS2).</u>

<u>Supplementary Video 2: Registered LSFM image of nerves in cleared normal colon (DS5).</u>

<u>Supplementary Video 3: High-resolution confocal imaging of a colonic crypt (DS7).</u>

<u>Supplementary Video 4: Fly-through of Tfr-cell density throughout the GC of TLS-2 (DS9).</u>

<u>Supplementary Video 5: Confocal imaging of a TLS in re-embedded 25 µm tissue (DS10).</u>

## MATERIALS AND METHODS

### Human sample collection

All samples were derived from FFPE tissue blocks consented for secondary use. See Supplementary Table 4 for clinical metadata for all specimens. A total of twelve specimens were used in either the main text or supplementary data. Four of these were primary datasets used in the main text, consisting of four normal colons and two CRC blocks (excluding adjacent sections; see supplementary table 1), were obtained from the NIH Cooperative Human Tissue Network (CHTN) with the exception of (DS6 & DS7; LSP22365), retrieved from the archives of the Department of Pathology at Brigham and Women’s Hospital and collected under Harvard Medical School Institutional Review Board approval (Protocol IRB 21-0656) under a waiver of consent. This also includes a melanoma metastasis to the lung and breast tissue. Other samples were sourced from the Cooperative Human Tissue Network (CHTN RRID: SCR-004446).

### Deparaffinization

Formalin-fixed paraffin-embedded (FFPE) tissue blocks were dewaxed at 60 °C for 2 h. In a fume hood, xylene or Envirene (Hardy Diagnostics) was preheated (150–200 mL) to 60 °C either in a beaker covered with aluminium foil on a magnetic hotplate stirrer or in a sealed glass bottle placed in an oven. Tissue samples were placed into tissue cassettes and immersed in preheated xylene with stirring at 60 °C for 24-48 h, with replacement of solution each day. As an alternative, tissues were incubated in sealed xylene-filled bottles at 60 °C with shaking at 100 rpm. The final wash was performed in fresh xylene at 37 °C for 1 h with stirring/shaking at 100 rpm.

Tissues were subsequently transferred to 100% ethanol (EtOH) for 2 × 1 h, then sequentially rehydrated in 70% EtOH, 50% EtOH, and PBS for 1 h each. Samples could be stored in 100% EtOH (up to 1 week), 70% EtOH (several months), or PBS (overnight) before proceeding. For partial sampling, tissue subsections were excised with a scalpel, and the original block was stored in 70% EtOH. If required, 1 mm sections were prepared using a vibratome (Leica VT100s).

### Methanol pretreatment

Bleaching can also help reduce tissue autofluorescence which is an inherent challenge in immunofluorescence staining of tissues. This involved with the inclusion of hydrogen peroxide during methanol pretreatment as described in the iDISCO protocol^29^. Samples were dehydrated sequentially in 50%, 80%, and 100% methanol (MeOH) for 1 h each. An ice-cold pretreatment solution was prepared consisting of 5% hydrogen peroxide in 20% DMSO/MeOH (1.67 mL 30% H₂O₂, 2 mL DMSO, 6.33 mL MeOH per 10 mL total volume). Samples were incubated overnight at 4 °C in this solution. Following incubation, samples were washed in 20% DMSO/MeOH for 1 h, then rehydrated through 100%, 80%, and 50% MeOH, followed by PBS, for 1 h each.

### Antigen retrieval and blocking

Tissues were incubated in a preheated (95–100 °C) 1× Enzo Tris-EDTA antigen retrieval buffer for 1 h. Samples were cooled in the buffer at room temperature for 20 min, rinsed with deionized water, and washed in PBS for 30 min. Tissues were stored in PBS at 4 °C (up to 1 week) prior to subsequent processing. For imaging cycle consistency, tissues were embedded in 8% agarose and sectioned to 1 mm using a vibratome. Samples were washed in PBS (3 × 15 min) on a shaker, followed by blocking in the SuperBlock™ Blocking Buffer at 4 °C.

### 3D sample mounting

The 1 mm tissue sections embedded with agarose were mounted onto 3D printed holders with additional 8% agarose placed between the sample and the frame, ensuring both faces of the tissue remain exposed. This was performed by placing the tissue at the centre of the 3D printed frame on a glass slide with melted agarose poured around the edges carefully. Another glass slide is placed on top to sandwich the tissue and agarose flat. It was then carefully placed at 4 °C to set for 5 min. The top glass slide was then carefully lifted off, and the holder and bottom glass slide was left in PBS to gently separate. The frame was designed using Fusion360 (Autodesk) and snapped into place with an external mount for the MegaSPIM light sheet. For the MuVI SPIM, the sample frame was screwed into the clamp on the sample pole (see supplementary information) .stl files and variations can be found at (https://github.com/Wonga008/CyCIF_holders)

### 3D Autofluorescence based virtual H&E Imaging

After sample mounting, the samples were stained with 50µL of 1mg/mL DAPI (Thermo Fisher Scientific) in 9mL at a final concentration of 15.8 µM in PBS for 24 hours at room temperature. Next, wash 3 x 15 min in PBS and clear in Easy index for 2 h or overnight. Finally, immerse the sample in immersion oil (LifeCanvas Technologies). The sample frame was secured down, DAPI and AF were captured from the 405 nm & 488 nm excitation laser respectively. H&E colouring was performed with simultaneous CyCIF using a modified “falsecolor” Python package (https://github.com/serrob23/falsecolor).

### Fluorophore erasure with photobleaching

To reduce excessive autofluorescence prior to CyCIF, tissues were submerged in a bleaching solution (7.5 mL 30% H₂O₂, 1.5 mL NaOH, 41 mL PBS) in a transparent container for 1 h under white light illumination from both above and below using LED panels. Photobleaching was repeated up to three times during the initial cycle.

### Stochastic electrotransport antibody labelling

Before labelling, samples were incubated for 1 h in the primary sample buffer, and SmartBatch+ staining devices and cups were prepared. Samples were transferred into mesh bags with 3D-printed hooks and submerged in staining cups containing validated antibodies (10 µg per sample; 30 µg if unvalidated for single cups). 20µL of 1mg/mL DAPI was added into the sample cup to achieve a concentration of 6.3µM. For direct immunofluorescence, the SmartBatch+ protocol was followed with appropriate secondary buffers. Samples were washed in PBS (3 × 30 min) on a shaker in the dark and could be stored in the final PBS wash at 4 °C prior to refractive index (RI) matching. For improved performance, Radiant buffer is used instead with a pH catalyst containing resin beads (LifeCanvas Technologies). The voltage was set at 90V with 350 mA current limit.

### Passive Labelling

To rapidly stain smaller samples without specialist equipment, antibody penetration was assisted with C-PRESTO^95^ by combining centripetal force and delayed binding. The tissue was first saturated in SWITCH-OFF^31^ buffer by incubating in PBS containing 0.5 mM SDS for 2 hrs, next the tissue was placed in an Eppendorf or falcon tube and stained with the desired antibodies with a 1 in 50 dilution in SWITCH-OFF buffer at 600 × g in a centrifuge for at least 8 hrs to overnight. Samples were washed with 0.0001% Tween20 (Fischer Scientific) PBST 3 × 30 min at 600 × g on a tabletop centrifuge. Alternatively, samples were stained in a 12 well plate for 24-48 hours at 30°C

### Volumetric CyCIF

Clearing was performed by simple immersion in refractive index (RI) matching solution (RI =1.52). To prepare this solution a 100 mL glass beaker containing a clean stir bar was placed on a hotplate maintained at 37-45 °C. Ultrapure water (3.25 mL) was added, followed by 2g N-Methyl-D-glucamine (Sigma, M2004) and 12g Histodenz (Sigma, D2158). The mixture was stirred until all solids were completely dissolved, after cooling to room temperature, 2.13 mL 2,2′-thiodiethanol (TDE; Sigma) was added slowly under continuous mixing until the solution became homogeneous. The pH was measured with a calibrated pH meter and adjusted to 7.5-8.0 by the dropwise addition of 0.8 M hydrochloric acid (HCl), with thorough mixing after each addition until a stable value was obtained. The solution was then allowed to equilibrate to room temperature, and the refractive index was determined using a digital handheld refractometer (MyBrix Handheld Refractometer, Mettler-Toledo). If the RI was below 1.52, small increments of (Histodenz, Sigma-Aldrich) were added and dissolved completely before re-measurement; if above, small volumes of ultrapure water were added. Alternatively, incubate in EasyIndex RI matching solution (Life Canvas Technologies) overnight at 4 °C or room temperature (minimum 2-3 h). Change to immersion oil prior to imaging. For subsequent imaging cycles, samples were uncleared by washing out EasyIndex in PBS (3 × 15 min) and the process is repeated from the photobleaching step as previously described.

### ECi processing

Samples were sequentially dehydrated in 1-propanol (100%, 70%, 50%, pH 9) before immersion in ECi. If another organic solvent based clearing method is desired, samples are rehydrated back to PBS. Conversion from aqueous based clearing such as EasyIndex to ECi was achieved by PBS washes (3 × 15 min) followed by 1-propanol dehydration and cleared in ECi. The process can also be reversed to convert ECi cleared samples to aqueous based clearing.

### OPTIClear processing

OPTIClear was performed by delipidation of 1 mm thick samples in 4% SDS for 3 days at 55 °C as described in the original protocol. Samples were then washed in PBST (0.1% Triton-X 100) overnight at 37 °C and optionally mounted and in 8% agarose. Refractive index matching solution (RI 1.47) was made with a mixture of Optiprep (Sigma-Aldrich), N-methyl Glucamine and 2,2’ Thioldiethanol to a pH of 7-8 as described in the original protocol.

### 3D Image Acquisition

Cleared tissue samples were imaged on three different light-sheet microscopes. The Muvi-SPIM Lightsheet (Luxendo, Bruker) was equipped with a 405 nm, 488 nm, 561 nm, 642 nm, 685 nm and 785 nm laser lines with a glycerol 10x/0.5 NA immersion objective and dual-sided illumination using two Nikon 10x/0.3 NA water immersion illumination objectives (Nikon). Each channel was acquired using dual sCMOS cameras (Hamamatsu Orca Flash 4.0 v3) with separate optical paths. In some datasets, all channels (excluding 785 nm) were acquired entirely using the “short camera” (405 nm - 642 nm) or only the 405 nm channel with the rest on the “long camera”. Imaging was acquired with 200 ms exposure time for each channel. The 3D printed sample frame has an extruding tab that was secured to a clamp and tightened with a screw and rotated at a 45-degree angle to the objective lens. Due to the restricted size of the chamber, an alternate method was devised using a 3D L-shaped holder that was secured to the pole by a lock and key mechanism with the smaller side of the sample facing the detection lens. However, the angle was more consistent using the sample frame. Images were sampled at 16-bit at 0.65 µm per pixel in X and Y with a step size of 1-2 µm in Z.

Light-sheet imaging was also acquired on a MegaSPIM and DALISPIM open-top inverted microscope (LifeCanvas Technology), equipped with a single camera (Hamamatsu Orca Flash 4.0 v3) and 405 nm, 488 nm, 561 nm and 639 nm laser lines. Samples were imaged with a 9x/0.3 multi-immersion objective lens (Applied scientific instrumentation). 1 mm samples were mounted to the 3D printed sample frame and external bracket which raised the height of the samples and allowed the metal clip from the manufacturer’s original holder to secure the bracket down. This allowed multi-round reproducibility with easy removal and swapping of the sample frames without changing the physical coordinates of the acquisition area, as the manufacturer’s holders are secured down by magnets. The oil bath is manually placed on the stage by eye, and it is not moved in subsequent imaging rounds. Images were sampled at 16-bit at 0.7 µm per pixel in X, Y and Z.

Other cleared tissue imaging was performed on a CellVoyager CQ1 benchtop spinning disk confocal microscope (Yokogawa) equipped with 405 nm 488 nm 561 nm and 640 nm laser lines, and 10×/0.4 NA air objective (Olympus) and a LSM 980 confocal laser scanning microscope (Carl Zeiss) equipped with a 405nm, 488nm, 561nm, 647nm, and 750nm laser lines, and 5×/0.16 NA air, 10×/0.45 NA air and 40×/1.3 NA oil immersion objective lenses.

### 3D Image processing and registration

Deskewing and stitching of z-blocks from datasets acquired on the MegaSPIM and DaliSPIM were performed on Lifecanvas Technologies post-processing software. Tiles from MuVI-SPIM datasets were stitched and registered using the LuxBundle software v5.1.0-beta.1 (Luxendo, Bruker). Registration was performed by using DAPI from the first cycle as the reference channel. For subsequent cycles, DAPI served as the moving image, the warped transformation generated was then applied to the other channels in the moving image’s cycle. The process was repeated for each cycle as the new moving image. Automatic 3D non-linear registration was performed on a ROI from a 4× downsampled volume using Advanced Normalization Tools (ANTs) via python code or as a 3D slicer extension. Semi-automatic methods include a custom MATLAB script from previous works that utilises MATLAB’s imregdemon’s function^96^. Full resolution registration was also performed manually using the Fiji (ImageJ) plugin BigWarp^97^ from .xml BDV and Imaris files.

### Re-embedding of cleared tissues into paraffin sections

To reimage the cleared tissue as paraffin sections on glass slides, the sample is washed in PBS, photobleached and stored in PBS. The surrounding agarose is carefully removed and embedded with HistoGel (Epredia) into a block, with excess removed. The sample is then dehydrated in 70%, 90%, and two changes of 100% ethanol for 15 min each before finishing with another two changes of 100% ethanol for 30 min and 45 min respectively. Next the tissue is cleared with three changes of xylene, twice for 20 min each with a final change for 45 min. The tissue can now be infiltrated with paraffin wax, first melt the pellets into a liquid (polyfin embedding medium, VWR) at 60°C, the sample (no greater than 4 mm thick) was then immersed in the molten paraffin with two changes at 30 min and a third change at 45 min, before setting in a cassette using an embedding station (Tissue Embedding Centre, Electron Microscopy Sciences). Finally, the block was sectioned at a thickness of 25 μm using a microtome and mounted on SuperFrost Plus glass slides (Fisher Scientific). 3D CyCIF was then performed on the confocal.

### 3D Segmentation of virtual H&E

Foundation models trained on histology images were used to extract morphological features from virtual H&E images for segmentation of normal glands, glandular tumour, and SILT from 3D datasets. Foundation models have been trained on more data than classical machine learning methods, such as ilastik, and therefore, exhibits superior accuracy. To reduce workload and mitigate errors introduced by manually delineating boundaries of these tissue structures, a sparse autoencoder (SAE) approach was applied to automatically identify and refine morphology features. We took all xy-planes of the 3D vH&E dataset as whole slide images, removed background regions, and tiled tissue regions into overlapping patches to be embedded by the UNI foundation model (FM)^55^. An SAE was then trained to sparsely reconstruct FM embeddings, and the resulting SAE features were reviewed by a pathologist to select for ones that correspond to tissue structures of interest. Token-level FM embeddings from the top scoring patches for selected features are then used to refine patch-level feature prediction to pixel-level by training a U-Net to predict the token cluster from vH&E patches. Finally, the U-Net was applied to 2D planes of the 3D dataset to segment the tissue structure of interest.

### Cell count estimation

Colon images were downsampled 4–8×, and Imaris machine-learning segmentation was used to generate surface meshes for mucosa, submucosa, muscularis, serosa, and SILT compartments. For each compartment, cell density was calculated from CellposeSAM-segmented representative 3D crops and multiplied by the corresponding Imaris-derived compartment volume to estimate total cell number. SILTs were segmented separately, excluded from the adjacent mucosal volume and cell estimate, and added back separately by applying the SILT cell density to the measured SILT volumes and summing across all SILTs. Estimated cell numbers from all compartments were then summed to obtain the final total colon cell estimate.

### Correlation of cell prolateness and distance to nerve

Cell-level distance-to-nerve measurements were paired with ellipticity/prolateness measurements from the same cells. Cells with missing or nonnumeric values for either measurement were excluded. Pearson correlation was used to quantify the linear association between distance to nerve and cell prolateness. Uncertainty was estimated using paired nonparametric bootstrapping. For each bootstrap iteration, 1,436 paired cell observations were sampled with replacement from the observed dataset, preserving each cell’s distance/prolateness pairing. Pearson’s r and the least-squares regression line were recalculated for each resample. This was repeated 10,000 times. The 95% confidence interval for Pearson’s r was calculated from the 2.5th and 97.5th percentiles of the bootstrapped r distribution. The regression confidence band was calculated pointwise from the bootstrapped regression predictions. A two-dimensional kernel density estimate was plotted to show the observed density of cells in distance/prolateness space.

### Quantification of spatial distance and spot accumulation analysis

3D spatial distances were calculated using distance transformation XTension within Imaris v10.0.1 to calculate the shortest distance from a channel of interest to a surface object. Depending on the image, cells were segmented by intensity thresholding or importing a segmentation mask to create surfaces. For spot accumulation analysis cell surfaces were first converted into spots, positions of cell surfaces were exported as .csv with headers removed containing only the x,y,z positions. Next, Create Spots from CSV.m (https://github.com/BIDCatUCSF/Create-Spots-From-Text) was executed in Imaris as an Xtension. Object-object statistics were enabled in the surface and spots of interest, the tabulated statistics for probability density and cumulative density function were obtained in the “Vantage” and “Spatial View” tab within Imaris and finally exported as .csv. The observed density and complete spatial random data were plotted using matplotlib python package. The attraction distance was defined as the point where the observed and CSR probability density functions intersect, and this value was overlaid on the cumulative density function plot.

### 3D morphology analysis of surface meshes

Cells were segmented with the u-Segment3D Python package^42^(https://github.com/DanuserLab/u-segment3D). The resultant cell masks were then converted into individual .obj surface mesh using a custom script utilizing the u-Unwrap^98^ Python package (https://github.com/DanuserLab/u-unwrap3D). These meshes were finally converted into point clouds as. ply and normalized to one unit sphere. The folder of point clouds was inputted into a dynamic graph convolutional neural network using the cellshape package^63^ (https://github.com/Sentinal4D/cellshape) trained on ShapeNet (https://huggingface.co/datasets/ShapeNet/ShapeNetCore) to visualize the latent space. Basic morphology metrics such as volume, sphericity and ellipticity were measured in Imaris 10.0.1.

### 3D spatial analysis of vasculature and T follicular regulatory (Tfr) cells in follicular dendritic cell (FDC) regions

Three-dimensional lightsheet fluorescence images of CD21, CD23, CD31, and FOXP3 from the DALISPIM were analysed in MATLAB (2024b). First, CD21, CD23, CD31, and FOXP3 channels were extracted from Imaris .ims volumes and downsampled by 2-fold across all axes to reduce computational requirements. Pixel-classification models for CD21, CD23, a normalized summed channel of CD21 and CD23, and CD31 were trained in Ilastik to generate probability maps. The summed CD21/CD23 probability map was thresholded at 0.5 to identify candidate FDC regions, followed by removal of small connected components.

For each candidate FDC region, a 70-voxel 3D buffer was applied. CD21 and CD23 probability maps were independently thresholded and morphologically processed to generate CD21- and CD23-positive masks. The FDC mask was defined as the union of these masks, and CD21, CD23, and total FDC volumes were quantified.

The vasculature was segmented using fibermetric, a standard MATLAB function, on the CD31 probability maps. These then underwent skeletonization, and vessel volume and area normalized to the CD21, CD23, and total FDC compartments. FOXP3-positive Tfrs were detected using Laplacian-of-Gaussian filtering and adaptive intensity thresholding. Nuclear counts were calculated within each compartment, and a three-dimensional distance transform was used to determine the mean distance of FOXP3-positive nuclei from the nearest blood vessel.

### Statistics

Statistical analyses were performed in Python (SciPy). Differences between direct-contact and non-contact cells were assessed using a two-sided Mann–Whitney U test, as prolate ellipticity values were non-normally distributed. Data are presented as individual data points with median and interquartile range indicated, and significance thresholds defined as *p < 0.05, **p < 0.01, ***p < 0.001 and ****p < 0.0001. All measurements were derived from a single 3D tissue volume, with n values indicating the number of cells analysed per group.

## ACKNOWLEDGEMENTS

This work was primarily supported by an Aspire award from the Mark Foundation for Cancer Research, by the Ludwig Center at Harvard, and by CCBIR grant U54-CA268072. Instrumentation was supported by Bits to Bytes Grant 44038 from the Massachusetts Life Sciences Center. A.Y.H.W. was supported in part by NIH Common Fund 3OT2OD033759-01S3; Z.M. was supported by Research Specialist Award R50-CA252138 and C.Y. by R50-CA305002. Histopathology was supported by P30-CA06516. S.S. is supported by the BWH President’s Scholars Award. Development of computational methods and image processing software is supported in part by a Team Science Grant from the Gray Foundation. We thank Luxendo (a Bruker company) for the generous loan of a Muvi-SPIM microscope and the Aquifer HIVE server used in this study. We also thank S. R. Talemi, J. Lin, Y. Chen, J. Muhlich, G. Baker, T. Vallius, S. Pant and J. Tefft for the helpful discussions. P. Llopis and team at MicRoN core facility for assistance with microscopy, and N. Girnius for materials. We are grateful to LifeCanvas Technologies, Alpenglow Biosciences and Carl Zeiss Microscopy for additional support with microscopy.

## AUTHOR CONTRIBUTION STATEMENTS

A.Y.H.W., Y.D.L., C.Y., and P.K.S. conceived and designed the study. G.D., S.S., C.Y. and P.K.S. supervised the work and acquired funding. Z.M., S.C., and S.S. assisted with biological interpretation. S.S. helped with human clinical specimens and oversaw histopathology annotations. A.Y.H.W., Y.D.L., C.Y. and Y.N.A.A. acquired the data and A.Y.H.W., F.Z., C.Y., Y.D.L. and S.W.C. performed data analysis. A.Y.H.W., Y.D.L., H.P., C.Y., and P.K.S. wrote the manuscript with input from all authors.

## Data Availability

All primary images and derived data (∼15 TB) will be available via AWS transfer at the time of publication. Instructions for accessing the primary and derived data is available via a data index page on Zenodo (https://doi.org/10.5281/zenodo.20170967). These images can be viewed using the free ImarisViewer (https://imaris.oxinst.com/imaris-viewer). A subset of data is available for 3D interactive viewing within the browser-based tool as 3D Gaussian Splats (see Github repo), including augmented/virtual reality within Vitessce (http://vitessce.io/)^99^ or in The Human Reference Atlas (HRA) Organ Gallery^100^. This effort is a work in progress and will be available in the future.

## Code Availability

Original code associated with this paper is available on GitHub and Zenodo at the time of paper publication.

