## Supplementary Information for "Volumetric Cyclic Immunofluorescence for 3D Spatial Profiling of Tumour, Stroma, and Immune Structures in Human FFPE Tissue"

### Content


| Title | Page |
|---|---|
| Supplementary Figure 1: Macroscopic inspection of FFPE tissues during sample preparation for v-CyCIF. | 2 |
| Supplementary Figure 2: Macroscopic assessment of colon tissue clearing. | 3 |
| Supplementary Figure 3: Representative v-CyCIF optical planes across imaging cycles. | 4 |
| Supplementary Figure 4: Cell segmentation across colonic compartments and immune aggregates. | 5 |
| Supplementary Figure 5: Comparison of antibody staining specificity across fluorophores. | 6 |
| Supplementary Figure 6: Orthogonal Views before and after registration of the centre slice in cycle 1-8. | 7 |
| Supplementary Figure 7: Orthogonal Views before and after registration of the centre slice in cycle 9-17. | 8 |
| Supplementary Figure 8: Nerve fibre diameter and orientation analysis. | 9 |
| Supplementary Figure 9: Spatial analysis of CD8 T-cell morphology in relation to nerve proximity. | 10 |
| Supplementary Figure 10: Cell shape and size differences associated with cell–nerve contact. | 11 |
| Supplementary Figure 11: Prolate ellipticity of T cells in relation to the distance to nerves. | 12 |
| Supplementary Figure 12: Vasculature, follicular dendritic cell network and Treg analysis in tertiary lymphoid structures and solitary intestinal lymphoid tissues. | 13 |
| Supplementary Figure 13: Spatial attraction of FOXP3+ cells to CD31+ Blood vessels. | 15 |
| Supplementary Figure 14: Immune cell phenotype dendrogram. | 16 |
| Supplementary Figure 15: cell-cell membrane interaction analysis. | 17 |
| Supplementary Figure 16: Workflow decision tree for selecting imaging modalities. | 18 |
| Supplementary note | 20 |

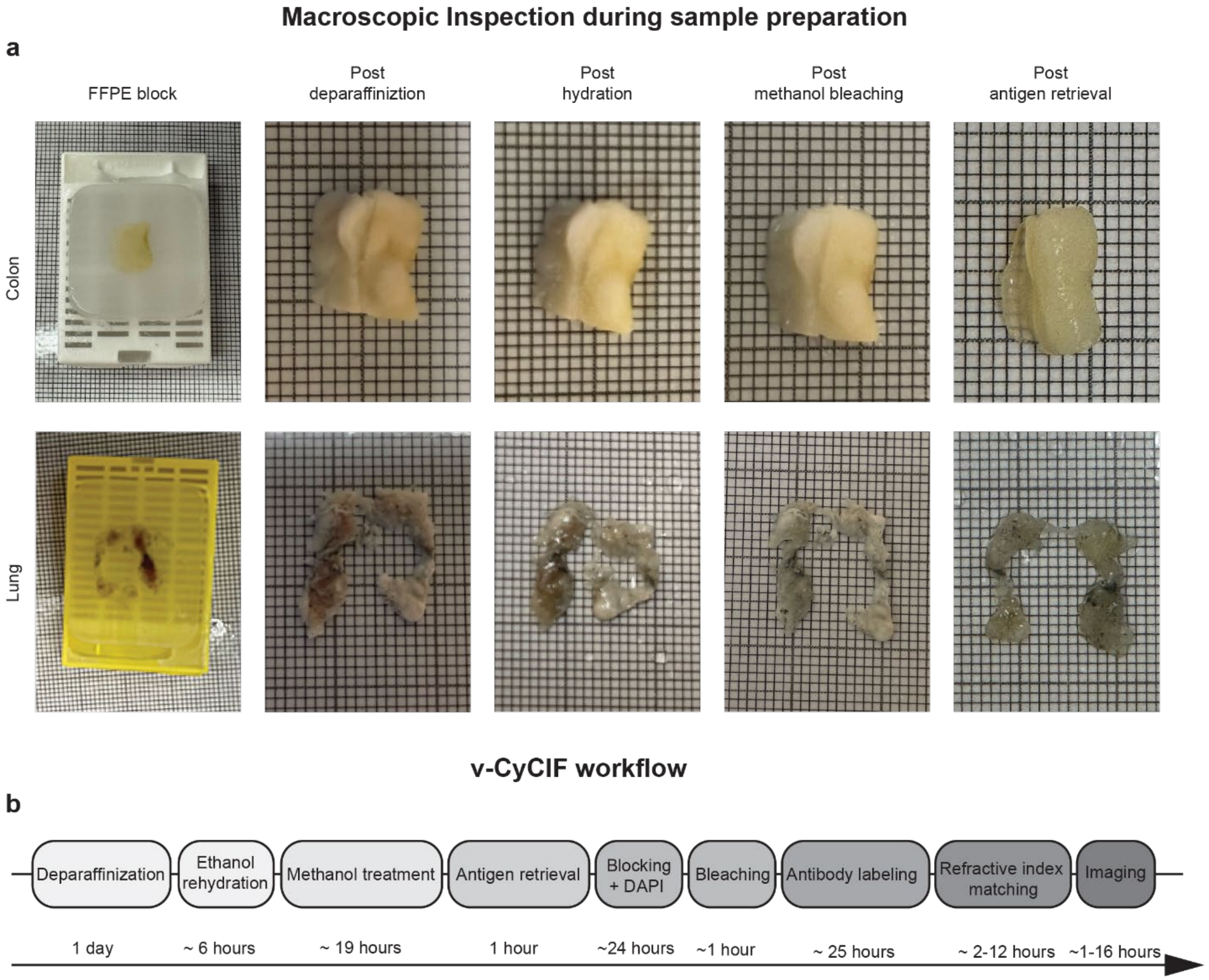


**Supplementary Figure 1: Macroscopic inspection of FFPE tissues during sample preparation for v-CyCIF.**

**a,** Representative photographs of FFPE human colon (top row) and lung (bottom row) specimens at key processing steps. Images show the tissue in the FFPE block, followed by appearance after deparaffinization, after hydration, after methanol bleaching, and after antigen retrieval. **b,** Workflow of v-CyCIF and estimated time for each processing step.

**Tissue transparency pre- and post-clearning**

**a**

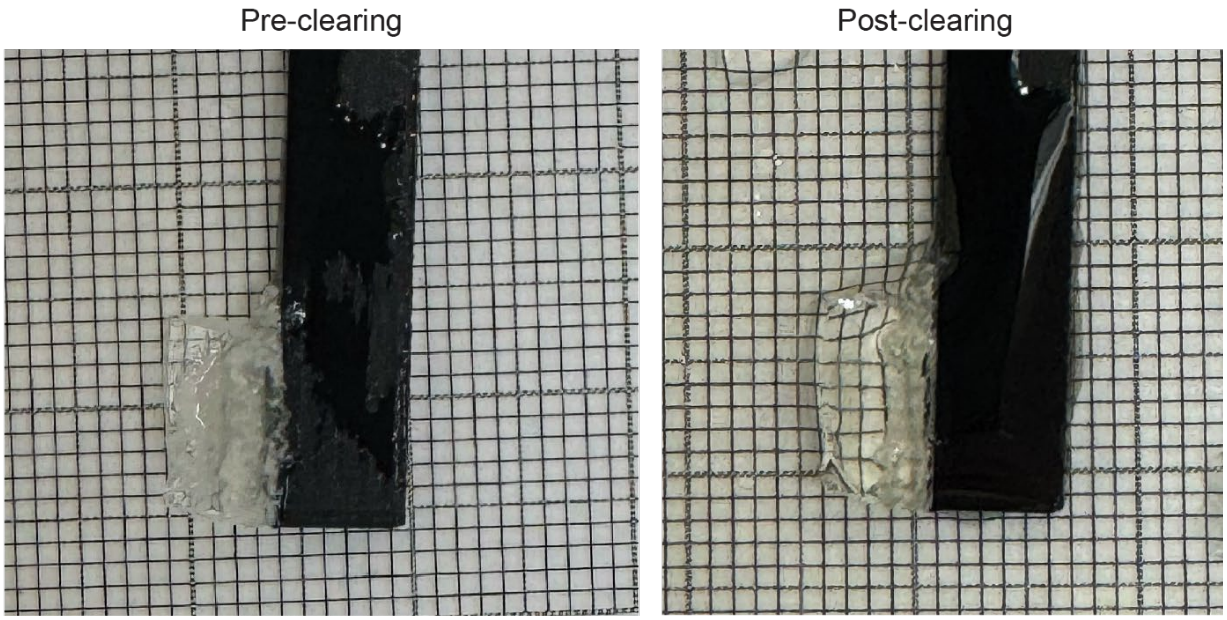


**Supplementary Figure 2: Macroscopic assessment of colon tissue clearing.**

**a,** Representative photographs of colon tissue before (left) and after (right) treatment with a refractive index–matching solution (RI = 1.52). Increased visibility of the background grid after treatment indicates enhanced tissue transparency following clearing.

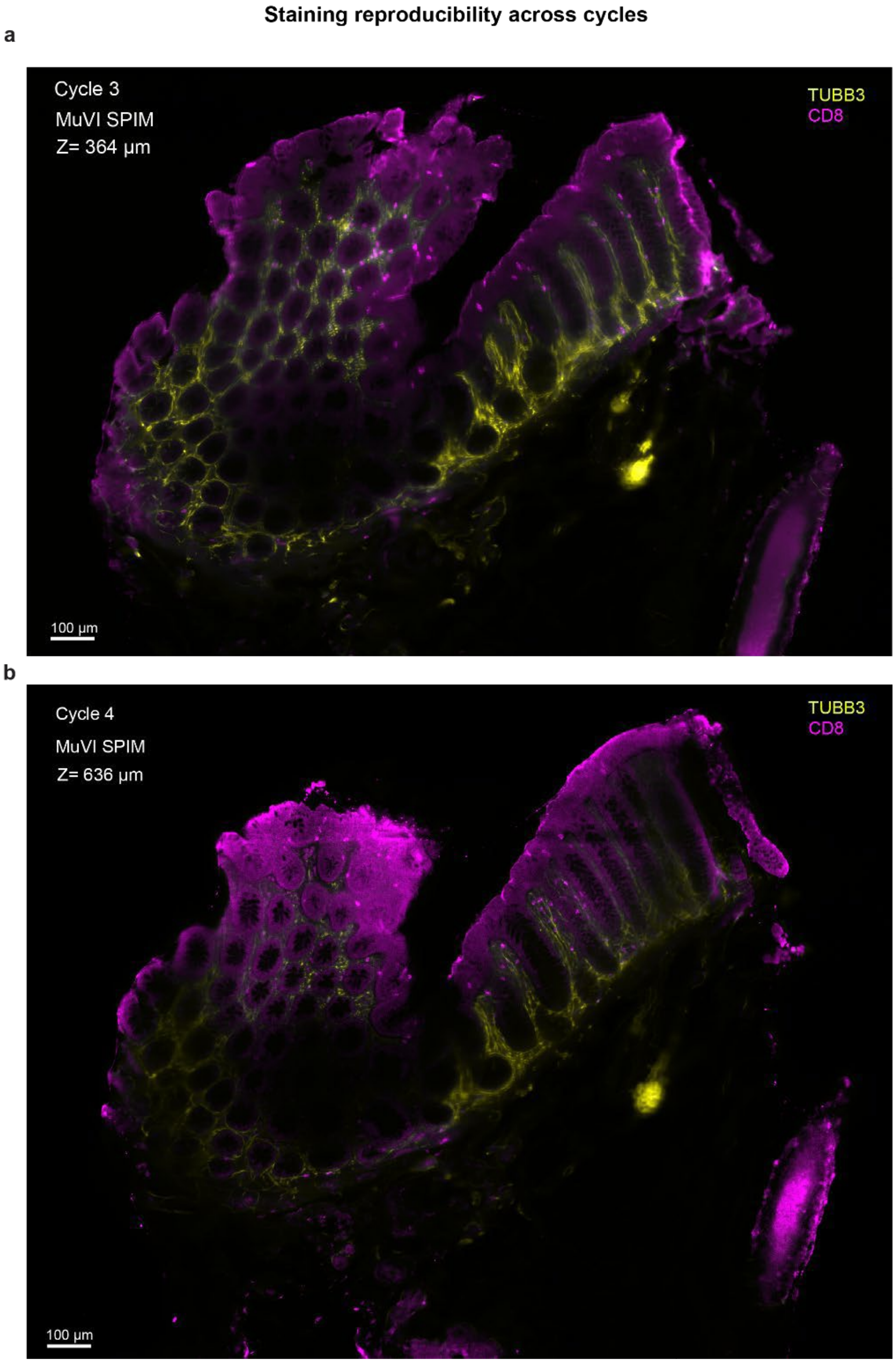


**Supplementary Figure 3: Representative v-CyCIF optical planes across imaging cycles.** **a,** Single optical plane (z = 364 µm) from a colonic region acquired by MuVI-SPIM in cycle 3. **b,** Single optical plane (z = 636 µm) from the same colonic region in (**a**) acquired by MuVI-SPIM in cycle 4. Neuronal structures are labelled by TUBB3 (yellow) and $CD8^+$ T cells by CD8 (magenta). Scale bars: 100 µm.

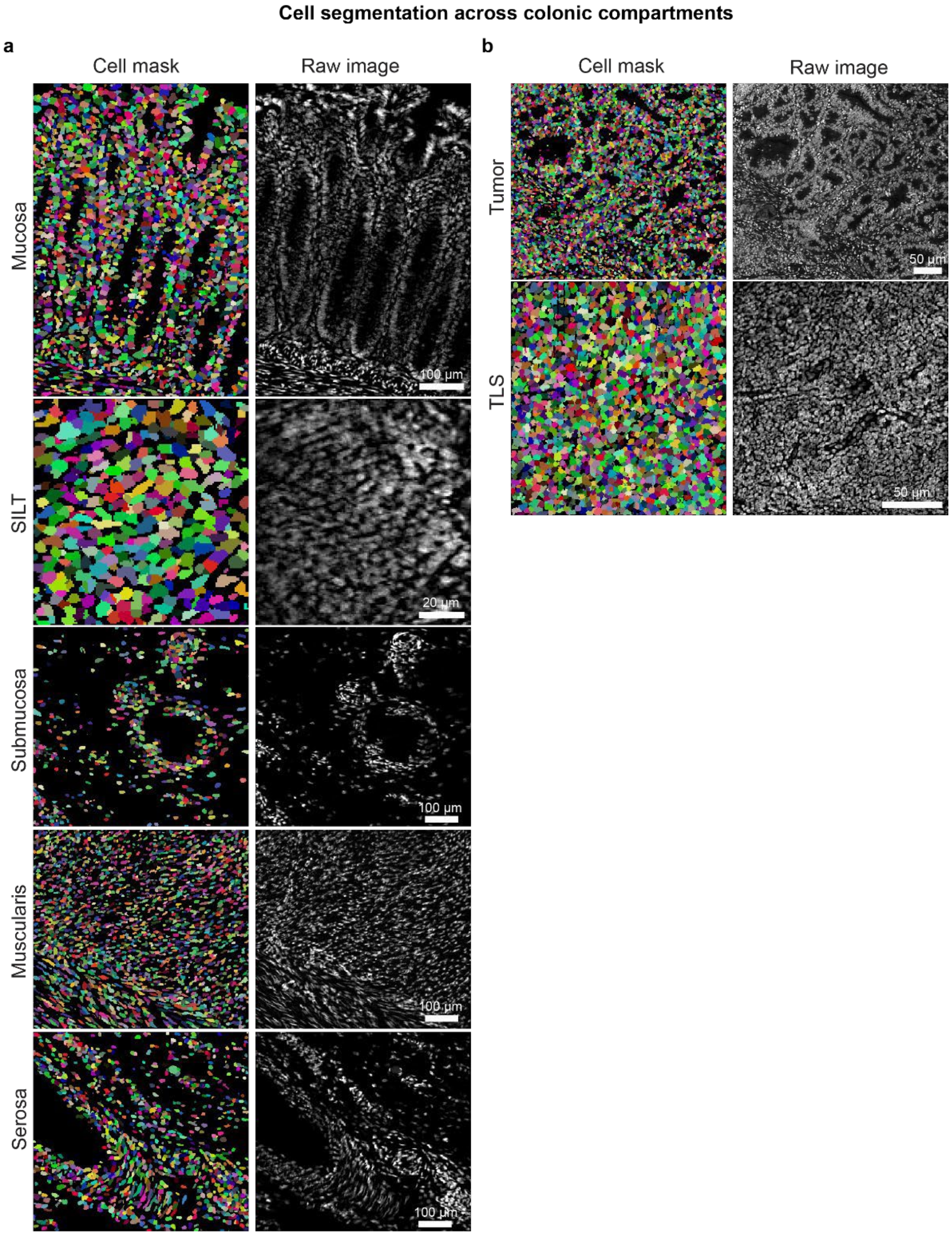


**Supplementary Figure 4: Cell segmentation across colonic compartments and immune aggregates.**

**a,** Example cell segmentation masks (left; pseudo-coloured by object) and corresponding raw DAPI signal (right) for different compartments (mucosa, SILT, submucosa, muscularis, serosa) in the normal colon. Scale bar: 100 µm (Scale bar: 20 µm for SILT). **b,** Example cell segmentation masks (left; pseudo-coloured by object) and corresponding raw DAPI signal (right) for different compartments in the tumour. Scale bar: 50 µm

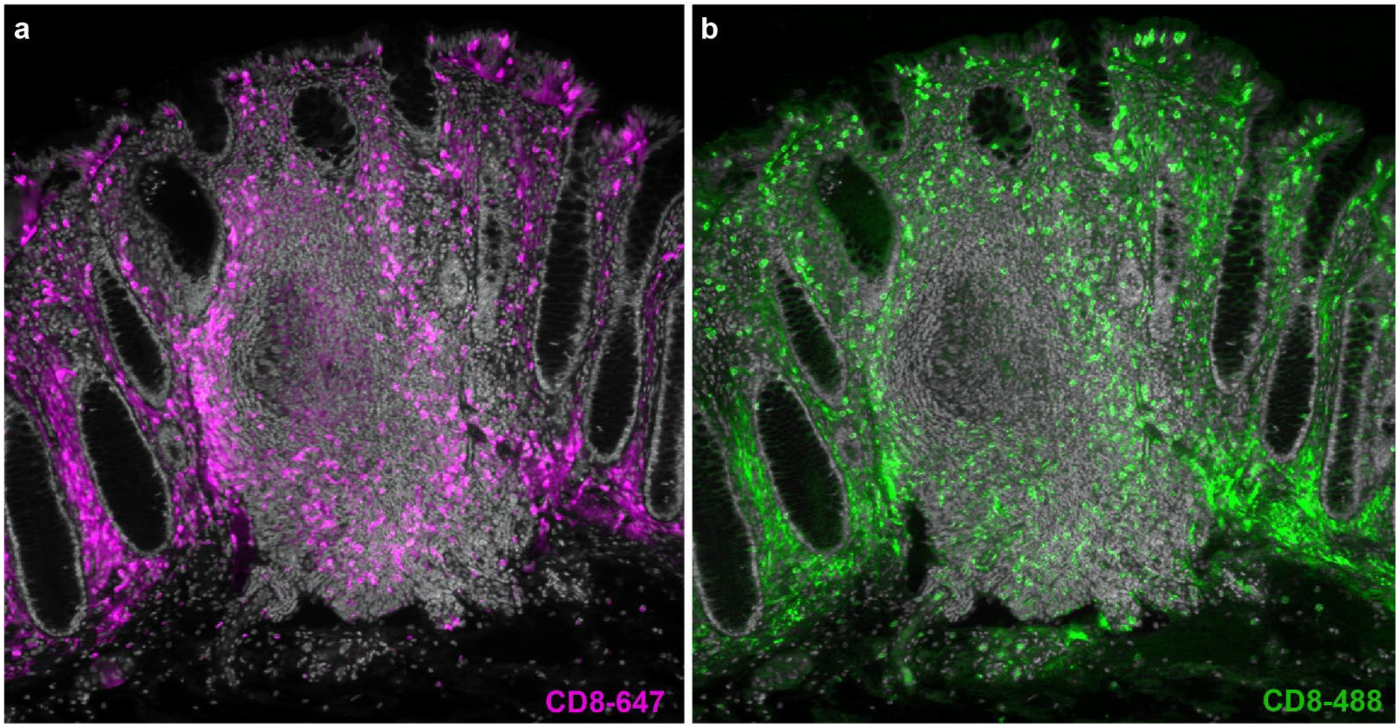


**Supplementary Figure 5: Comparison of antibody staining specificity across fluorophores**

**a-b**, Single-channel optical sections from the volume in (**Fig. 3a**) showing CD8 conjugated to Alexa fluor 647 (**a**, magenta) and Alexa fluor 488(**b**, green) distribution relative to the SILT.

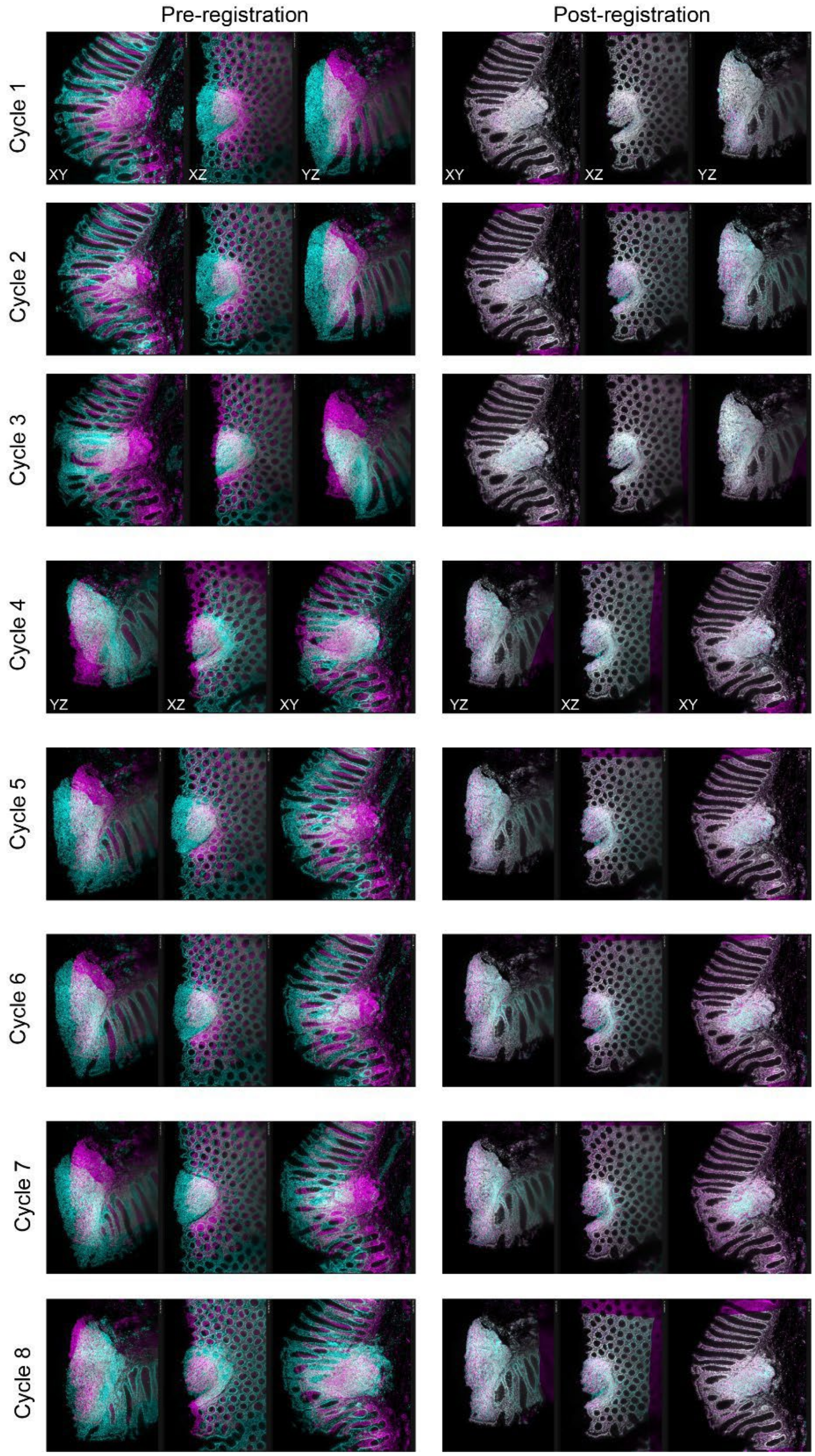


**Supplementary Figure 6: Orthogonal Views before and after registration of the centre slice in cycle 1-8.**

**a**, Quality control of centre slices from XY, YZ and XZ views showing before and after ANTs registration from image cycles 1-8.

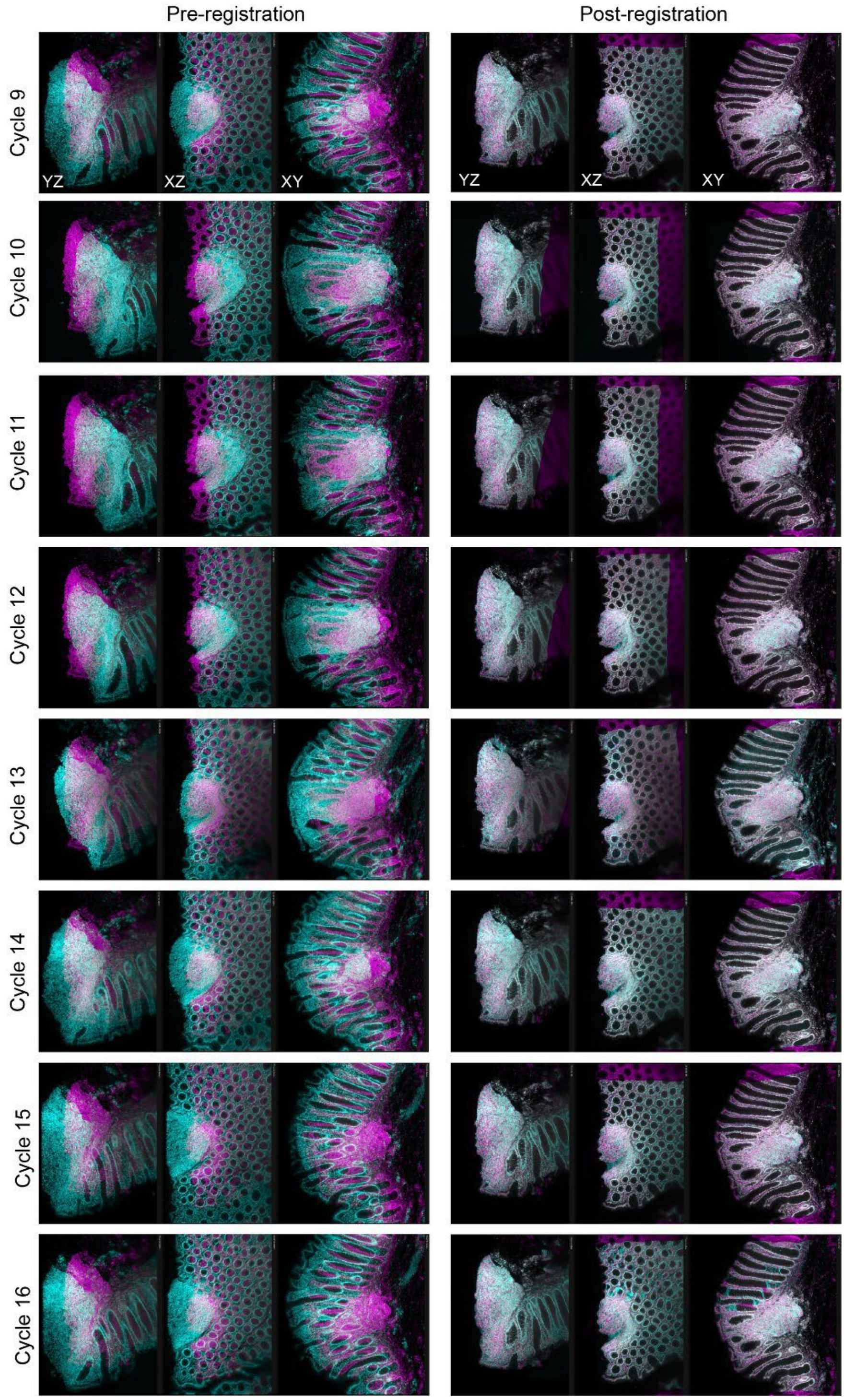


**Supplementary Figure 7: Orthogonal Views before and after registration of the centre slice in cycle 9-17.**

**a,** Quality control of centre slices from XY, YZ and XZ views showing before and after ANTs registration from image cycles 9-17.

#### Nerve fiber and orientation analysis

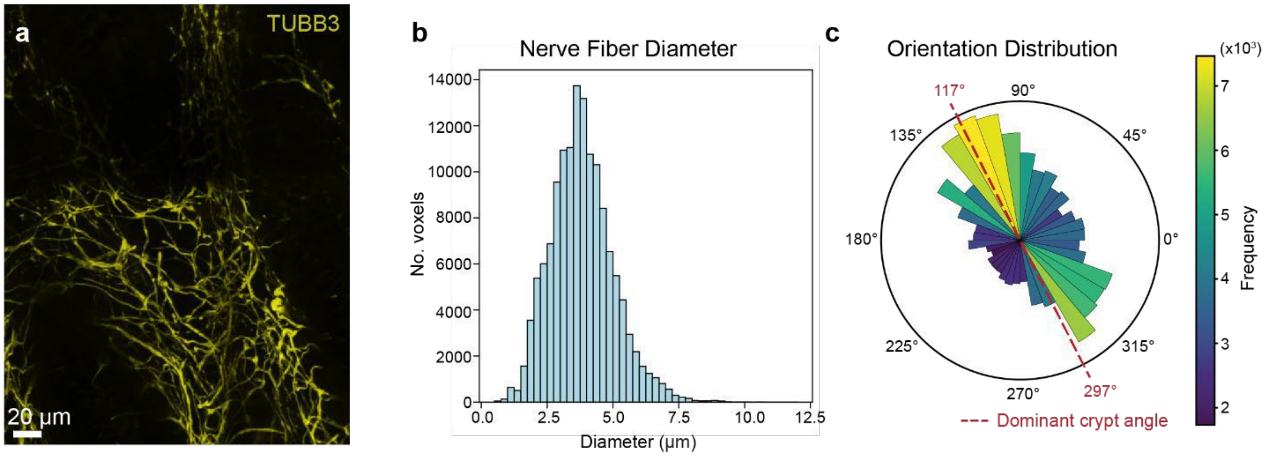


**Supplementary Figure 8: Nerve fibre diameter and orientation analysis.**

**a,** Zoomed-in view of the TUBB3⁺ nerve fibre from (**Figure 6**) used for fibre analysis in (**b,c**). Scale bar: 20 µm. **b,** Distribution of TUBB3⁺ nerve fibre diameters from the analysed volume (n = 140,490 fibres). **c,** Polar histogram of TUBB3⁺ nerve fibre orientation distribution (colour indicates binned frequency). Red dashed line indicates dominant crypt orientation (117° and 297°).

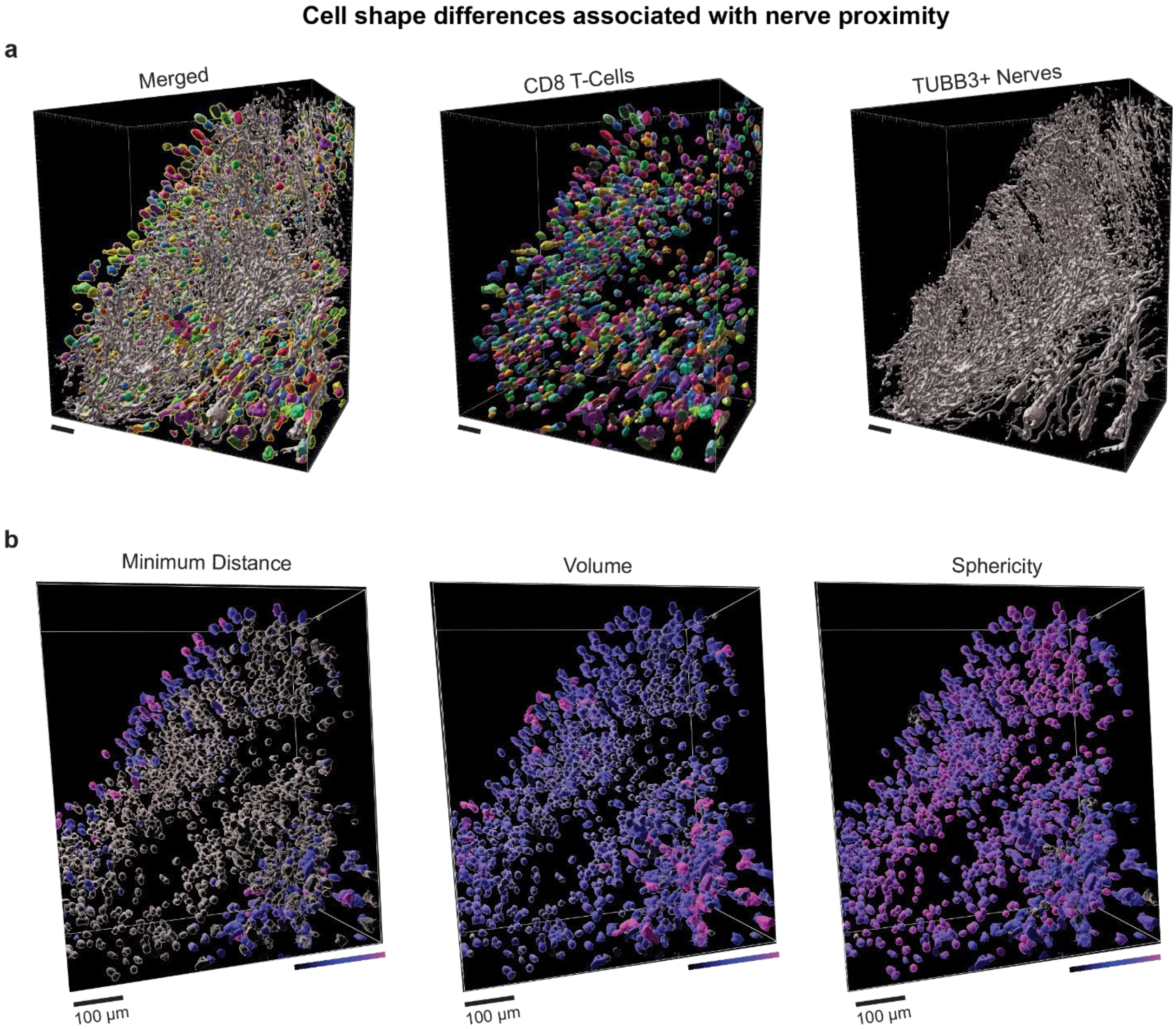


**Supplementary Figure 9: Spatial analysis of CD8 T-cell morphology in relation to nerve proximity.**

**a,** Surface meshes for individual T cells (colour coded, middle), nerve fibres (grey, right) and merged (left) from the zoomed-in ROI in (**Figure 6c**). Scale bar: 70 μm. **b,** $CD8^+$ T cells colour coded by morphological and spatial attributes, including minimum distance to nerves, cell volume, and shape parameters. Scale bar: 100 μm.

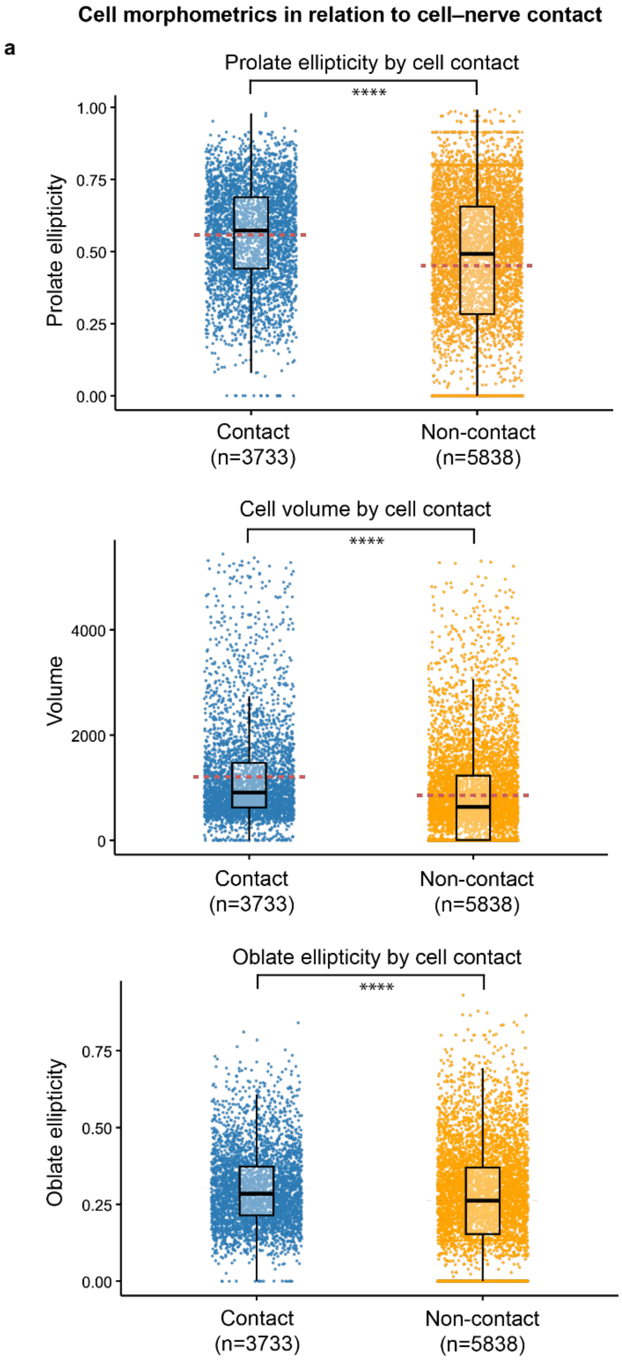


**Supplementary Figure 10: Cell shape and size differences associated with cell–nerve contact.**

**a,** Quantification of 3D morphological features for $CD8^+$ T cells classified as in contact with nerve fibres ("Contact", *n* = 3,733) or not in contact ("Non-contact", *n* = 5,838). From top to bottom: prolate ellipticity, cell volume, and oblate ellipticity. Each point represents a single cell; box plots show median and interquartile range with jitters indicating the distribution. Statistical significance is indicated (****).

a

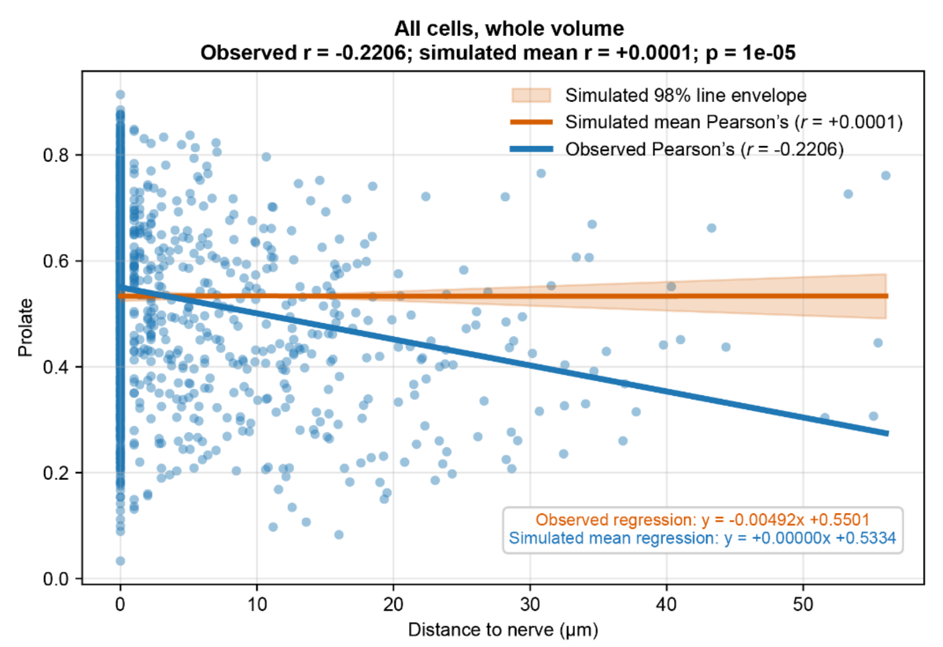


b

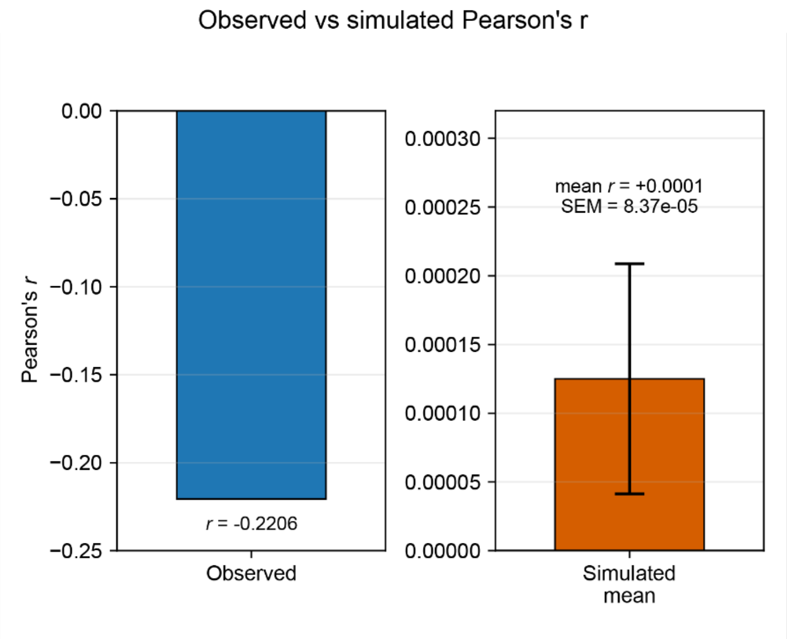


**Supplementary Figure 11: Prolate ellipticity of T cells in relation to the distance to nerves.**

**a,** Scatter plot of prolate ellipticity versus distance to the nearest nerve across all T cells in the whole volume. The observed data show a negative correlation (r = −0.2206), while spatial simulations remain near zero (r = 0.0001); shaded area indicates the 98% simulation interval.

**b,** Observed Pearson's r compared with the simulated mean ± SEM, showing that the observed correlation is significantly more negative than expected by simulation ($p = 1 \times 10^{-5}$).

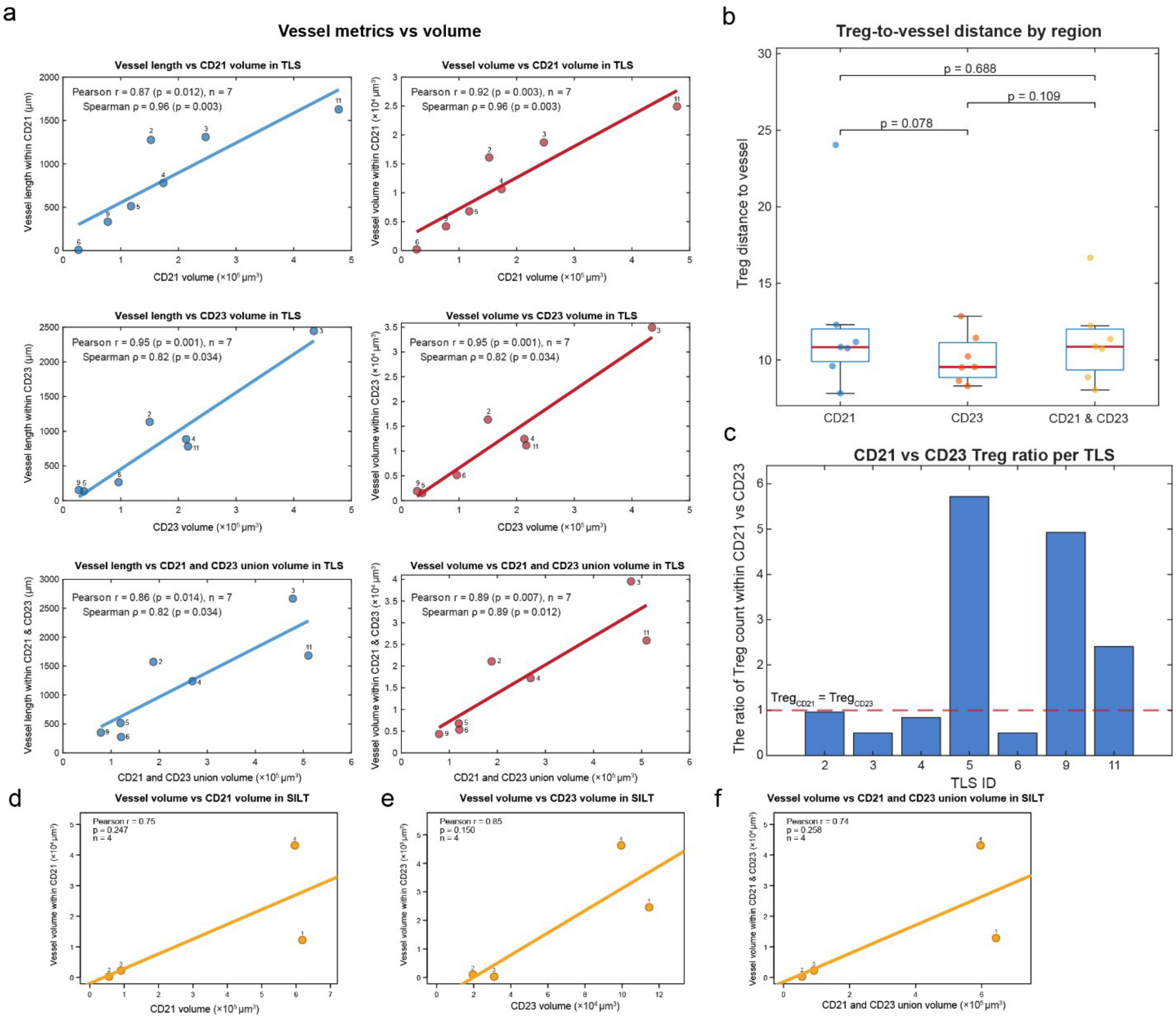


**Supplementary Figure 12: Vasculature, follicular dendritic cell network and Treg analysis in tertiary lymphoid structures and solitary intestinal lymphoid tissues**

**a**, Relationship between vessel metrics and FDC network volume (CD21 and CD23) across 7 TLS. Left column (blue): total vessel length within the $CD21^+$ only (top), $CD23^+$ only (middle), and $CD21^+/CD23^+$ union (bottom) compartment plotted against the corresponding compartment volume. Right column (red): total vessel volume within the same three compartments plotted against compartment volume. Solid lines indicate linear least-squares fits, individual TLS are labelled by ID. Pearson r and Spearman ρ with associated p values are shown in each panel (Pearson r = 0.86–0.95, Spearman ρ = 0.82–0.96, all $p < 0.05$). **b**, Top: Treg to nearest vessel distance (μm) stratified by FDC compartment ($CD21^+$, $CD23^+$, $CD21^+$ and $CD23^+$). Boxes show median and interquartile range, individual TLS are overlaid as coloured points. Pairwise comparisons are shown (p = 0.078, 0.109 and 0.688). **c**, per-TLS ratio of Treg counts within the

CD21$^+$ versus CD23$^+$ compartment. The dashed red line marks a ratio of 1 (equal Treg numbers in both compartments), bars above the line indicate more Treg localization in CD21+ compartment. **d**, Relationship between vessel volume and FDC network volume in SILT (n = 4), plotted against CD21$^+$ only (left), **e**, CD23$^+$ only (middle) and **f**, CD21$^+$/CD23$^+$ union (right) volume. Pearson r with associated p values is shown in each panel (Pearson r = 0.74–0.85) but did not reach statistical significance given the small sample size (p = 0.150–0.258).

a

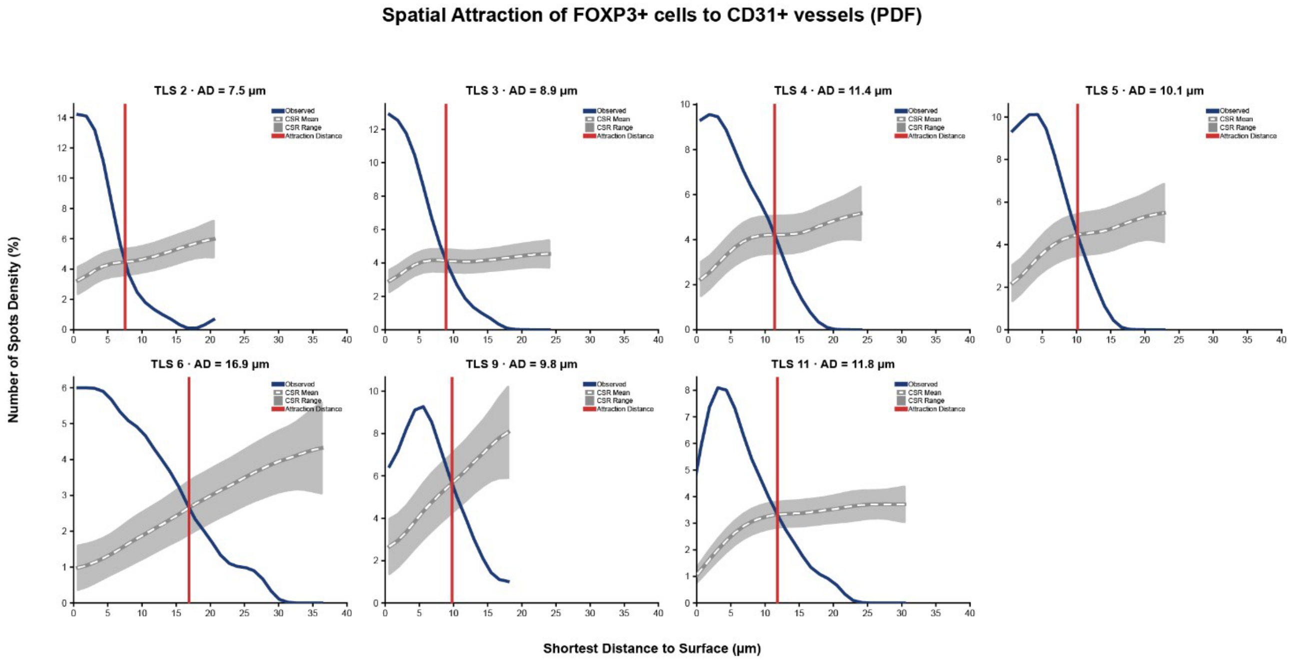


b

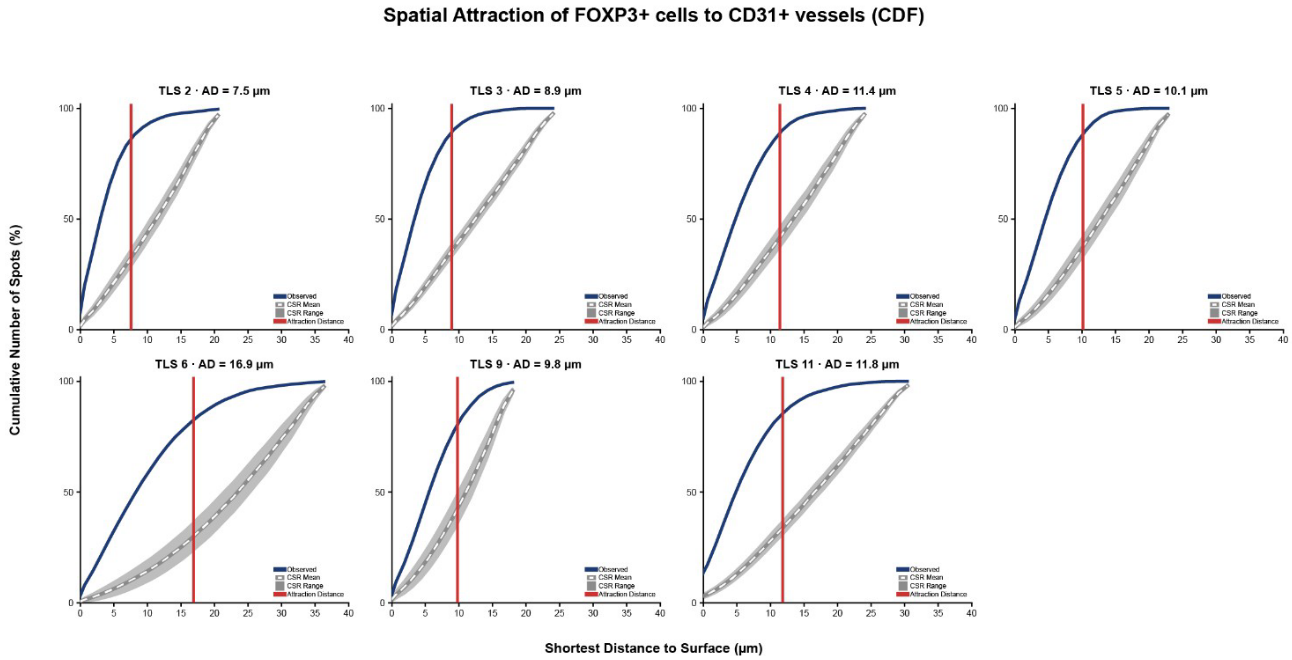


**Supplementary Figure 13: Spatial attraction of FOXP3$^+$ cells to CD31$^+$ Blood vessels**

**a,** probability density function (PDF) and **b**, cumulative density function (CDF) of the shortest distance to surfaces shown for TLS 2, 3, 4, 5, 6, 9 and 11. The red line indicates the attraction distance. The blue line shows the observed density (percentage of spots) as a function of the shortest distance to surfaces, plotted against the expectation under a complete spatial randomness (CSR) model.

#### Immune cell phenotype dendrogram

**a**

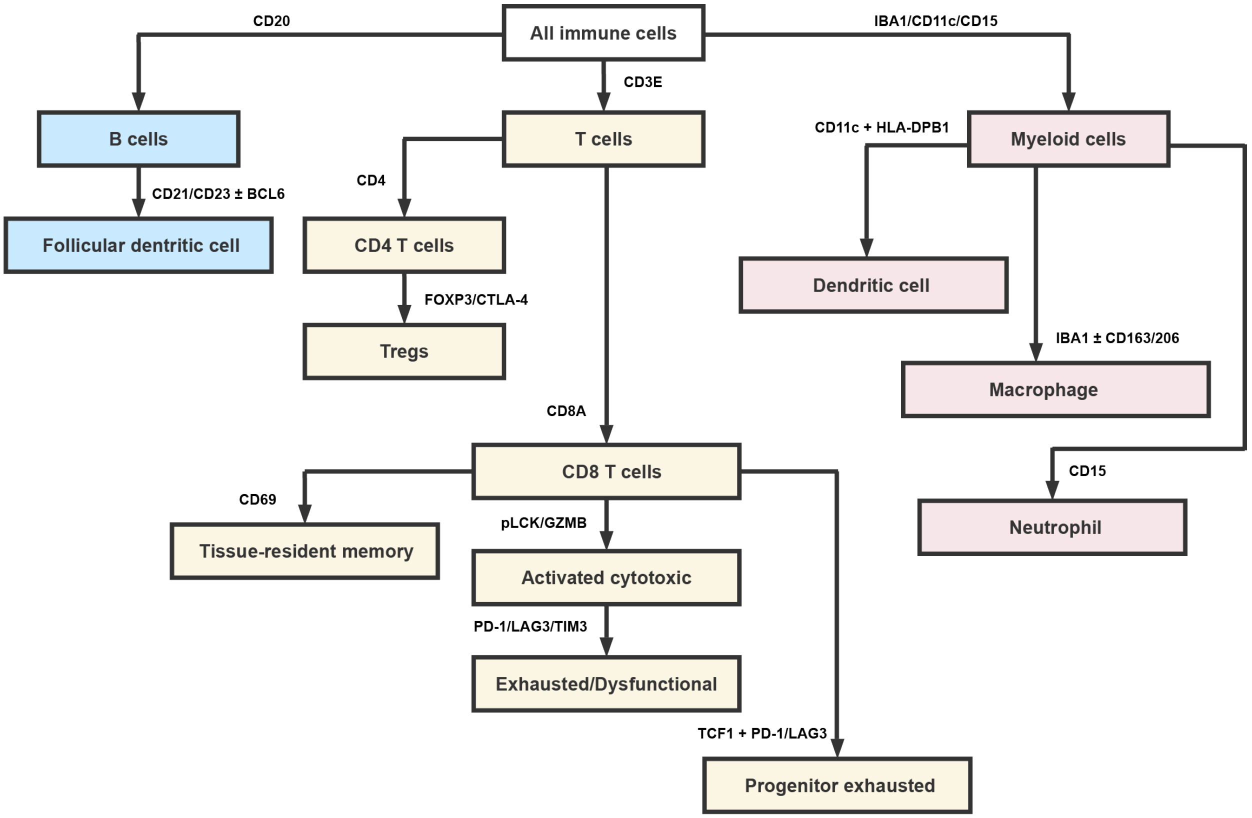


Additional markers used to determine cell states and non-immune cells

| | | | |
|---|---|---|---|
| Vimentin | CDX2 | Connexin43 | β-actin |
| PanCK | Ki67 | TUBB3 | MX1 |
| E-cadherin | IDO1 | PDPN | α-SMA |

**Supplementary Figure 14: Immune cell phenotype dendrogram.**

**a,** Schematic dendrogram of the marker-based workflow used to classify immune cell populations. All immune cells (B cell, T cell, myeloid cell) were first identified and then subdivided into major lineages. **b,** Additional markers used to determine cell states and identify non-immune compartments.

#### Cell–cell membrane interaction analysis

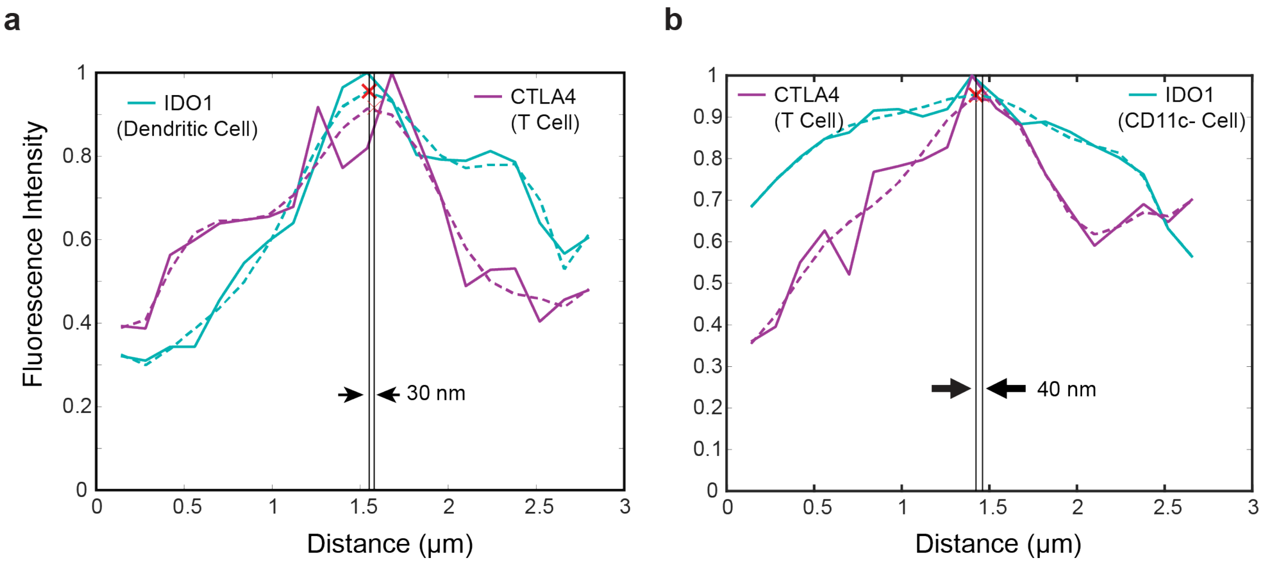


**Supplementary Figure 15: cell—cell membrane interaction analysis.**

**a,** membrane interaction graphs between CD11c$^{+}$ IDO1$^{+}$ cell (cyan, IDO1 channel) and Treg (magenta, CTLA-4 channel). **f,** membrane interaction graphs between CD11c$^{-}$ IDO1$^{+}$ cell (cyan, IDO1 channel) and Treg (magenta, CTLA-4 channel).

#### v-CyCIF workflow decision Tree

**a**

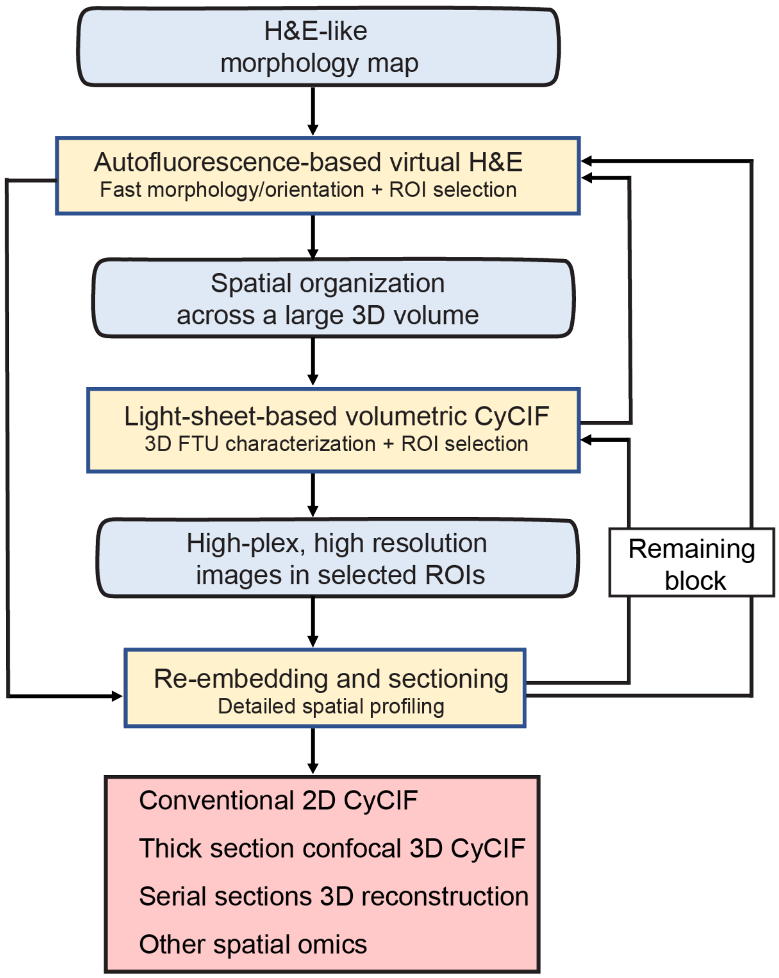


**Supplementary Figure 16: Workflow decision tree for selecting imaging modalities.**

**a,** Decision tree guiding selection from autofluorescence-based virtual H&E, light-sheet–based volumetric CyCIF for large-volume 3D organization, to re-embedding/sectioning for high-plex high-resolution 3D profiling in selected ROIs, with alternatives for downstream spatial assays listed. Virtual H&E can skip v-CyCIF and go straight to re-embedding, and remaining block after sectioning can also go back to v-H&E and v-CyCIF for further analysis.

#### SUPPLEMENTARY INFORMATION

### Volumetric Cyclic Immunofluorescence for 3D Spatial Profiling of Tumour, Stroma, and Immune Structures in Human FFPE Tissue

Alex Y. H. Wong[1,2*], Yi Daniel Lu[1,2*], Ziyuan Zhao[1,2,3], Seo Woo Choi[1,4], Hojeong Park[5], Felix Zhou[6], Zoltan Maliga[1,2], Yvonne N. A. Anang[1,7,8], Soheil R. Talemi[1,2], Shannon Coy[1,7], Gaudenz Danuser[6,9], Sandro Santagata[1,2,3,7], Clarence Yapp[1,2†] & Peter K. Sorger[1,2,3†]

[1] Laboratory of Systems Pharmacology, Harvard Medical School, Boston, MA, USA

[2] Ludwig Centre at Harvard, Harvard Medical School, Boston, MA, USA

[3] Department of Systems Biology, Harvard Medical School, Boston, MA, USA

[4] LifeCanvas Technologies, Cambridge, Boston, MA, USA

[5] Broad Institute of MIT and Harvard, Cambridge, MA, USA

[6] Lyda Hill Department of Bioinformatics, UT Southwestern Medical Center, Dallas, TX, USA

[7] Department of Pathology, Brigham and Women's Hospital, Boston, MA, USA

[8] Current address: Tissue Biomarker Laboratory of the Center for Immuno-Oncology, Department of Medical Oncology, Dana-Farber Cancer Institute, Boston, MA, USA

[9] Current address: Institute for Human Biology, Roche Pharma Research and Early Development (pRED), Roche Innovation Center, Basel, Switzerland

*Authors contributed equally

#### SUPPLEMENTARY NOTE: APPROACHES TO 3D TISSUE PROCESSING, IMAGING, ANALYSIS AND VISUALIZATION

This supplementary note covers aspects of tissue clearing and volumetric imaging that were not fully discussed in the main text due to space limitations.

##### 1. Choice of Tissue clearing methods

We tested and compared a wide range of clearing protocols for use in v- CyCIF. Clearing frequently requires delipidation since lipids play a substantial role in causing tissues to scatter light. Delipidating clearing enables deeper light penetration and a reduction in optical aberrations. However, we found that delipidation was incompatible with antibody staining of many cluster of differentiation (CD) proteins, which are essential for phenotyping immune cells. Using organic solvents such as dichloromethane (DCM) as opposed to detergents did not improve staining. This was in contrast to structural components of cells such as microtubules, cytokeratins and other proteins in the nucleus and cytoplasm; staining of these proteins was generally resistant to delipidation. We therefore decided that a delipidation-free approach would be necessary for efficient staining of immune cells and complexes.

First, we tested OPTIClear, which has been previously used to stain FFPE human brain[1], and belongs to a class of aqueous-delipidiation methods including SWITCH[2] and FASTclear[3]. We also omitted hydrogel embedding since it was not essential for OPTIClear delipidation to work and its elimination reduced tissue expansion and avoided crosslinking of acrylamides to proteins[4]. We tested it on normal colon, which required a few days of robust SDS-based delipidation. Once again, we found that immunofluorescence (IF) staining was limited to stromal markers with poor signal to noise for CD proteins. We also tested a non-toxic solvent-based method based on ethyl-3-phenylprop-2-enoate (Ethyl-cinnamate, ECi). In this case, immune cells could be stained effectively with anti-CD marker antibodies. However, ECi, like many solvent-based methods, requires a series of dehydration steps after staining. This causes tissue shrinkage as well as antibody de-binding and reduction in staining intensity.

In contrast, the delipidation-free v-CyCIF protocol described in the text was found to preserve staining of immune cells and effectively clear specimens up to 1mm thick (RI 1.52). In our approach EasyIndex (Lifecanvas Technologies) achieved delipidation-free reversible refractive index matching following 2-12 hours of incubation in and retained staining for immune, stromal, and tumor markers (as shown in the main figures). To avoid RI mismatch when imaging on the DALISPIM or inverted confocal microscope, it is important to prevent evaporation by adding a layer of immersion oil on top of the layer of EasyIndex on top of the tissue.

#### 1. Imaging configurations evaluated

We compared four imaging platforms for collecting v-CyCIF data: laser scanning confocal microscopy (LSCM), spinning disk confocal microscopy (SDCM), open-top light sheet microscopy (OTLS), and MuVi SPIM Light Sheet Fluorescence Microscopy (LSFM) with respect to multiplexing capacity, sensitivity, spatial resolution, imaging speed, and imaging depth.

Confocal microscopy provided the highest optical resolution and contrast among systems we evaluated. When confocal pinholes are paired with high magnification oil immersion lenses (40×/1.3NA) they enable detailed visualization of small structures with high plexity (as recently described in Yapp et al.[5]) when signal-to-background ratio is adequate. However, these advantages come at the cost of reduced imaging depth due to (i) the limited working distance of immersion objective lenses (typically 200-400 microns) and (ii) excitation of the full tissue volume leading to photobleaching. Between the two variants of confocal microscopy, LSCM is significantly slower than SDCM due to the need to raster scan a laser across the sample; however, it generally provides superior contrast. In practice, LSCM performs optimally across a limited region on thinner sections (30–50 μm) mounted on glass slides. As described in the main text, it is ideally paired with LSFM.

Spinning disk confocal microscopy provides a balance between optical sectioning and acquisition speed by parallelizing excitation through a fixed array of pinholes and collecting emitted photons on an array detector instead of a photomultiplier tube. Compared to point-scanning confocal systems, SDCM achieves substantially faster imaging while maintaining comparable lateral resolution, although axial sectioning is somewhat reduced due to the fixed pinhole size. This configuration improves photon throughput relative to traditional confocal microscopy, but sensitivity remains lower than light sheet–based approaches. The fixed pinhole geometry also limits flexibility in optimizing detection conditions across markers of varying intensity, which can constrain multiplexed imaging performance.

Open-top light sheet microscopes (OTLS) such as MegaSPIM or DALISPIM (Lifecanvas Technologies) or 3Di (Alpenglow Biosciences) are optimized for imaging large, laterally extended cleared tissues, which simplifies handling and preserves sample geometry. This configuration supports high imaging depth when combined with clearing techniques and allows rapid volumetric acquisition due to plane illumination and wide field-of-view detection. This configuration is ideal for highly heterogenous tissue such as human tumours.

When configured with axially swept light-sheet microscopy (ASLM), which utilizes the narrowest portion of the excitation Gaussian beam, and detection objective lenses having shallow depth of focus, the axial resolution can increase by approximately 10-fold relative to a conventional Gaussian beam. These advantages come at the cost of illumination confinement, spatial duty cycle and field of view[6].

The multi-view selective plane illumination microscopy system commercially known as MuVi SPIM (Luxendo, Bruker) similarly achieves excellent imaging depth and high sensitivity by combining light sheet illumination with multi-view acquisition. The ability to illuminate the sample from opposing directions improves excitation light penetration and reduces shadowing artifacts, which is particularly advantageous for thicker or optically heterogeneous specimens. Reconstruction from multiple views further enhances signal recovery and spatial completeness. However, in practice, multiplexed imaging with weakly expressed markers presents a significant limitation, as repeated acquisition across multiple angles increases total light exposure and can lead to photobleaching before all views are collected. While imaging speed is inherently high for single-view acquisition, it is effectively reduced when multiple views are required. Additionally, the imaging geometry is optimized for samples that are narrow and long (such as colon mucosa and skin epidermis) as opposed to flat. Such samples can be mounted in a vertical orientation orthogonally to the detection and excitation objective lenses.

The v-CyCIF approach was consciously designed to work with a wide range of microscopes both to increase the accessibility of the method and because no single data collection modality is ideal for all types of samples. We recommend using the spinning disk confocal microscope for studying the distribution of markers with punctate diffraction limited spots; however, light-sheet is the more obvious choice to find cell-niche interactions or 1 mm thick functional tissue units.

**2. Sample mounting and 3D printed holders**

We designed custom sample holders to support and stabilize tissue specimens during imaging and then fabricated them by 3D printing. The holder consists of a rigid frame design with comb-like structures along the inner edges. These comb features support the sample, which we encase in agarose, ensuring firm adhesion and preventing sample drift. We bond specimens to the combed frame using agarose (or phytagel) and glue. This design maintains optical accessibility and follows the sample through staining, imaging, bleaching, clearing and washes.

The holder is modular and compatible with multiple light-sheet microscopy configurations. For use with the MuVi-SPIM system, the frame can be directly mounted via a clamp. In the MegaSPIM, the holder is positioned within a spacer adapted to a standard 6-well plate format. For the MegaSPIM system, the frame incorporates a click-in design that fits into a spacer with dimensions equivalent to a conventional glass slide. This interchangeable architecture enables consistent sample preparation across different imaging modalities while minimizing handling variability. Additional mounting methods for normal colonic mucosa included using the muscle or submucosa (which was of less scientific interest in our studies) as the only areas of tissue embedded in agarose gel and adhered to the holder. For the DALIPSIM, 1 mm samples were stained in a 48-well plate (staining volume of 500 µl) and imaged in a 12-well plate fitted with a rectangular spacer. The agarose block containing the specimen was trimmed to a rectangle whose corners aligned with the spacer, maintaining a consistent orientation across imaging sessions. One corner (top left) was clipped to break the symmetry of the agarose, providing a fiducial mark that identified the correct side of the cleared specimen at each remount. A 3D printed weight made of PETG was placed on top of the tissue to flatten it and reduce the distance from the glass at the bottom of the well. Design files for the holders and weights are available at (https://github.com/labsyspharm/Volumetric-CyCIF/tree/main/3D%20printed%20sample%20holders).

##### 3. Overview of virtual H&E

A rapidly growing literature on virtual H&E (vH&E) has demonstrated that it is a practical way to generate histology-like images from minimally processed tissue. The first applications involved 2D sections;[7–18] more recently slide-free and volumetric imaging workflows compatible with light-sheet microscopy and other 3D modalities such as microCT have been described[6–16]. Initial descriptions of vH&E transformed autofluorescence images from unstained tissues into H&E-like images computationally using GAN-based models[9], establishing virtual staining as a possible alternative to conventional histochemical processing. Subsequent work broadened the input modalities used for virtual H&E to include label-free autofluorescence lifetime imaging microscopy (FLIM)[25], quantitative phase imaging approaches such as qOBM[26], ultraviolet photoacoustic remote sensing (UV-PARS)[14], and optical coherence–based methods[27], all with the aim of producing diagnostically familiar H&E-like contrast while reducing preparation time and tissue consumption. In parallel, 3D virtual histology approaches using light-sheet microscopy[22–27], X-ray[21–24] and microCT[27] have demonstrated the value of nondestructive volumetric tissue assessment, arbitrary volume reslicing (AVR), and improved sampling of heterogeneous structures.

Inspired by this previous work, we integrated v-CyCIF with vH&E by staining tissues with a nuclear stain to improve the imaging of nuclei (which are normally stained by hematoxylin) and autofluorescence to provide information for eosin estimation. We found that this approach provided rapid insight into 3D tissue morphology without the need for (expensive) antibodies.

This is valuable when implementing v-CyCIF for dissecting complex 3D tissue structures, which are often larger than a single FFPE block, making interpretation dependent on overall specimen orientation within the block (which is usually determined during the process of sample grossing). Generation of a rapid morphological preview (**Fig. 4a-c**) made it possible to orient subsequent rounds of imaging to acquire data from the most informative regions of a multi-layered specimen. In colon, for example, this corresponded to the epithelial layer and the immune-rich lamina propria (the colonic mucosa), not the muscle.

vH&E images were acquired on specimens that had been subjected to antigen retrieval, overnight DAPI incubation, and refractive index matching in EasyIndex. Images were then collected by LSFM prior to sample bleaching: autofluorescence was excited with the 488 nm laser line and detected using a Chroma ET525/50m emission filter; DAPI was excited with the 405 nm laser line and detected using a Chroma ET445-58m emission filter. We then applied a Beer–Lambert-law-based transformation to convert fluorescence into pseudo-absorbance, followed by color mapping to yield blue–purple nuclear staining and pink cytoplasmic/stromal hues consistent with conventional H&E. This simple “physics-based” approach is similar in spirit to prior fluorescence-based false-coloring approaches[31] and contrasts with GAN-based methods that have the potential for hallucination.

This strategy is advantageous because H&E is the diagnostic standard in histopathology, and the extensive body of knowledge for interpreting H&E morphology helps pathologists and tissue biologists familiarize themselves with 3D images[34]. In 2D specimens, H&E and high-plex IF data are highly complementary, linking classical morphology to molecular phenotypes. vH&E extends this complementarity into three-dimensional tissue volumes by providing familiar H&E-like contrast in a format compatible with volumetric imaging. Unlike conventional H&E sections, which are typically performed on ~5 µm sections and evaluated by brightfield microscopy, vH&E can be extended to volumetric 3D imaging of ~1mm stained tissue using LSFM in our case. Moreover, by using histopathology foundation models such as UNI[35], feature-specific segmentation can be performed in 3D vH&E to extract morphological features and interpret the data. For example, segmentation masks generated using foundation models can be converted into surface meshes, allowing crypt topology to be quantified in three dimensions (**Fig. 4d-f**). Overall, we believe that the v-CyCIF implementation of v-H&E is not only a useful addition to antibody-based volumetric tissue imaging, but it is also likely to have application as a stand-alone method.
