## Extended Figures for "Volumetric Cyclic Immunofluorescence for 3D Spatial Profiling of Tumour, Stroma, and Immune Structures in Human FFPE Tissue"

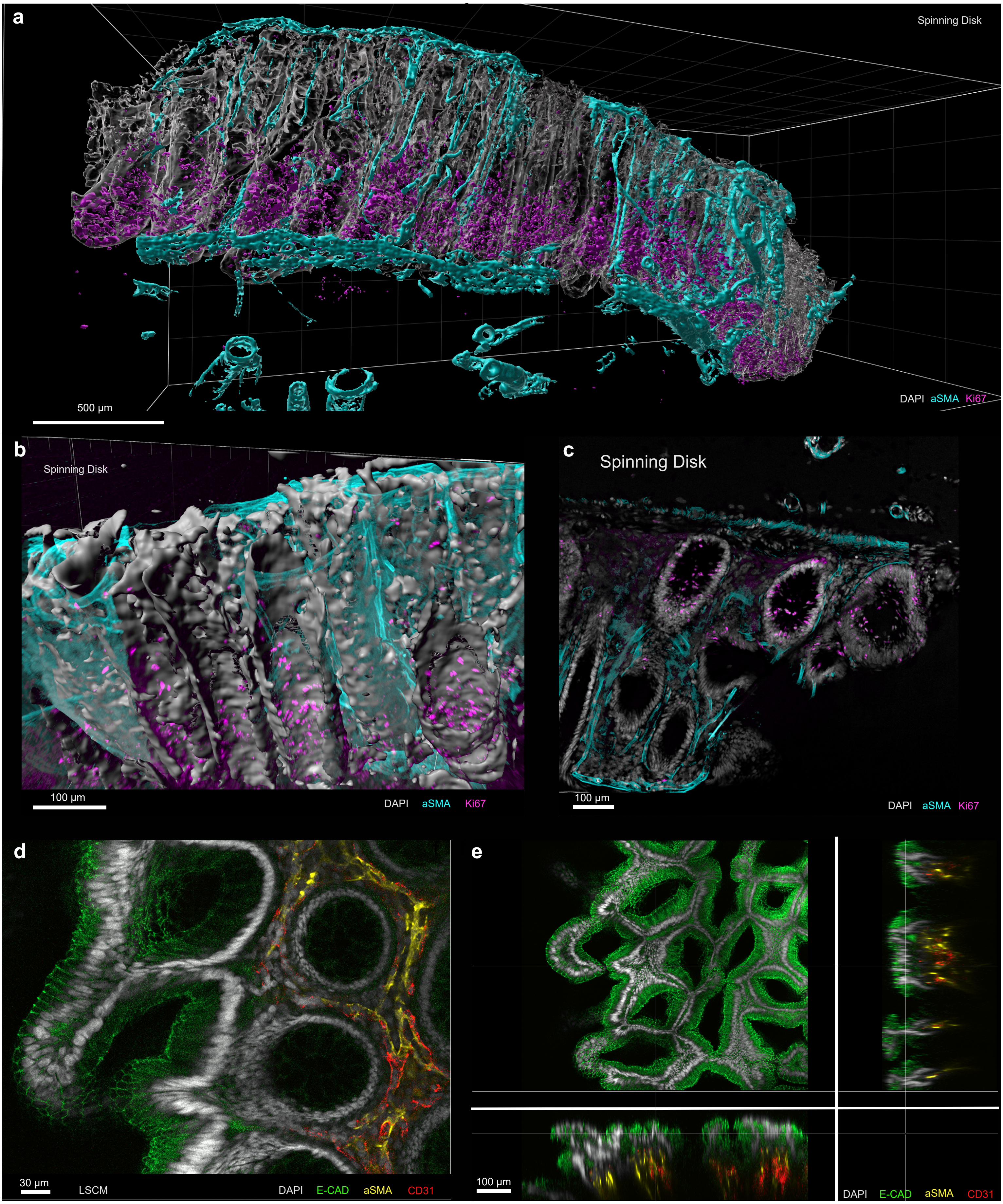
a
Spinning Disk
500 μm
DAPI aSMA Ki67
b
Spinning Disk
100 μm
DAPI aSMA Ki67
c
Spinning Disk
100 μm
DAPI aSMA Ki67
d
30 μm
LSCM
DAPI E-CAD aSMA CD31
e
100 μm
DAPI E-CAD aSMA CD31

**Extended Data Figure 1: v-CyCIF using spinning disk and laser scanning confocal microscopy**

**a,** 3D rendering of a cleared human colon specimen imaged on spinning disk confocal. $\alpha SMA^{+}$ vasculature (cyan) and Ki67 (magenta) rendered as surfaces. Scale bar, 500 µm. **b,** Close-up view of crypts from (**a**) of $Ki67^{+}$ proliferative cells (magenta), nuclei (DAPI, gray) and vasculature (αSMA, cyan) within crypts. Scale bar: 100 µm. **c,** Single optical plane of the colon region. Scale bar: 100 µm. **d,** Maximum intensity projection image of the cleared human colon imaged on laser-scanning confocal microscopy (LSCM), labelled with E-CAD (green), αSMA (yellow) and CD31 (red). Scale bar: 30 µm. **e,** LSCM orthogonal views (XY, XZ and YZ). Scale bar: 100 µm.

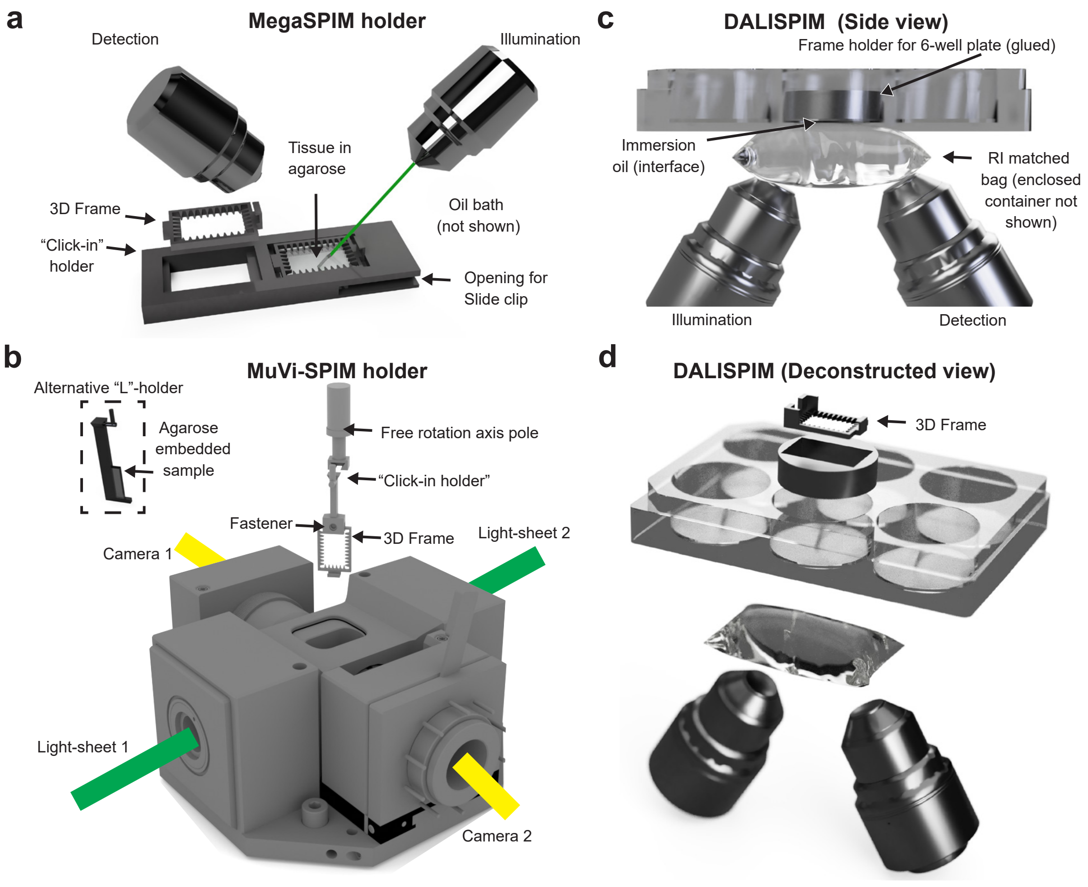

a
MegaSPIM holder
Detection
Illumination
Tissue in agarose
3D Frame
"Click-in" holder
Oil bath (not shown)
Opening for Slide clip
b
MuVi-SPIM holder
Alternative "L"-holder
Agarose embedded sample
Free rotation axis pole
"Click-in holder"
Fastener
3D Frame
Camera 1
Light-sheet 2
Light-sheet 1
Camera 2
c
DALISPIM (Side view)
Frame holder for 6-well plate (glued)
Immersion oil (interface)
RI matched bag (enclosed container not shown)
Illumination
Detection
d
DALISPIM (Deconstructed view)
3D Frame

**Extended Data Figure 2: Refractive-index matching, sample holder design, and fluorescence imaging before and after photobleaching**

**a,** MegaSPIM holder showing an agarose-embedded tissue mounted in a 3D-printed frame and secured in a “click-in” holder for dual-objective illumination and detection. **c,** MuVi-SPIM holder design, including the 3D frame, fastener, and click-in holder used to position the agaroseembedded sample within the dual illumination setup. **d,** DALISPIM side view showing the glued frame holder for a 6-well plate, with the sample immersed in oil and positioned between the illumination and detection objectives inside a sealed imaging setup.

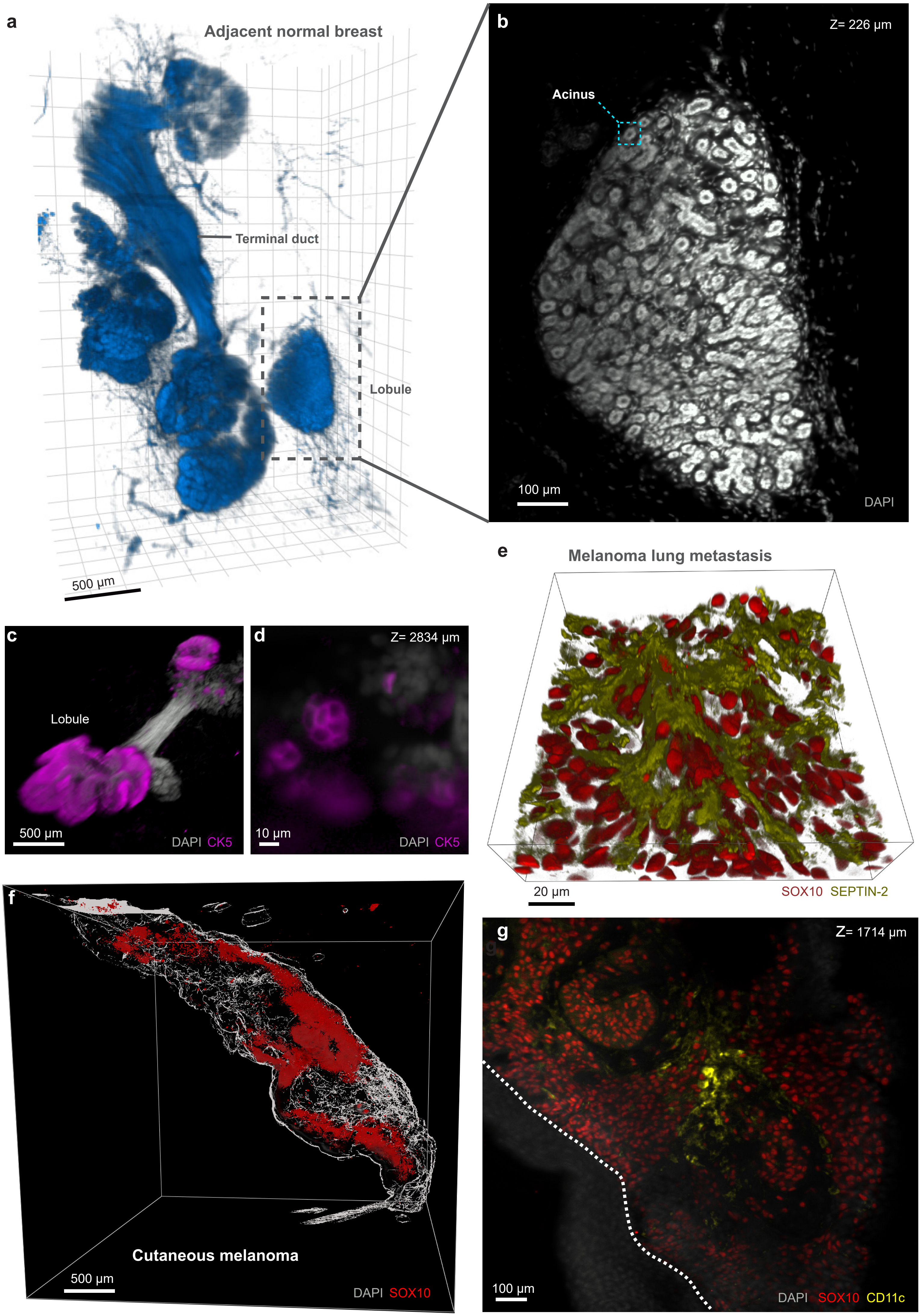

a
Adjacent normal breast
Terminal duct
Lobule
500 µm
b
Z= 226 µm
Acinus
100 µm
DAPI
c
Lobule
500 µm
DAPI CK5
d
Z= 2834 µm
10 µm
DAPI CK5
e
Melanoma lung metastasis
20 µm
SOX10 SEPTIN-2
f
Cutaneous melanoma
500 µm
DAPI SOX10
g
Z= 1714 µm
100 µm
DAPI SOX10 CD11c

**Extended Data Figure 3: Additional v-CyCIF examples demonstrating applicability across organs and disease contexts.**

**a,** 3D volumetric view of adjacent normal breast tissue highlighting ductal and lobular architecture (terminal duct and lobule annotated). Scale bars: 500 μm. **b,** DAPI signal of acinar region (dashed box in **a**) at the z depth of 226 μm. Scale bars: 100 μm. **c,** 3D volumetric view of breast locular structures (DAPI, gray; CK5, magenta). Scale bars: 500 μm. **d,** Single optical plane of (**c**) at the z depth of 2834 μm. Scale bars: 10 μm **e,** 3D volumetric view of melanoma lung metastasis showing tumor marker (SOX10, red) and associated structure (SEPTIN-2, dark yellow). Scale bars: 20 μm. **f,** 3D volumetric view of cutaneous melanoma with tumor marker (SOX10, red) over nuclei (DAPI, gray). Scale bars: 500 μm. **g,** A ROI showing immune-associated signal (CD11c, yellow) in the tumor at the z depth of 1714 μm. Scale bars: 100 μm.

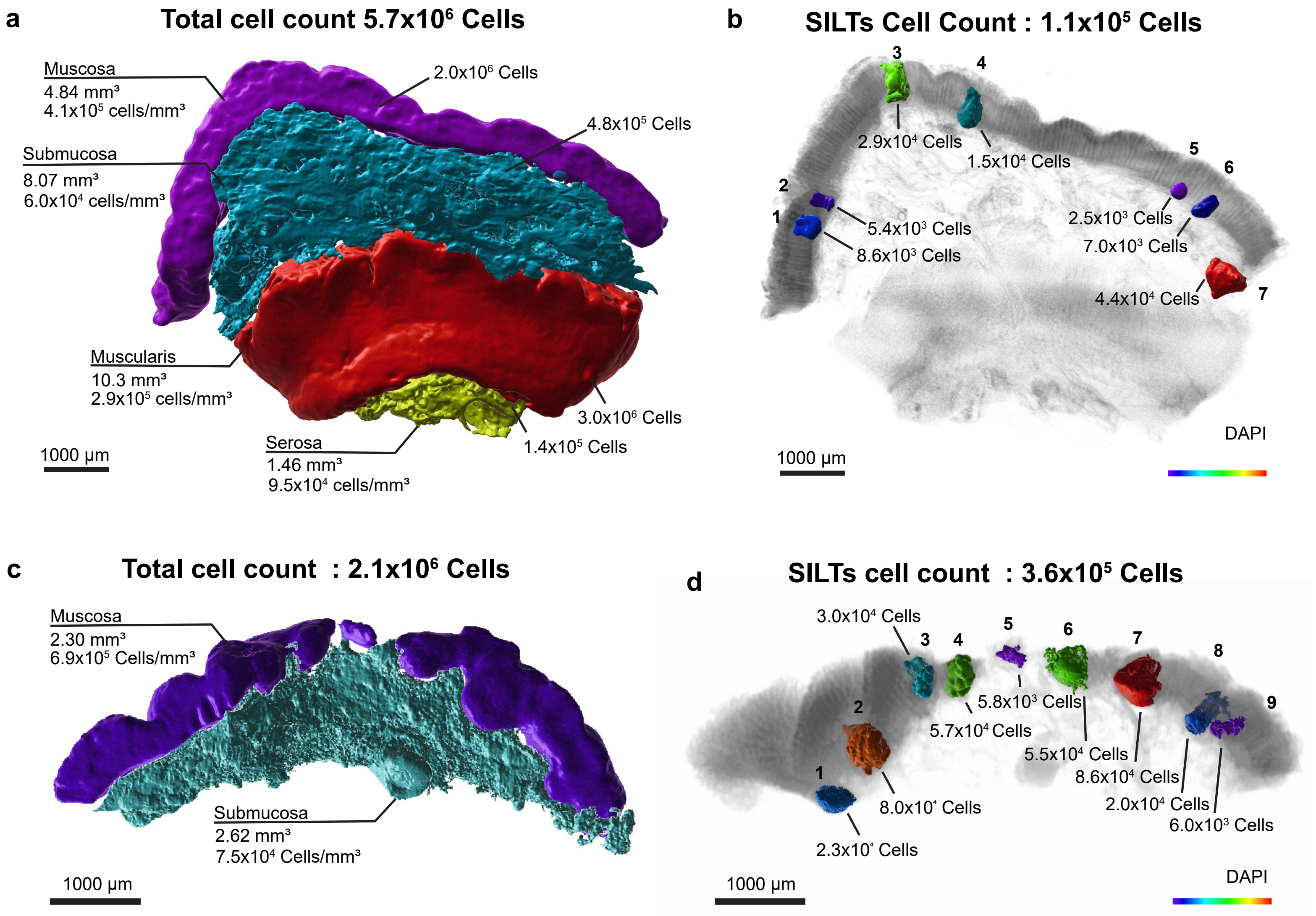

a
Total cell count 5.7x10$^{6}$ Cells
Muscosa
4.84 mm$^{3}$
4.1x10$^{5}$ cells/mm$^{3}$
2.0x10$^{6}$ Cells
4.8x10$^{5}$ Cells
Submucosa
8.07 mm$^{3}$
6.0x10$^{4}$ cells/mm$^{3}$
Muscularis
10.3 mm$^{3}$
2.9x10$^{5}$ cells/mm$^{3}$
3.0x10$^{6}$ Cells
1.4x10$^{5}$ Cells
Serosa
1.46 mm$^{3}$
9.5x10$^{4}$ cells/mm$^{3}$
1000 µm
b
SILTs Cell Count : 1.1x10$^{5}$ Cells
3
4
2.9x10$^{4}$ Cells
1.5x10$^{4}$ Cells
5
6
2
1
5.4x10$^{3}$ Cells
8.6x10$^{3}$ Cells
2.5x10$^{3}$ Cells
7.0x10$^{3}$ Cells
4.4x10$^{4}$ Cells
7
DAPI
1000 µm
c
Total cell count : 2.1x10$^{6}$ Cells
Muscosa
2.30 mm$^{3}$
6.9x10$^{5}$ Cells/mm$^{3}$
Submucosa
2.62 mm$^{3}$
7.5x10$^{4}$ Cells/mm$^{3}$
1000 µm
d
SILTs cell count : 3.6x10$^{5}$ Cells
3.0x10$^{4}$ Cells
3
4
5
6
7
8
9
5.8x10$^{3}$ Cells
5.7x10$^{4}$ Cells
5.5x10$^{4}$ Cells
8.6x10$^{4}$ Cells
2.0x10$^{4}$ Cells
6.0x10$^{3}$ Cells
2
1
8.0x10$^{4}$ Cells
2.3x10$^{4}$ Cells
DAPI
1000 µm

**Extended Data Figure 4: Cell count, region density and SILT quantification across the colon.**

**a,** 3D segmentation of an intact colon specimen showing major anatomical layers (mucosa, submucosa, muscularis, and serosa; colour-coded) with total cell counts per layer and corresponding volumes and estimated cell densities (cells $mm^{-3}$) as indicated. **b,** Identification and quantification of SILTs within the same specimen. Individual SILTs are pseudo-colored and numbered; labels indicate the cell count for each SILT. Background grayscale signal shows DAPI to provide tissue context. **c,** 3D segmentation and layer-resolved total cell count for an additional colon segment (mucosa and submucosa shown), with estimated volume and cell densities indicated. **d,** SILT detection and cell-count quantification for the specimen shown in (**c**), pseudo-colored SILTs over a DAPI background. Scale bars: 1000 µm (**a–d**).

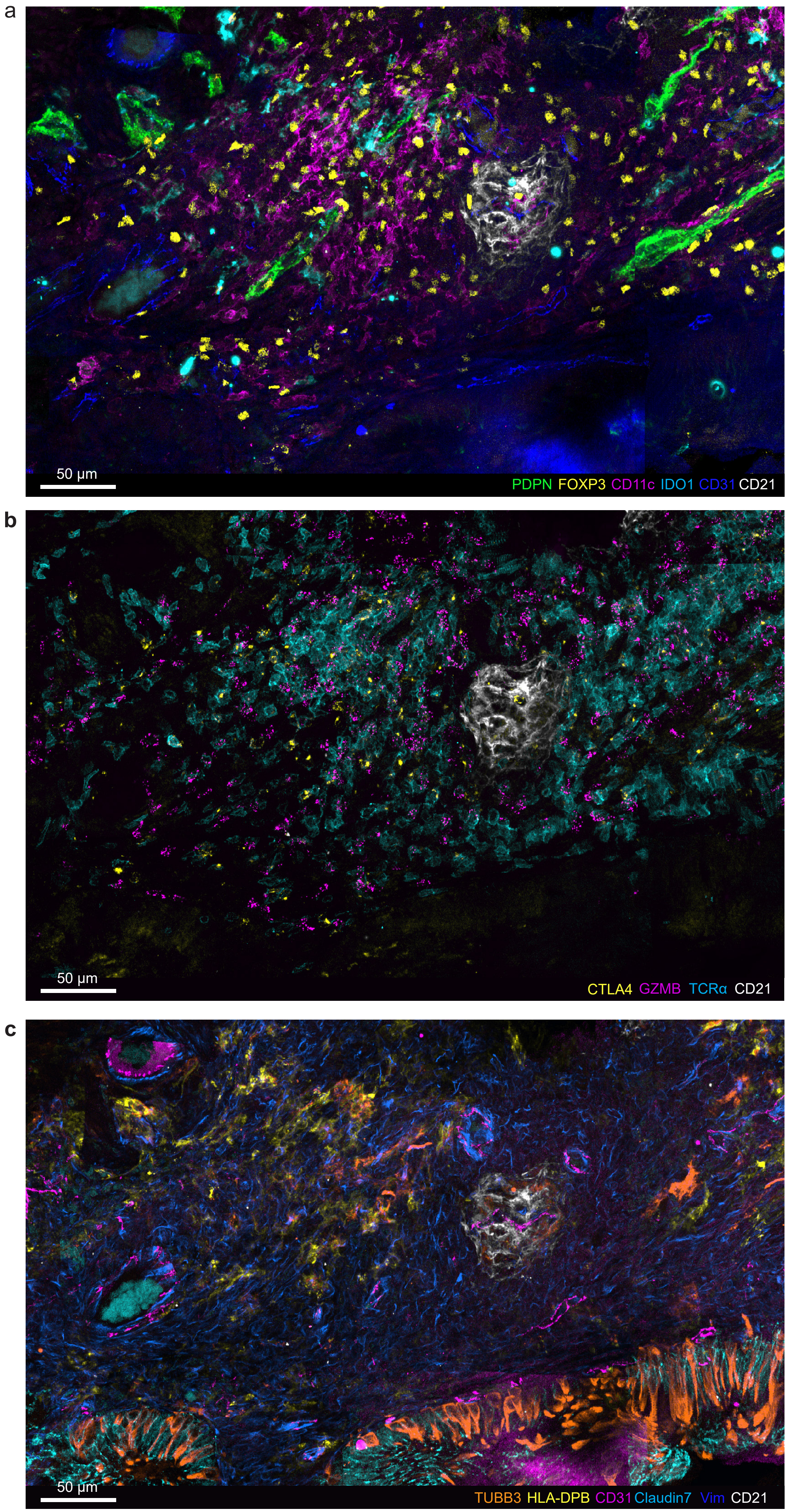
a
50 µm
PDPN FOXP3 CD11c IDO1 CD31 CD21
b
50 µm
CTLA4 GZMB TCRα CD21
c
50 µm
TUBB3 HLA-DPB CD31 Claudin7 Vim CD21

**Extended Data Figure 5: Re-embedded 3D CyCIF of germinal center in tertiary lymphoid structures (TLS)**

**a,** Re-embedded 3D CyCIF image showing spatial organization of a germinal center defined by a $CD21^+$ follicular dendritic cell (FDC) network (white) surrounded by $PDPN^+$ lymphatic vessels (green) and CD31+ vasculature (dark blue), with TLS-associated immune populations including $FOXP3^+$ regulatory T cells (yellow), $CD11c^+$ dendritic cells (magenta), and $IDO1^+$ immunoregulatory myeloid cells (cyan). Scale bar: 50 μm. **b,** Complementary marker set highlighting the immune composition of TLS, showing $TCR\alpha^+$ T cells (cyan), CTLA-$4^+$ T cells (yellow), and $GZMB^+$ cytotoxic T cells (magenta) arrayed around the $CD21^+$ FDC network (white). Scale bar: 50 μm. **c,** Tissue-context view of the same region showing $CD21^+$ FDC organization (white) relative to local tumour microenvironment, including HLA-$DPB^+$ antigen presenting cells (yellow), $CD31^+$ vasculature (magenta), Vimentin $(Vim)^+$ mesenchymal/stromal area (dark blue), $Claudin7^+$ tumour epithelial structures (Cyan), and $TUBB3^+$ nerve fibers (orange). Scale bar: 50 μm.

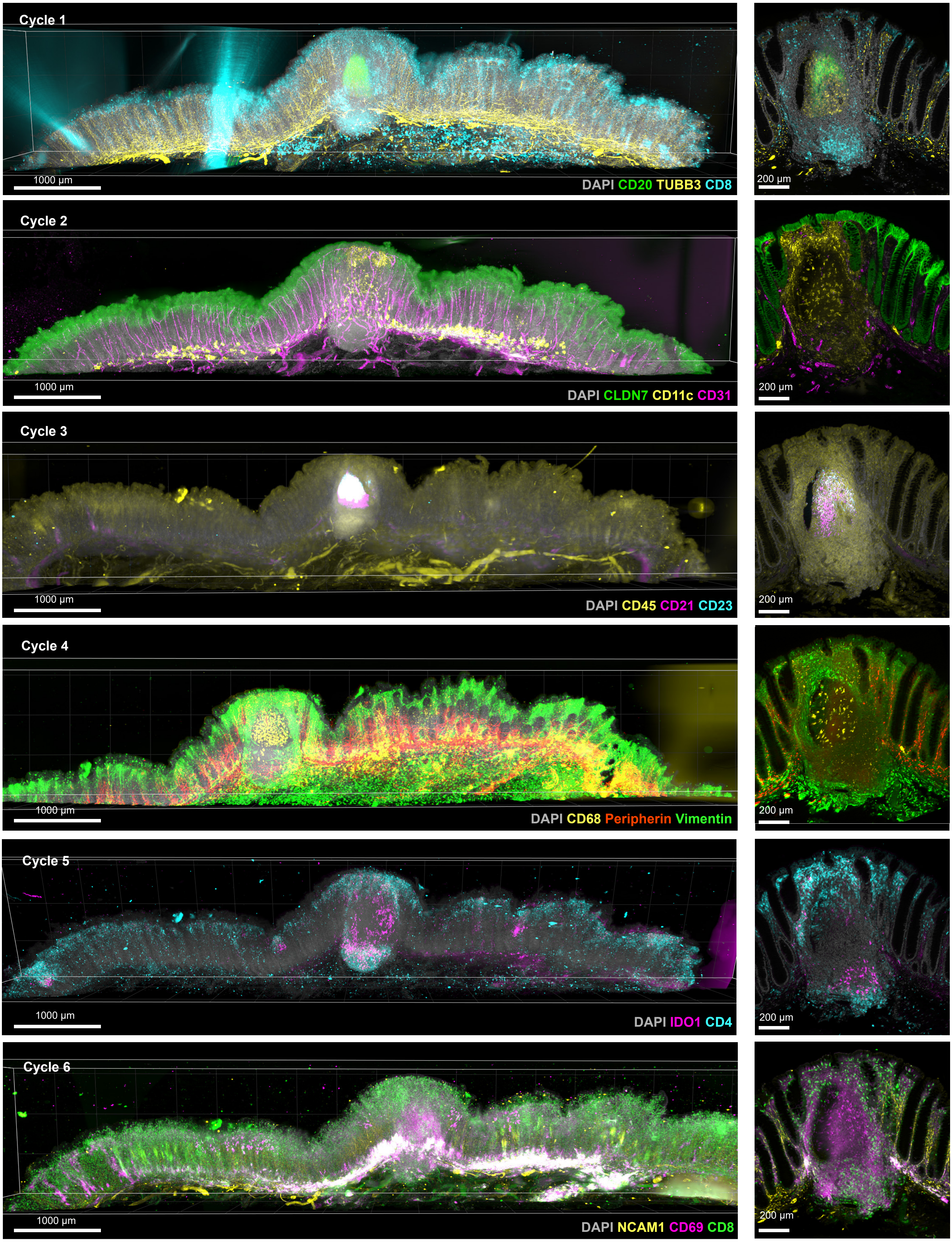

Cycle 1
1000 µm
DAPI CD20 TUBB3 CD8
200 µm
Cycle 2
1000 µm
DAPI CLDN7 CD11c CD31
200 µm
Cycle 3
1000 µm
DAPI CD45 CD21 CD23
200 µm
Cycle 4
1000 µm
DAPI CD68 Peripherin Vimentin
200 µm
Cycle 5
1000 µm
DAPI IDO1 CD4
200 µm
Cycle 6
1000 µm
DAPI NCAM1 CD69 CD8
200 µm

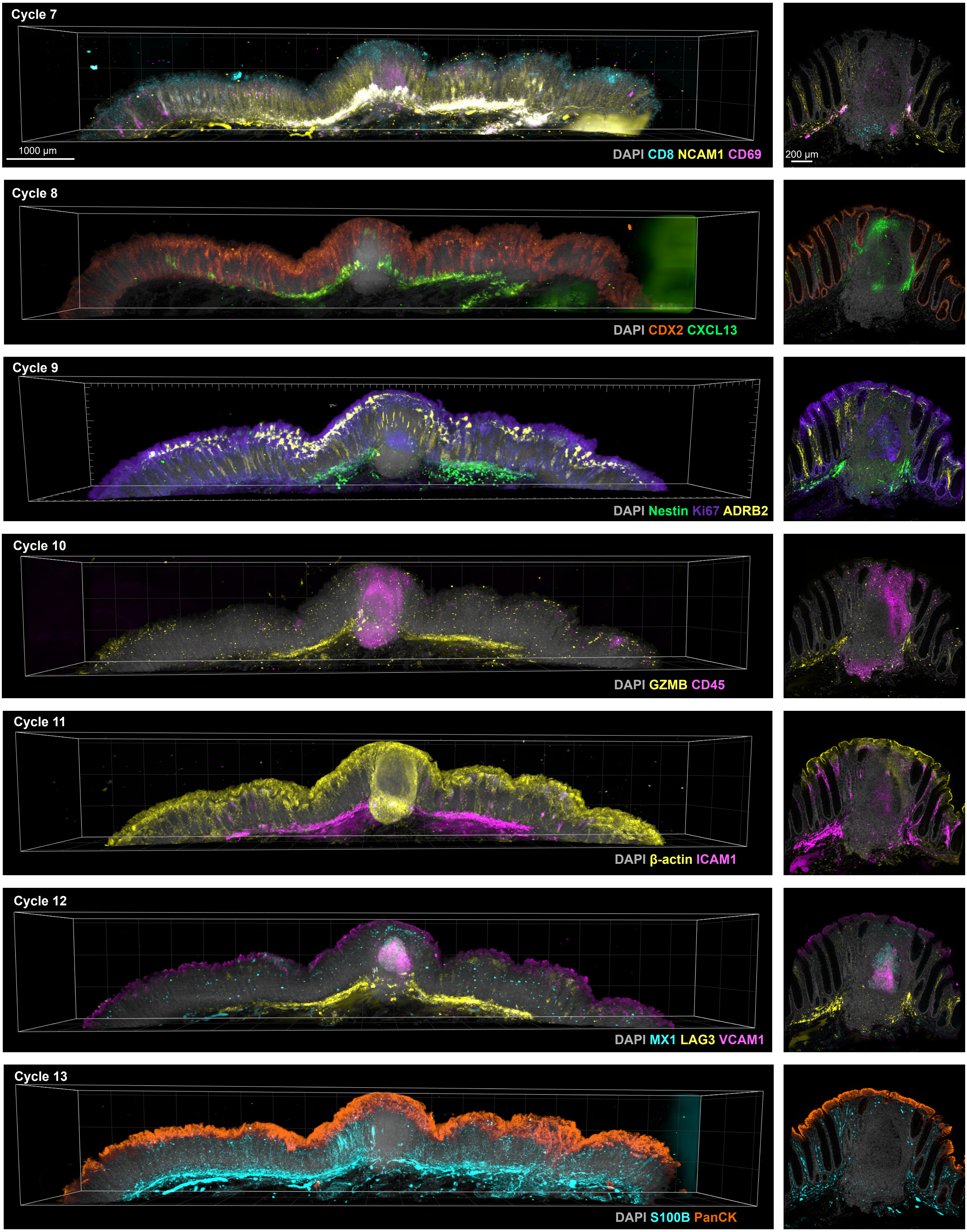

Cycle 7
1000 µm
DAPI CD8 NCAM1 CD69
200 µm
Cycle 8
DAPI CDX2 CXCL13
Cycle 9
DAPI Nestin Ki67 ADRB2
Cycle 10
DAPI GZMB CD45
Cycle 11
DAPI β-actin ICAM1
Cycle 12
DAPI MX1 LAG3 VCAM1
Cycle 13
DAPI S100B PanCK

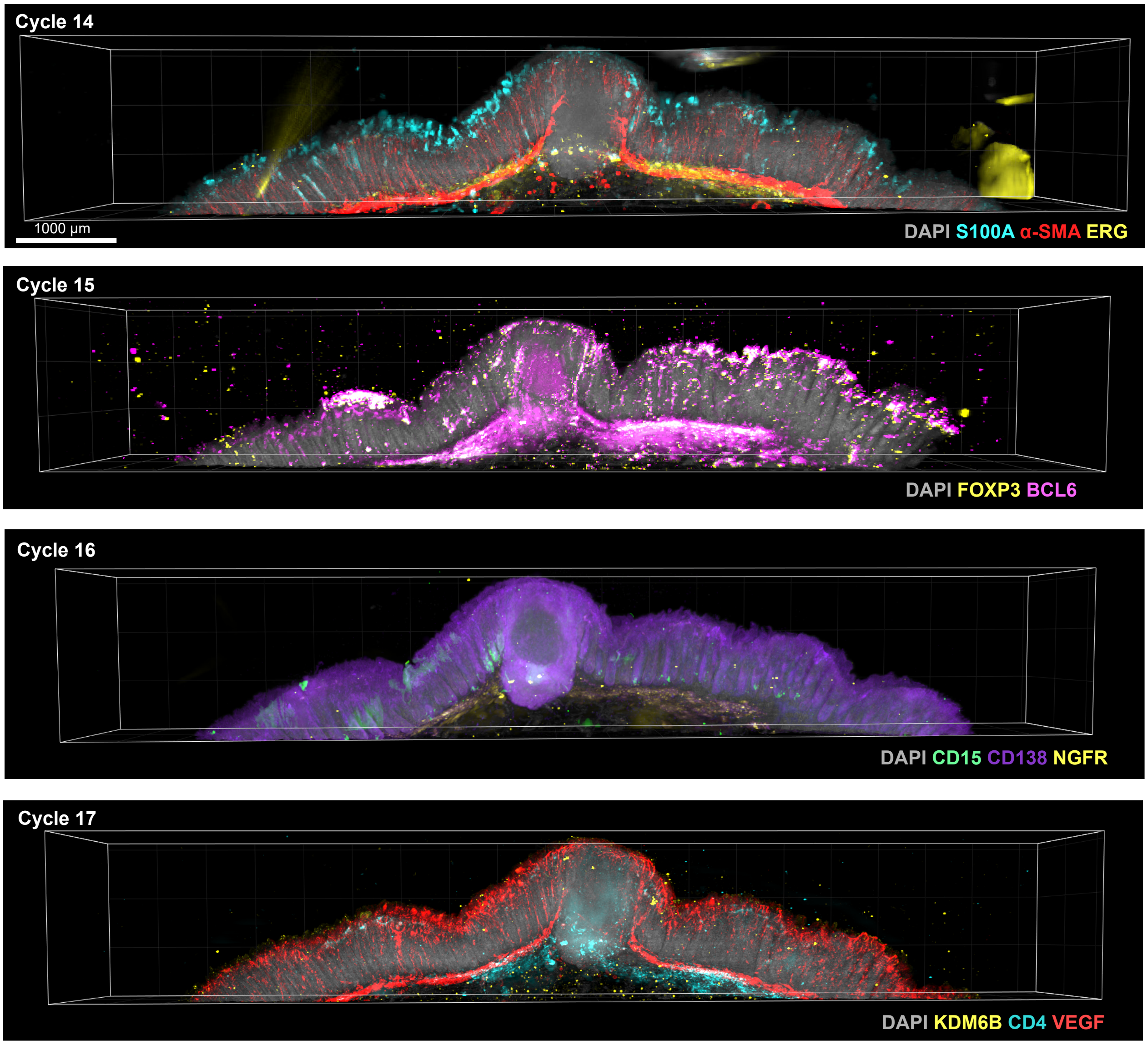
Cycle 14
1000 µm
DAPI S100A α-SMA ERG
Cycle 15
DAPI FOXP3 BCL6
Cycle 16
DAPI CD15 CD138 NGFR
Cycle 17
DAPI KDM6B CD4 VEGF

**Extended Data Figure 6: v-CyCIF of solitary intestinal lymphoid tissue across cycles**

Representative volumetric renderings (left; scale bars: 1000 μm) and corresponding zoomed-in cross-sectional views (right; scale bars: 200 μm) of intact normal colon tissue imaged on DALISPIM light sheet microscope over 17 iterative cycles, with three to four markers visualized per cycle. Cycles 1–17 are shown; markers for each cycle are indicated in the panel labels.

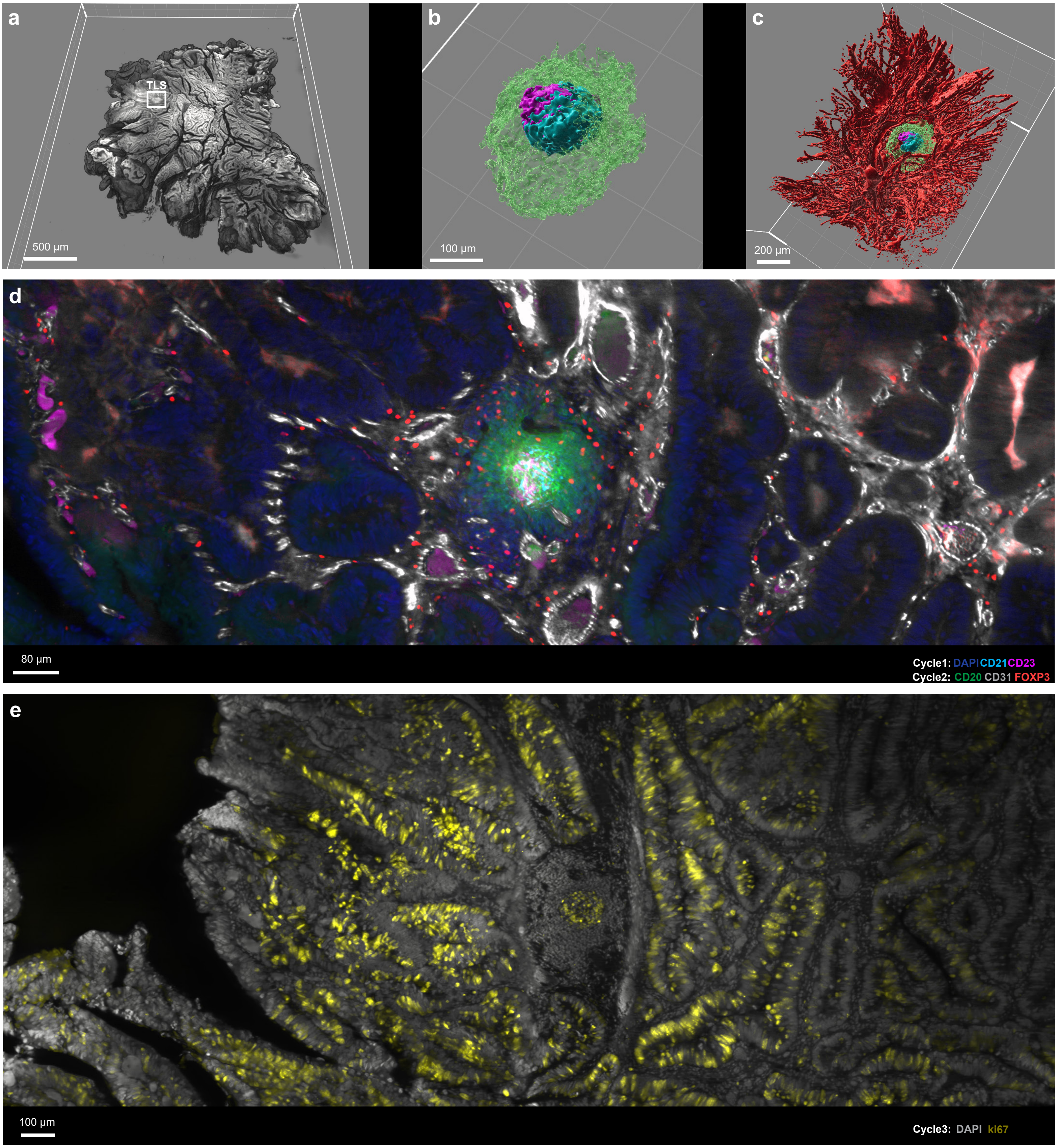

a
TLS
500 µm
b
100 µm
c
200 µm
d
80 µm
Cycle1: DAPI CD21 CD23
Cycle2: CD20 CD31 FOXP3
e
100 µm
Cycle3: DAPI ki67

**Extended Data Figure 7: Isolated intratumoral TLS within a 1 mm adjacent CRC tissue slab**

**a,** Overview of 1 mm CRC tissue slab stained with DAPI. Region of interest (white) shows the location of the TLS within the tumour. Scale bar: 500 µm. **b,** 3D rendering of the FDC network with CD21 (Cyan), CD23 (Magenta) and CD20 (green) surfaces. Scale bar: 100 µm. **c,** Surface rendering of $CD31^+$ blood vessels surrounding the TLS. Scale bar: 200 µm. **d,** Registered images containing CD21 and CD23 in cycle 1, and CD20, FOXP3 and CD31 in cycle 2. Scale bar: 80 µm. **e,** Proliferative $ki67^+$ germinal centre and glandular tumour. Scale bar: 100 µm.

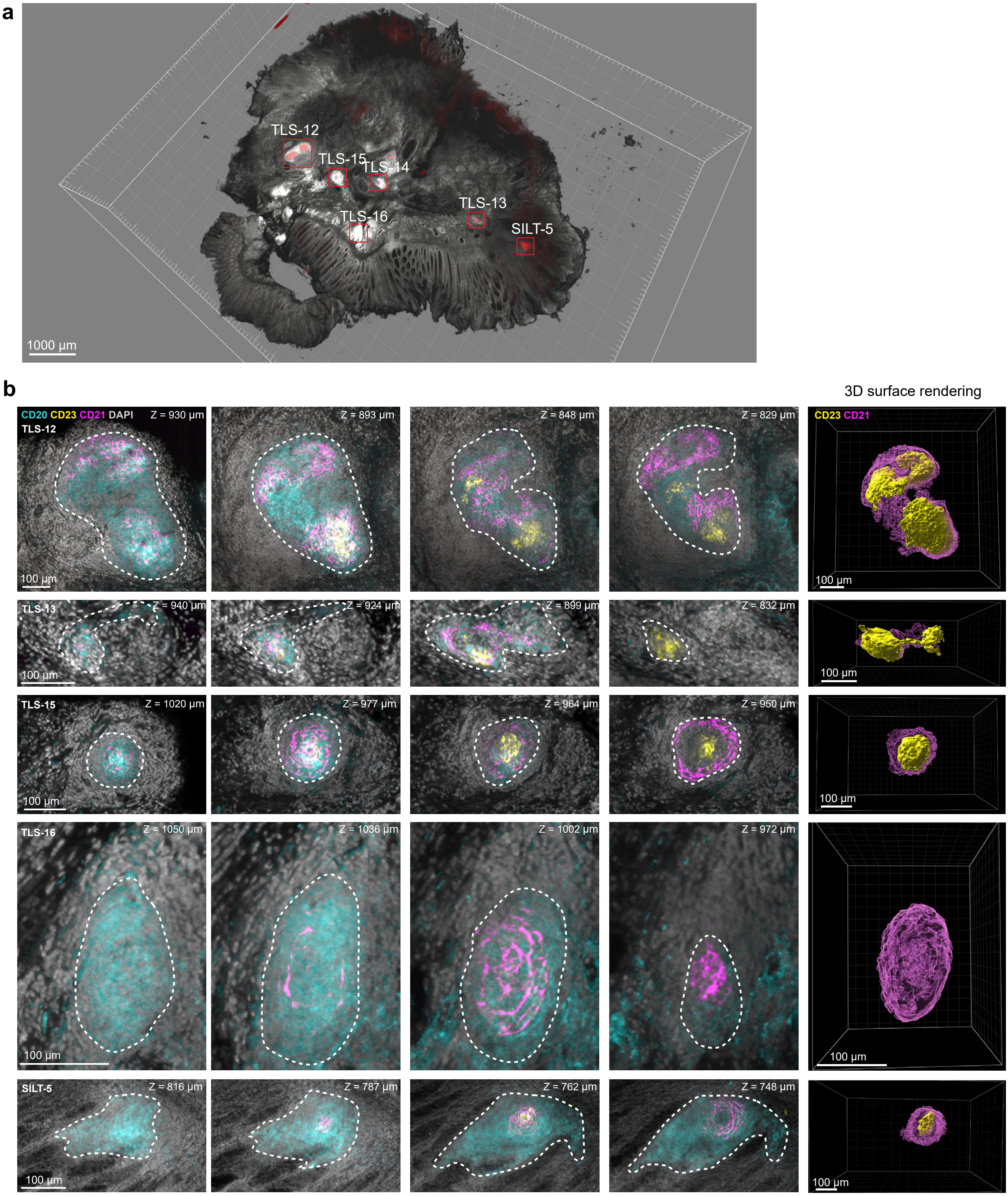
a
TLS-12
TLS-15
TLS-14
TLS-13
TLS-16
SILT-5
1000 µm
b
3D surface rendering
CD20 CD23 CD21 DAPI
TLS-12
Z = 930 µm
Z = 893 µm
Z = 848 µm
Z = 829 µm
CD23 CD21
100 µm
TLS-13
Z = 940 µm
Z = 924 µm
Z = 899 µm
Z = 832 µm
TLS-15
Z = 1020 µm
Z = 977 µm
Z = 964 µm
Z = 950 µm
TLS-16
Z = 1050 µm
Z = 1036 µm
Z = 1002 µm
Z = 972 µm
SILT-5
Z = 816 µm
Z = 787 µm
Z = 762 µm
Z = 748 µm

**Extended Data Figure 8: 3D visualization of tertiary lymphoid structures and solitary intestinal lymphoid tissue within colorectal cancer tissue.**

**a,** 3D volume rendering of the cleared colorectal cancer specimen (DAPI, gray) with red boxes marking the locations of individual lymphoid aggregates (5 TLS and 1 SILT) identified within the tissue volume. Scale bar: 1000 µm. **b,** Serial optical sections at different z-depth (left to right) through five representative structures (TLS-12, TLS-13, TLS-15, TLS-16, and SILT-5; dashed outlines), stained for CD20 (cyan, B cells), CD23 (yellow, follicular dendritic cells), CD21 (magenta, follicular dendritic cells), and DAPI (gray, nuclei). Scale bars: 100 µm. Right column: 3D surface renderings of the $CD23^+$ (yellow) and $CD21^+$ (magenta) FDC networks for each structure. Scale bars: 100 µm.
